# Pseudouridine Residues as Substrates for Serum Ribonucleases

**DOI:** 10.1101/2025.06.18.660232

**Authors:** Clair S. Gutierrez, Bjarne Silkenath, Volga Kojasoy, Jaroslaw A. Pich, Daniel C. Lim, Ronald T. Raines

## Abstract

In clinical uses, RNA must maintain its integrity in serum that contains ribonucleases (RNases), especially RNase 1, which is a human homolog of RNase A. These omnipresent enzymes catalyze the cleavage of the P–O^5′′^ bond on the 3′ side of pyrimidine residues. Pseudouridine (Ψ) is the most abundant modified nucleoside in natural RNA. The substitution of uridine (U) with Ψ or *N*^1^-methylpseudouridine (m^1^Ψ) reduces the immunogenicity of mRNA and increases ribosomal translation, and these modified nucleosides are key components of RNA-based vaccines. Here, we assessed the ability of RNase A and RNase 1 to catalyze the cleavage of the P–O^5′′^ bond on the 3′ side of Ψ and m^1^Ψ. We find that these enzymes catalyze the cleavage of UpA up to 10-fold more efficiently than the cleavage of ΨpA or m^1^ΨpA. X-ray crystallography of enzyme-bound nucleoside 2′,3′-cyclic vanadate complexes and molecular dynamics simulations of enzyme·dinucleotide complexes show that U, Ψ, and m^1^Ψ bind to RNase A and RNase 1 in a similar manner. Quantum chemistry calculations suggested that the higher reactivity of UpA is intrinsic, arising from an inductive effect that decreases the p*K*_a_ of the 2′-hydroxy group of U and enhances its nucleophilicity toward the P–O^5′′^ bond. Experimentally, we found that UpA does indeed undergo spontaneous hydrolysis faster than does m^1^ΨpA. Our findings inform the continuing development of RNA-based vaccines and therapeutic agents.

## INTRODUCTION

Katalin Karikó and Drew Weissman were awarded the 2023 Nobel Prize in Physiology or Medicine for their discovery of nucleobase modifications that led to effective mRNA vaccines against COVID-19 (Krammer and Palese 2024). These modifications overcome an immune response elicited by unmodified mRNA (Weissman et al. 2000; Sahin et al. 2014; Damase et al. 2021) that could have evolved as a defense mechanism against viral RNA (Chen et al. 2021). Endosomal Toll-like receptors are responsible for this response (Karikó et al. 2004; Karikó et al. 2005), and Karikó and Weissman discovered that replacing uridine (U) with pseudouridine (Ψ) enabled synthetic mRNAs to evade the receptors while enhancing ribosomal translation (Karikó et al. 2005; Karikó et al. 2008; Anderson et al. 2010; Karikó et al. 2011; Morais et al. 2021).

U and Ψ residues form similar hydrogen bonds with adenosine residues in canonical Watson–Crick–Franklin base pairs (Fig. 1). In addition, Ψ can donate an additional hydrogen bond in the major groove, enabling enhanced local RNA stacking, which is amplified by neighboring nucleosides (Charette and Gray 2000; Hudson et al. 2013; Kierzek et al. 2013; Spenkuch et al. 2014). The methylation of Ψ at *N*^1^ to provide *N*^1^-methylpseudouridine (m^1^Ψ) obviates the additional hydrogen bond but elicits more protein production than Ψ and further diminishes the immunogenicity (Andries et al. 2015; Svitkin et al. 2017; Parr et al. 2020). Ultimately, m^1^Ψ was used in most of the 14 billion administered doses of the COVID-19 vaccine (Nance and Meier 2021; Vogel et al. 2021; Demongeot and Fougère 2022).

**FIGURE 1.**
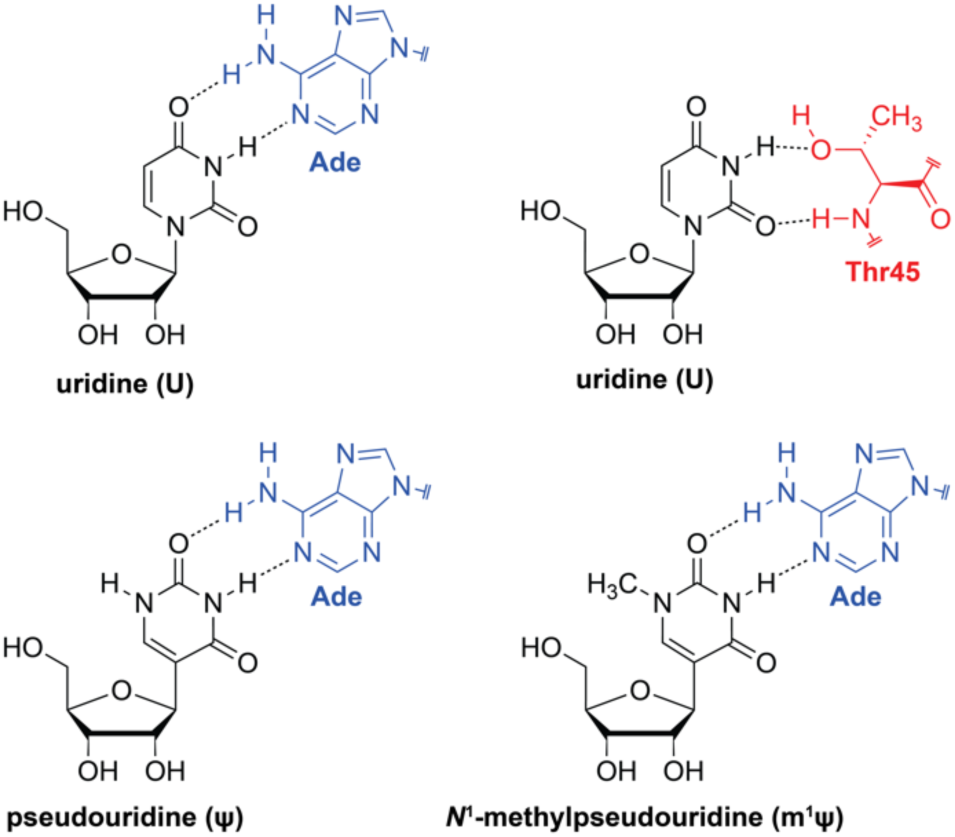
Structure of uridine (U), pseudouridine (Ψ), and *N*^1^-methylpseudouridine (m^1^Ψ), their base-pairing with adenine (Ade), and the interaction of U with Thr45 of ptRNases.

Ψ, which is the *C*^5^-glycoside isomer of U, was discovered in 1951 as the first post-transcriptional modification in RNA (Cohn and Volkin 1951). Today, Ψ is known to be the most abundant modified nucleoside and is found in RNA from all domains of life (Spenkuch et al. 2014). Ψ is enriched in the coding sequence and 3′-untranslated regions of mRNAs (Carlile et al. 2014; Schwartz et al. 2014; Karijolich et al. 2015; Li et al. 2015; Cerneckis et al. 2022). In nearly all tRNAs, Ψ is found in the TΨC stem-loop (and elsewhere) and can stabilize tRNA conformation (Gray and Charette 2000; Motorin and Helm 2010; Guzzi et al. 2018). Ψ accounts for 1.4% of all nucleosides in human rRNA and modulates its conformational dynamics (Jiang et al. 2015; Penzo and Montanaro 2018; Cerneckis et al. 2022). m^1^Ψ is also a natural nucleoside but is much less common than Ψ (Wurm et al. 2012).

Surprisingly little is known about Ψ or m^1^Ψ as substrates for ribonucleases. Ψ does impede the ability of human RNase L (Anderson et al. 2011) and bacterial RNase E (Islam et al. 2021) to degrade RNA. Yet, no analyses have been performed with pancreatic-type ribonucleases (ptRNases). These secretory enzymes are by far the most abundant and active ribonucleases in humans and other vertebrates (Green and Sambrook 2019; Sun et al. 2022) and are of utmost concern for RNA-based vaccine development and delivery (Wang et al. 2021). In ptRNases, a conserved residue, Thr45 (Fig. 1), forms hydrogen bonds with pyrimidine residues in an RNA substrate and precludes the binding of purine residues (delCardayré and Raines 1994; delCardayré and Raines 1995; Kelemen et al. 2000).

Here, we report on Ψ and m^1^Ψ as substrates for ptRNases. We focus our analyses on two homologs: (1) bovine pancreatic ribonuclease (RNase A), which served as a model protein for seminal studies in biological chemistry during the twentieth century (D’Alessio and Riordan 1997; Raines 1998; Marshall et al. 2008), and (2) human ribonuclease 1 (RNase 1), which is the most abundant ribonuclease in human serum and has especially high catalytic activity (Lomax et al. 2017; Garnett and Raines 2022). We investigate the kinetics of the enzyme-catalyzed and uncatalyzed cleavage of synthetic dinucleotide substrates: uridylyl(3′→5′)adenosine (UpA), pseudouridylyl(3′→5′)adenosine (ΨpA), and *N*^1^-methylpseudouridylyl(3′→5′)adenosine (m^1^ΨpA). To correlate structure and function, we deploy X-ray crystallography to obtain high-resolution structures of RNase A complexed with the 2′,3′-cyclic vanadyl diesters of U, Ψ, and m^1^Ψ. To gain structural insight into substrate binding, we carried out molecular dynamics simulations of RNase 1 complexes with UpA, ΨpA, and m^1^ΨpA. Finally, we examined the instrinsic stability of UpA, ΨpA, and m^1^ΨpA. The ensuing data provide guidance for the development of RNA-based vaccines and therapeutic agents along with new insight about the most common post-transcriptional modification.

## RESULTS

### Synthesis of dinucleotide substrates

ptRNases catalyze the cleavage of the P–O^5′′^ bond in RNA between a pyrimidine residue and (preferentially) a purine residue (Fontecilla-Camps et al. 1994). UpA has been the most often used dinucleotide substrate, and its cleavage forms uridine 2′,3′-cyclic phosphate and adenosine (Cuchillo et al. 1993; Thompson et al. 1994). An ensuing change in UV absorption enables the facile determination of steady-state kinetic parameters (Witzel and Barnard 1962). Reasoning that ΨpA and m^1^ΨpA would also be amenable to this assay, we synthesized UpA, ΨpA, and m^1^ΨpA from their component nucleosides by using the phosphoramidite method (Caruthers 2011) (Scheme 1).

Briefly, AgNO_3_-mediated silylation allowed for the synthesis of the three 5′- and 2′-silyl- protected nucleosides **1a**–**c** (Stowell et al. 1995). The reaction of the 3′-hydroxy group with 2-cyanoethyl tetraisopropylphosphorodiamidite and tetrazole gave phosphoramidites **2a**–**c**. 2′,3′-Silyl-protected adenosine **4** was accessed by the protection of the three hydroxy groups of adenosine with TBS to provide **3**, followed by selective 5′ deprotection in aqueous acetic acid (Ogilvie et al. 1978).

Activating phosphoramidites **2a**–**c** with tetrazole and coupling with **4**, followed by oxidation with iodine, gave protected dinucleotides **5a**–**c**. β-Elimination of the cyanoethyl groups with ammonium hydroxide followed by silylether deprotection with tetrabutylammonium fluoride and purification by anion-exchange chromatography gave UpA, ΨpA, and m^1^ΨpA as triethylammonium salts.

**SCHEME 1.**
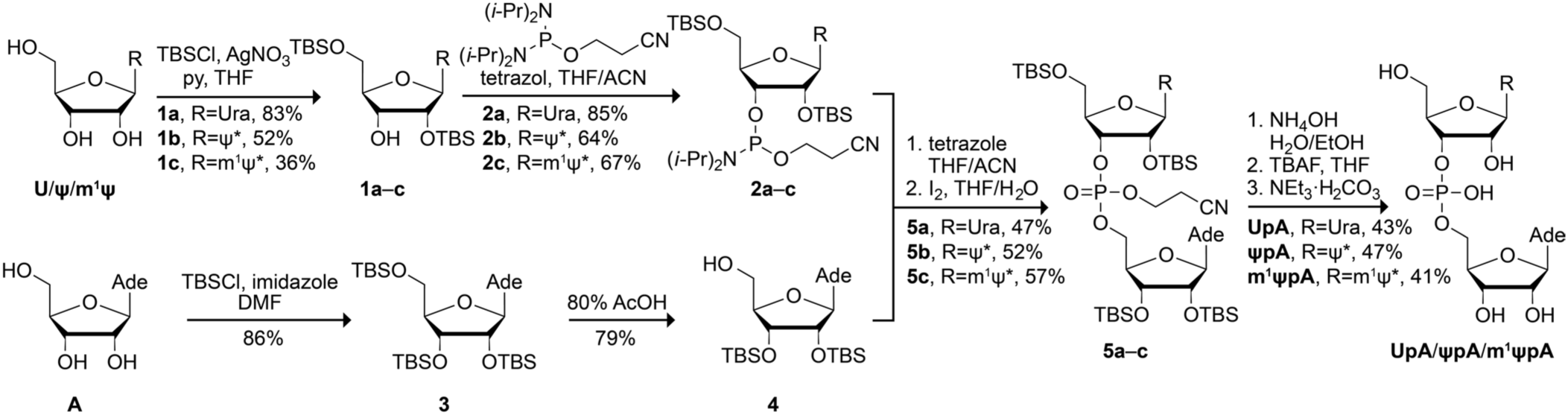
Synthetic route to UpA, ΨpA, and m^1^ΨpA. Ψ* = a pseudouracil nucleobase.

### Heterologous Production of RNase 1

Human RNase 1 was produced in *Escherichia coli* by recombinant DNA technology as described previously (Ressler et al. 2019). Both purified RNase 1 (Fig. S1) and RNase A (which was obtained from a commercial vendor) catalyzed the cleavage of a model substrate, FAM– dArUdAdA–6-TAMRA (Fig. S2), with *k*_cat_/*K*_M_ values similar to those in the literature (Table S1).

### UpA, ΨpA, and m^1^ΨpA as substrates for RNase A and RNase 1

Catalysis of UpA, ΨpA, and m^1^ΨpA was assayed by using UV spectroscopy. The UV spectra of UpA, ΨpA, and m^1^ΨpA are similar, though that of m^1^ΨpA is shifted slightly to higher wavelengths (Fig. S3). To assess the three dinucleotides as substrates for ptRNases, we sought changes in absorbance near 280 nm that accompany dinucleotide cleavage (Witzel and Barnard 1962). Those changes were apparent, albeit small (Fig. S4), and we chose an optimal wavelength for monitoring each cleavage reaction (Table S2). We performed assays at the optimal pH for catalysis by RNase A (pH 6.0) and RNase 1 (pH 7.5), which reflects their physiological environments (Lomax et al. 2017).

We found that all three dinucleotides are substrates for both RNase A and RNase 1 (Fig. S5). Plots of the initial rates are shown in Fig. 2, and steady-state kinetic parameters are listed in Table 1. In general, UpA is a better substrate than either ΨpA or m^1^ΨpA. The differences, which are more significant for RNase A, are due primarily to decreases in the value of *k*_cat_ rather than changes in the value of *K*_M_. The largest differences are nearly 10-fold, which are for the *k*_cat_/*K*_M_ values of RNase A. UpA is also a better substrate than either ΨpA or m^1^ΨpA at the pH optimum of the other enzyme, that is, RNase A at pH 7.5 and RNase 1 at pH 6.0 (Table S3; Fig. S6).

**FIGURE 2.**
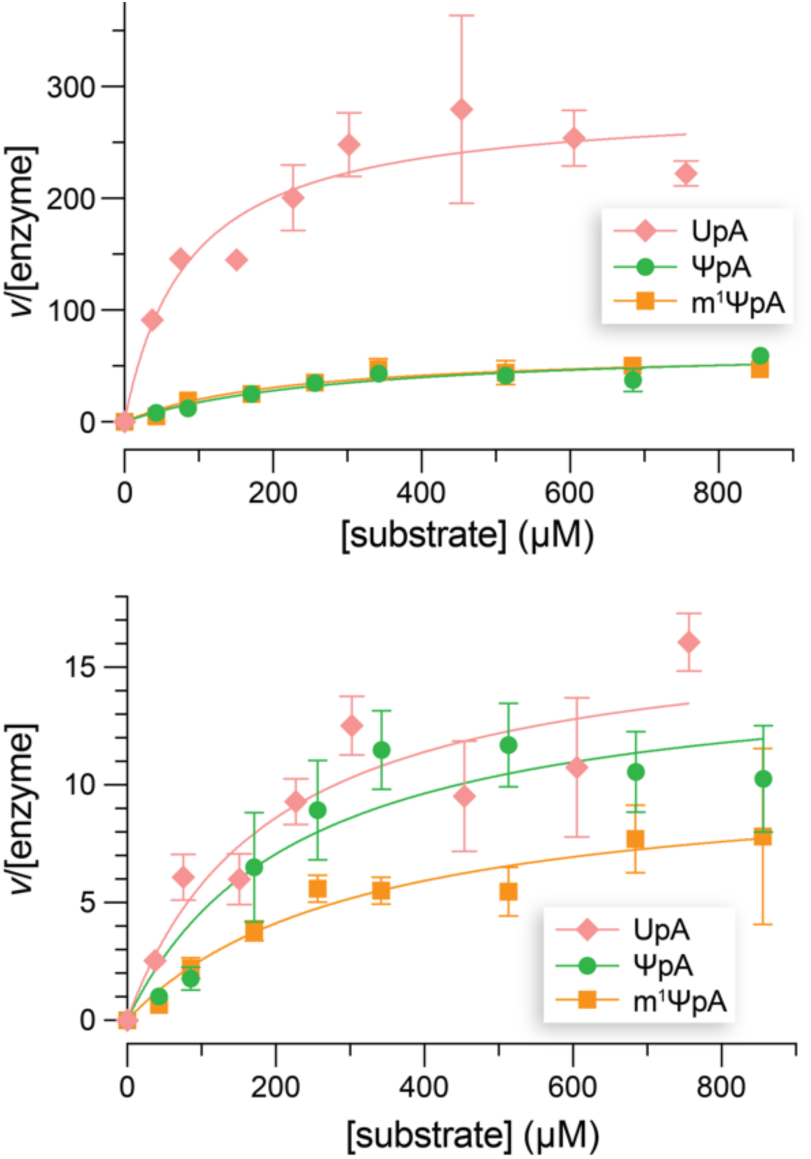
Initial rates for catalysis of the cleavage of dinucleotide substrates by RNase A (pH 6.0) and RNase 1 (pH 7.5) at 25 °C. Values are the mean ± SD of 4 replicates. The resultant steady-state kinetic parameters are listed in Table 1.

**Table 1:** Steady-state kinetic parameters for the cleavage of UpA, ΨpA, and m^1^ΨpA by RNase A and RNase 1

| Enzyme | pH | Substrate | $k_{cat}$ (s<sup>-1</sup>) | $K_M$ (μM) | $k_{cat}/K_M$ (10<sup>5</sup> M<sup>-1</sup> s<sup>-1</sup>) |
| --- | --- | --- | --- | --- | --- |
| RNase A | 6.0 | UpA | 313 ± 25 | 143 ± 39 | 21.8 ± 6.2 |
| | | ΨpA | 65 ± 7 | 305 ± 84 | 2.1 ± 0.6 |
| | | m¹ΨpA | 70 ± 7 | 300 ± 82 | 2.3 ± 0.7 |
| RNase 1 | 7.5 | UpA | 17 ± 2 | 253 ± 77 | 0.67 ± 0.22 |
| | | ΨpA | 14 ± 2 | 254 ± 88 | 0.58 ± 0.21 |
| | | m¹ΨpA | 9 ± 1 | 328 ± 116 | 0.30 ± 0.12 |

### Structural interactions of U, Ψ, and m^1^Ψ with RNase A

Next, we sought to understand the basis for the decrease in catalytic efficiency for the cleavage of UpΨ and Upm^1^Ψ versus UpA. To do so, we focused initially on structure. Uridine 2′,3′-cyclic vanadate (U>v) is a potent inhibitor of RNase A, superior to uridine or inorganic vanadate alone (Lindquist et al. 1973). This complex is thought to be a mimic of the enzymic transition state, albeit an imprecise one (Krauss and Basch 1992; Messmore and Raines 2000b). Given that precedent, we sought and obtained crystal structures of RNase A bound to U>v, Ψ>v, and m^1^Ψ>v, which we solved at resolutions of 1.83 Å, 1.70 Å, and 1.71 Å, respectively. The data and refinement statistics for each structure are listed in Table S4. In these structures, we found that each vanadyl group is in a tetrahedral geometry (Fig. S7), instead of the previously reported trigonal bipyramidal geometry(Wlodawer et al. 1983; Ladner et al. 1997). That valency, however, did not alter the location of the uridine moiety. In addition, we observed decavanadates in each crystal lattice (Fig. S8). This byproduct of the nucleoside plus vanadate complexation reaction is likewise known to be an inhibitor of catalysis by RNase A (Messmore and Raines 2000a), though its structural interaction with the enzyme had not been described previously.

An overlay of the U>v, Ψ>v, and m^1^Ψ>v moieties bound to RNase A is shown in Fig. 3. The binding of the ligand is virtually identical in the three structures. Specifically, each uridine ring is in the *anti* conformation and interacts with Thr45 in a similar manner. The near-identity of these three structures indicates that the observed differences in steady-state kinetic parameters (Table 1) do not arise primarily from differences in structure.

**FIGURE 3.**
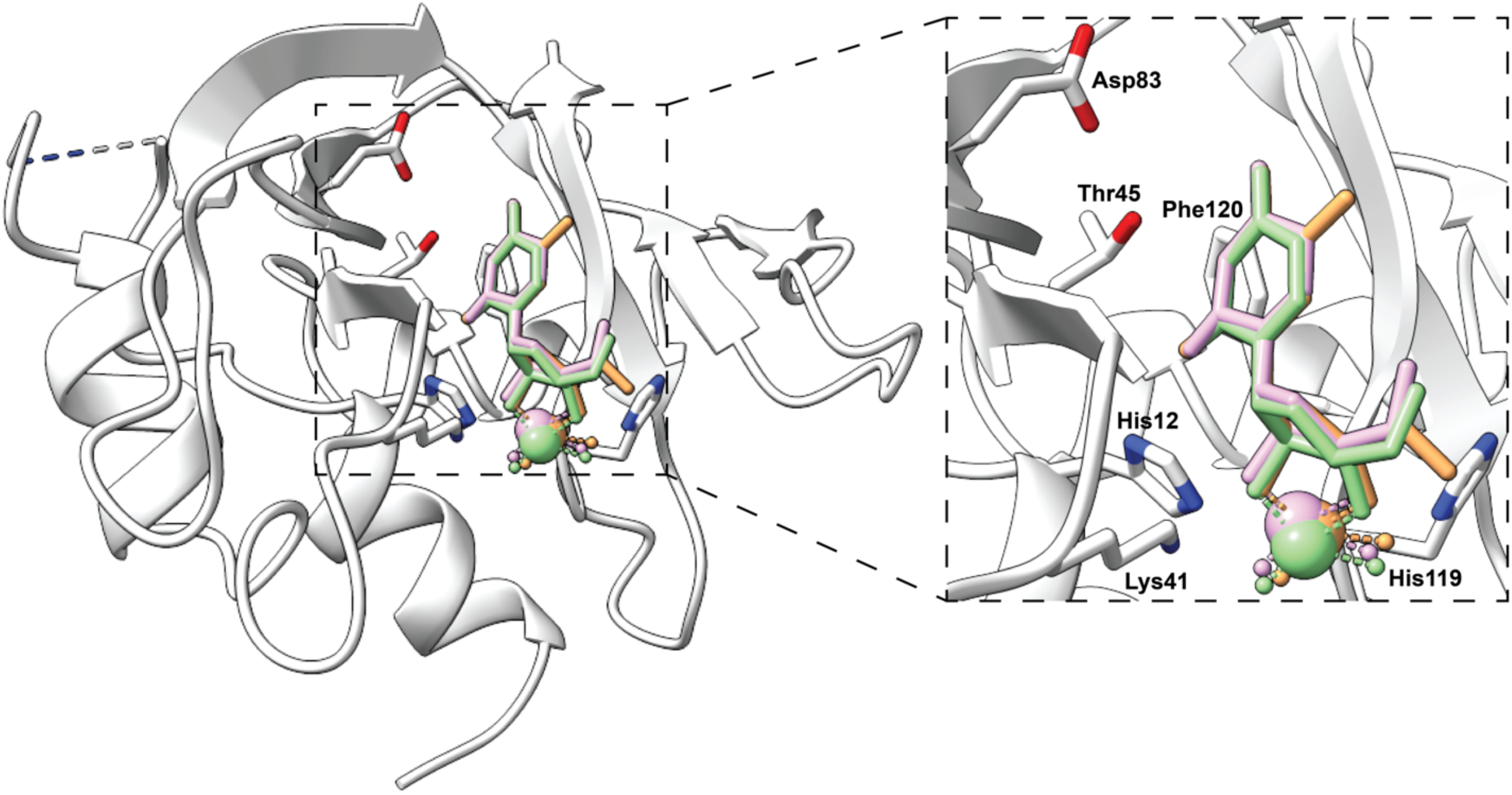
Overlay of the crystal structures of RNase A bound to U>v (pink), Ψ>v (green), and m^1^Ψ>v (orange). The protein representation (white) is from the Ψ>v crystal structure, and key residues for binding and catalysis are indicated explicitly. The nucleobase···Thr45 distances are RNase A·U>v: N^3^···O^γ1^, 2.72 Å; O^2^···N, 2.78 Å; RNase A·Ψ>v, N^3^···O^γ1^, 2.80 Å; O^4^···N, 2.83 Å; and RNase A·m^1^Ψ>v, N^3^···O^γ1^, 2.81 Å; O^4^···N, 2.80 Å.

### Simulation of RNase 1 binding to UpA, ΨpA, and m^1^ΨpA

We were unable to obtain the crystal structure of RNase 1 bound to a nucleoside vanadate. That is not surprising, as RNase 1 has been crystallized in a complex with its inhibitor protein (Johnson et al. 2007) but not alone or in a complex with a small-molecule ligand. Accordingly, we sought insight about RNase 1 from molecular docking studies and molecular dynamics (MD) simulations with the three dinucleotide substrates. The structures of UpA, ΨpA, and m^1^ΨpA were optimized at the M06-2X/ 6-31+G(d,p) level of theory (Zhao and Truhlar 2008a; Zhao and Truhlar 2008b) and docked into the active site of RNase A (Fig. S9) and RNase 1 (Fig. S10). MD simulations for each substrate were initiated from the enzyme·substrate complexes resulting from the docking studies and were performed for 1500 ns of simulation time. For each simulation, an average structure was generated using 7,500 frames from the last 750 ns. Overlays of the dinucleotide substrates bound to RNase 1 obtained from the MD simulations with the crystal structures of the nucleoside vanadates bound to RNase A are in accord (Fig. 4), suggesting that each substrate binds in a similar manner. We also performed MD simulations of the dinucleotide substrates bound to RNase A and observed highly similar binding modes, consistent with the crystal structures (Fig. S11).

**FIGURE 4.**
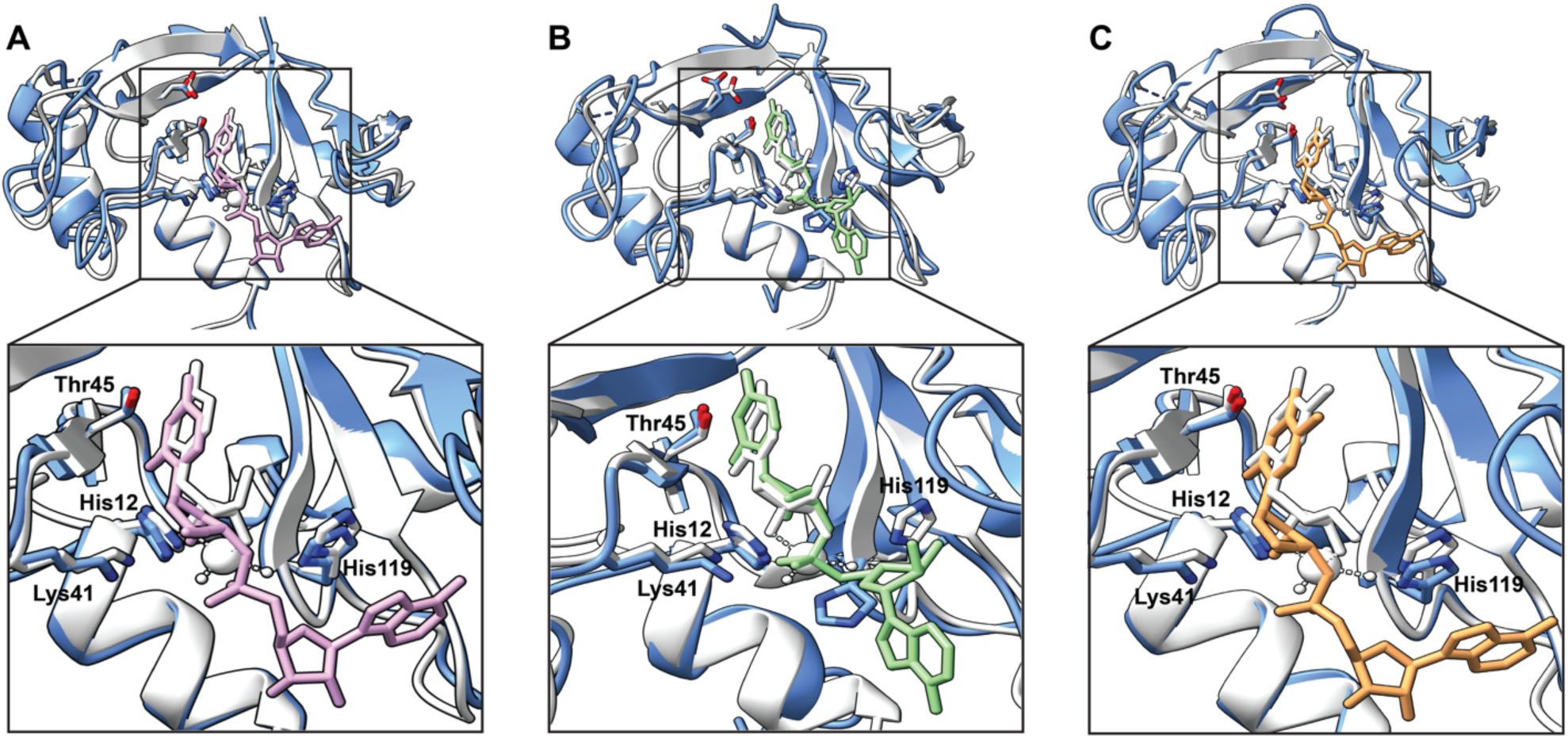
Overlay of UpA (*A*, pink), ΨpA (*B*, green) and m^1^ΨpA (*C*, orange) bound to RNase 1 (blue) obtained from the MD simulations with the respective crystal structures of U>v (*A*, white), Ψ>v (*B*, white), and m^1^Ψ>v (*C*, white) bound to RNase A (white).

A major challenge in computational chemistry is the development of methods to reliably predict the binding free energy (Δ*G*_bind_) of a ligand to its receptor in a complex. Various methods have been developed, ranging from simple and fast scoring functions (Rajamani and Good 2007) to more rigorous but time-intensive approaches such as free energy perturbation (Michel and Essex 2010; Cournia et al. 2021). In between are the so-called end-point methods that utilize molecular dynamics simulations, which take into account the dynamic nature of the bimolecular interaction, along with a molecular mechanics (MM) potential (Wang et al. 2019).

MMGBSA (molecular mechanics-generalized Born surface area) is one of the widely used end- point methods for estimating the energetics of substrate binding (Kollman et al. 2000; Tsui and Case 2000; Rastelli et al. 2010; Ylilauri and Pentikäinen 2013; Mikulskis et al. 2014; Genheden and Ryde 2015). Here, binding free energies from the MD simulations were sampled from the last 750 ns (using 7,500 equally spaced frames).

The three dinucleotides share a similar binding pattern to RNase 1, with UpA generally showing the highest affinity and m^1^ΨpA generally showing the lowest affinity in accord with the experimental data. The decomposition of binding energies into gas-phase energy contributions (electrostatic and van der Waals) and solvation free energy components (polar and nonpolar) shows that electrostatics dominate the overall binding of each substrate (Table 2). Similar nonpolar contributions imply that each substrate has a similar exposure to solvent and comparable hydrophobic properties. To further assess the energetic contributions of the important residues within the active site of RNase 1, we decomposed the binding energies on a pairwise per-residue basis. Once again, we observed similar contributions from four key residues, His12, Thr45, Asp83, and Phe120, to the overall binding free energies (Fig. 5). Lys41 makes a stronger contribution to binding the pseudoridines, and His119 makes a weaker contribution.

**FIGURE 5.**
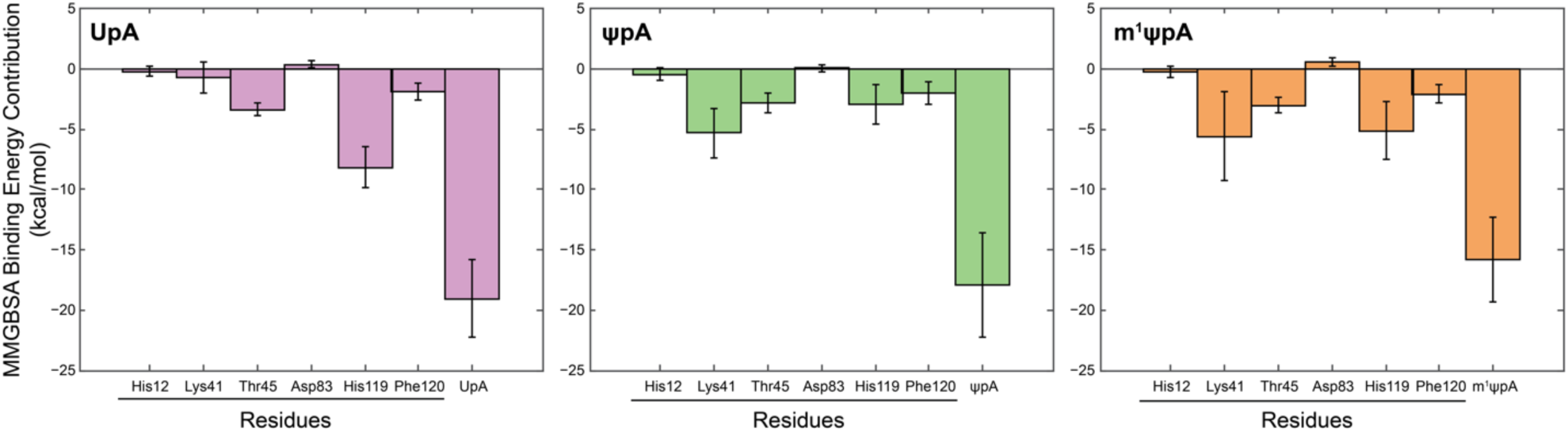
MMGBSA binding energy contributions of key residues of RNase 1 to dinucleotide substrates.

**TABLE 2.** Decomposition of average MMGBSA binding energies (Δ*G*_bind_, kcal/mol) of RNase 1 for dinucleotide substrates into gas-phase energy contributions (Δ*E*_van der Waals_ and Δ*E*_electrostatics_) and solvation free energy components (*G*_polar_ and *G*_nonpolar_).

| Energy Component | UpA | ΨpA | m<sup>1</sup>ΨpA |
| --- | --- | --- | --- |
| $\Delta E_{\text{van der Waals}}$ | -48.75 | -44.57 | -45.75 |
| $\Delta E_{\text{electrostatics}}$ | -275.43 | -263.43 | -286.11 |
| $\Delta G_{\text{polar}}$ | 283.16 | 268.15 | 294.64 |
| $\Delta G_{\text{nonpolar}}$ | -6.28 | -5.67 | -5.59 |
| $\Delta G_{\text{bind}}$ | -47.29 | -45.52 | -42.81 |

### Basis for the differential catalysis of UpA, ΨpA, and m^1^ΨpA cleavage

A comparison of the crystal structures of the three nucleoside vanadates bound to RNase A did not reveal a basis for the observed difference in cleavage kinetics. The same observation holds true for the molecular dynamics simulation of the binding of different UpA derivatives to RNase 1, where no significant differences were detected that could explain the difference in cleavage kinetics. Having ruled out binding modes as the main contributor to differential kinetics, we wondered whether an unappreciated electronic effect could explain the observed difference in rate constants for the pseudouridine substrates.

In uridine, the pyrimidine nucleobase is linked to the ribose by a β-*N*-glycosidic bond, whereas in the pseudouridines, that linkage is a β-*C*-glycosidic bond (Fig. 1). On the Pauling scale, the electronegativity of nitrogen (*χ* = 3.0) is greater than that of carbon (*χ* = 2.5) (Pauling 1939). Moreover, *N*^1^ of uridine is conjugated to functional groups in the nucleobase (Figure 1), further increasing its ability to withdraw electron density. We reasoned that the enhanced inductive effect of *N*^1^ in uridine decreases the p*K*_a_ of its 2′-hydroxy group relative to that in pseudouridines. The deprotonation of that 2′-hydroxy group is necessary for cleavage by ptRNases (Findlay et al. 1961; Cuchillo et al. 2011), and a lower p*K*_a_ should increase reactivity (Dantzman and Kiessling 1996).

To provide insight into the relative acidities of the 2′-hydroxy groups of the substrates, we modeled the first step in the catalysis of RNA cleavage. In that step, the imidazolyl group in His12 acts as a base that abstracts a proton from the 2′-hydroxy group (Jackson et al. 1994; Thompson and Raines 1994). In UpA, the p*K*_a_ of the uridylyl 2′-hydroxy group is 12.54 (Järvinen et al. 1991). We used density functional theory with the SMD solvation model to calculate the free energy that accompanies a proton transfer between three nucleoside 3′-methylphosphates and 4-methylimidazole. We selected this approach to circumvent issues with accurately predicting the solvation energy of a proton (Ho and Coote 2010; Prasad and Tantillo 2021). The resulting free energies suggest that the p*K*_a_ of the 2′-hydroxy group of uridine 3′-phosphate is lower than those of pseudouridine 3′-methylphosphate and *N*^1^-methylpseudouridine 3′- methylphosphate, which are similar (Table 3).

**Table 3.** Relative free energies for the deprotonation of the 2′-hydroxy group of nucleoside 3′-methylphosphates by 4-methylimidazole with respect to the sum of the reactant energies at infinite separation ([*SMD*-H_2_O] M06-2X/6- 31+G(d,p)).

| Nucleobase (R) | $\Delta G$ (kcal/mol) |
| --- | --- |
| uracil | 18.4 |
| pseudouracil | 21.3 |
| $N^1$-methylpseudouracil | 21.0 |

We reasoned that the differential p*K*_a_ for uridine versus a pseudouridine would be manifested in their intrinsic reactivity. To test this hypothesis, we assessed the uncatalyzed cleavage of UpA and m^1^ΨpA. At pH 6.0 and 25 °C, the rate constant for the uncatalyzed cleavage of UpA is *k*_uncat_ = 5 × 10^−9^ s^−1^, which corresponds to *t*_1/2_ = 4 years (Thompson et al. 1995). To accelerate the time course, we chose to monitor the cleavage reactions at a higher pH (10.0) and temperature (90 °C), using a buffer (2-(cyclohexylamino)ethanesulfonic acid; CHES) with a p*K*_a_ of known temperature-dependence (Roy et al. 1997). We note that the p*K*_a_ of the *N*^3^–H imido group of uridine and *N*^1^-methylpseudouridine are similar (Jones et al. 2022), and both will be mostly unprotonated at pH 10.0.

We used ^31^P NMR spectroscopy to monitor the non-enzymatic cleavage of UpA and m^1^ΨpA (Figs. S14 and S15). We found that UpA (*k*_uncat_ = 4.7 ± 0.1 × 10^−6^ s^−1^; *t*_1/2_ = 41 h) is more vulnerable to spontaneous cleavage than is m^1^ΨpA (*k*_uncat_ = 2.5 ± 0.1 × 10^−6^ s^−1^; *t*_1/2_ = 77 h) (Fig. 6). These experimental data are consistent with the inductive effect of the uracil nucleobase on the p*K*_a_ of the 2′-hydroxy group in uridine (Table 3).

**FIGURE 6.**
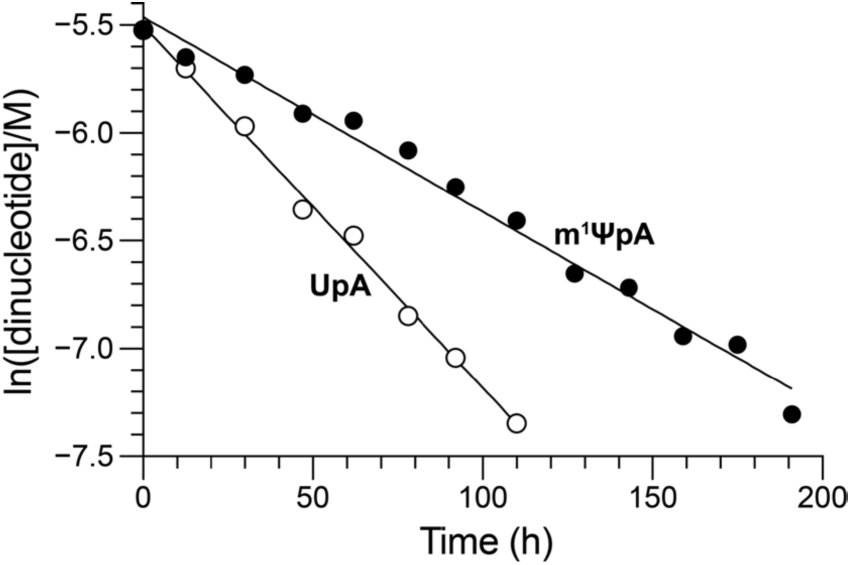
Non-enzymatic cleavage rate of UpA and m^1^ΨpA in 0.10 M CHES–NaOH buffer, pH 10.0, containing NaCl (0.10 M) at 90 °C as monitored with ^31^P NMR spectroscopy.

## Discussion

ptRNases degrade RNA in a largely non-specific manner, governed only by the requirement that cleavage of the P–O^5′′^ bond occurs directly after a pyrimidine residue. We discovered that nucleotides containing Ψ and m^1^Ψ are accommodated in the active site of both RNase A and RNase 1 and allow for catalysis, though at a lower level than U. Additionally, we found that the binding of Ψ and m^1^Ψ occurs in a similar manner to that of U and obtain, to the extent of our knowledge, the first crystal structure of a protein bound to m^1^Ψ and one of few crystal structures of a protein bound to Ψ (Hoang and Ferré-D’Amaré 2001).

Catalysis by RNase 1 had not been examined previously with a dinucleotide substrate. With a tetranucleotide substrate (Fig. S2), catalysis by RNase A is only twofold less efficient than by RNase 1 (Table S1). This difference increases to 20-fold with a dinucleotide substrate. Although RNase A and RNase 1 are homologous and share 68% of their amino acid residues, the two enzymes evolved in distinct niches and have different physiological roles (Eller et al. 2014). For example, RNase A could be an important scavenger of inorganic phosphate for ruminants (Barnard 1969), a role that benefits from the turnover of even the smallest RNA substrates.

Along these lines, we found that RNase A is more sensitive than RNase 1 to uracil modifications. This drop off could be related to the more specific function of RNase A in the ruminant gut, where RNase A might have evolved to turnover RNA containing canonical nucleotides. Unlike RNase A, human RNase 1 can turnover a large diversity of RNA substrates, including double-stranded RNA (Lomax et al. 2017), the RNA strand of RNA:DNA hybrids (Potenza et al. 2006), and even poly(A), albeit at a low level (Sorrentino 1998). These enzymatic activities are consistent with RNase 1 being expressed at significant levels in many tissues and having functions that require a broader substrate scope that modulates the innate immune response, vascular homeostasis, and scavenging extracellular RNA (Sorrentino 2010; Koczera et al. 2016; Garnett et al. 2019). Apparently, RNase 1 can accommodate pyrimidine variations better than RNase A. In living systems, RNA occurs in diverse sizes, structures, compositions, and sequences, and the trends reported here might be amplified or reduced, depending on the context as well as the biochemical environment.

Structurally, we observed little difference in the binding of the three transition state analogs to RNase A. Nor did molecular dynamics simulations reveal significant overall differences in substrate binding to either RNase A or RNase 1. We discovered, however, that the differential glycosidic connectivity of the nucleobase in uridine and the pseudouridines alters the p*K*_a_ of the 2′-hydroxy group, decreasing the intrinsic reactivity of pseudouridine nucleotides. In accord, we found a lesser difference in the enzymatic turnover of UpA, ΨpA, and m^1^ΨpA at pH 7.5 than at pH 6.0. This discovery expands our fundamental understanding of the most abundant modified nucleoside in natural RNA, and has implications for RNA biology as well as the ongoing development of RNA-based vaccines and therapeutic agents.

## Materials and Methods

### Production and purification of RNase 1

Human RNase 1 was produced in *E. coli* without its signal peptide and with an N-terminal methionine residue as described previously (Ressler et al. 2019), with minor modifications. The expression plasmid was transformed into BL21(DE3) cells. A starter culture (50 mL) was inoculated from a single colony and grown overnight at 37 °C in TB containing ampicillin (200 µg mL^−1^) with constant shaking at 250 rpm. Cultures (1.0 L) were initiated at OD_600 nm_ = 0.05 from the starter culture and grown at 37 °C in TB containing ampicillin (200 µg mL^−1^) with constant shaking at 250 rpm. Gene expression was induced with IPTG (final concentration: 1.0 mM) when the cultures reached OD_600 nm_ = 1.8–2.2 and were grown for an additional 3 h at 37 °C. Cells were pelleted by centrifugation at 6,000*g* for 15 min at 4 °C, and cell pellets were stored at −80 °C until resuspension and lysis.

Cell pellets containing RNase 1 were resuspended in 20 mM Tris–HCl buffer, pH 7.6, containing 10 mM EDTA (1 g of wet pellet per 10 mL of buffer). Cells were lysed at 19.0 kpsi at 4 °C using a benchtop cell disruptor from Constant Systems (Daventry, England). Inclusion bodies were isolated by centrifugation at 30,000*g* for 1.5 h at 4 °C. The resultant inclusion bodies were resuspended in 20 mM Tris–HCl buffer, pH 8.0, containing guanidine–HCl (7 M), DTT (0.10 M), and EDTA (10 mM) for 2 h at room temperature (4 mL of buffer per 1 L of expression culture). The solution was then diluted 10-fold with 20 mM acetic acid, and the insoluble material was removed by centrifugation at 16,500*g* for 30 min at 4 °C. The clarified supernatant was dialyzed against 16 L of 20 mM acetic acid overnight at 4 °C using a 3.5k MWCO bag. The dialysate was then subjected to centrifugation at 30,000*g* for 1 h at 4 °C to remove additional insoluble material. To fold the RNase 1, the clarified supernatant was added dropwise with gentle stirring to 100 mM Tris–HCl buffer, pH 7.8, containing NaCl (0.10 M), reduced glutathione (1.0 mM), and oxidized glutathione (0.2 mM). This incubation was continued at 4 °C without stirring for at least 2 days. The pH of the solution was then adjusted to 5.0 by adding 3 M sodium acetate buffer, pH 5.0, and the resulting solution was passed through a 0.45-μm filter. The sterilized solution was concentrated using an Amicon Stirred Cell concentrator from EMD Millipore (Billerica, MA) with 10-kDa filters. Further purification was done with an ÅKTA pure FPLC system from Cytiva (Marlborough, MA). Gel-filtration chromatography was performed with a Superdex HiLoad 26/600 75 pg gel filtration column and 50 mM sodium acetate buffer, pH 5.0, containing NaCl (0.10 M) and sodium azide (0.05% w/v). Fractions containing the ribonuclease were pooled and purified further by chromatography with a HiTrap SP cation-exchange column and 50 mM sodium acetate buffer, pH 5.0, containing a linear gradient of NaCl (0.35–0.70 M) over 35 column volumes. Fractions containing RNase 1 were pooled, concentrated, and buffer-exchanged into 50 mM Tris–HCl, pH 7.5, containing NaCl (50 mM) using an Amicon 15 mL 10kDa MWCO spin concentrator. The identity of the protein was validated with QTOF mass spectrometry and SDS–PAGE. Aliquots were flash-frozen in N_2_(l) and stored at −70 °C. Typically, ∼2 mg of RNase 1 was obtained per liter of culture.

### Kinetic measurements of dinucleotide cleavage

UpA, ΨpA, and m^1^ΨpA were synthesized from their component nucleosides by using the phosphoramidite method (Caruthers 2011) (Scheme 1). (For experimental details, see the Supplemental Information.) The cleavage of UpA, ΨpA, and m^1^ΨpA by RNase A and RNase 1 was assayed by monitoring the change in absorbance of the substrate near 280 nm (Witzel and Barnard 1962). (Note: a coupled assay using adenosine deaminase (Ipata and Felicioli 1968) was not feasible because commercial sources of adenosine deaminase catalyze the hydrolysis of the adenosine in the substrate as well as the product.) For each substrate, the exact wavelength was chosen by analyzing UV spectra before and after cleavage and choosing the wavelength with the maximal difference (Table S2 and Fig. S4). Assays were performed with a Spark plate reader from Tecan (Männedorf, Switzerland) in 96-well half-area UV star microplates with a final volume of 50 µL. For RNase A, reactions were performed in 0.10 M DEPC-treated OVS-free MES–NaOH buffer, pH 6.0, containing NaCl (0.10 M), which is the pH at which RNase A is most active (Lomax et al. 2017). RNase 1 reactions were performed in 0.10 M Tris–HCl buffer, pH 7.5, containing NaCl (0.10 M) NaCl, which is the pH at which RNase 1 is most active (Lomax et al. 2017). This buffer was made from DEPC-treated water and a DEPC-treated stock solution of NaCl (1.0 M). Assays were conducted at 25 °C with a substrate concentration of 50–1000 µM. Due to the lack of sensitivity of this assay, the concentration of ribonuclease in each reaction was optimized to have a high dynamic range while maintaining linearity in the initial region. For RNase A, concentrations of 1 nM, 4 nM and 10 nM were used; for RNase 1, concentrations of 10 nM, 20 nM, and 20 nM were used for UpA, ΨpA, and m^1^ΨpA, respectively. To obtain the baseline absorbance of a well and ensure no contamination from exogenous ribonucleases, 25 µL of a 2× solution of substrate in the appropriate buffer was added to each well, and the absorbance was measured over 2 min. Then, a 2× solution of RNase A or RNase 1 was added to the substrate, and the absorbance was measured over 20 min. To obtain the absorbance of fully cleaved substrate, 25 µL of a solution of RNase A (4 µM) was added to 25 µL of the 2× solution of substrate in a separate well, and the absorbance was measured over 10 min. The measurements were performed at wavelengths optimized for each enzyme–substrate pair as listed in Table S2. Initial rates were plotted against substrate concentration and fitted to the Michaelis–Menten equation to determine values of *k*_cat_ and *K*_M_. (See the Supplementary Information for additional details.)

For the pH-dependent experiments, assays were performed as above, except in the buffer corresponding to the desired pH. Values of *k*_cat_/*K*_M_ were determined by performing linear fits in regions of low [S]. (See the Supplementary Information for additional details.)

For the non-enzymatic cleavage experiments, a solution of UpA (4.0 mM) and a solution of m^1^ΨpA (4.0 mM) were prepared in 0.10 M CHES–NaOH buffer, pH 10.0, containing NaCl (0.10 M) under ribonuclease-free conditions. The solutions were added to NMR tubes. Sealed capillaries containing phenylphosphonic acid (50 mM) in D_2_O containing NaCl (0.10 M) were placed into the tubes as a reference. Both tubes were kept under identical conditions at all times. The tubes were incubated at 90 °C for the specified amount of time. Before acquiring ^31^P NMR spectra, the samples were cooled to room temperature. Undecoupled ^31^P NMR spectra (scans: 64, relaxation delay: 12 s, center: 10 ppm, spectral width: 50 ppm) were obtained at each time point. To calculate the concentration of UpA or m^1^ψpA at a particular time point, the integral of the signal from UpA (−0.69 ppm) or m^1^ΨpA (−0.48 ppm) was normalized to the integral of the signal from phenylphosphonic acid (15.99 ppm). Signals from each 2′,3′-cyclic phosphate (∼20 ppm) and 2′- and 3′-phosphates (∼3–4 ppm) appeared during the time course of the reaction (Figs. S15 and S16). Integrals of sufficient accuracy could be obtained only to a dinucleotide concentration of ≥0.8 mM. Spectra were processed with MestReNova software (version 14.1.1-24571) using automatic baseline and phase correction tools. Rate constants were obtained with linear least-squares regression analyses using Origin 2021 software (version: 9.8.0.200).

### Crystallization of RNase A with nucleoside vanadates

RNase A was crystallized as described previously with some minor modifications. RNase A was dissolved in water to 20 mg/mL stock concentration. Crystals were initially grown via the hanging drop method using a reservoir of 20 mM sodium citrate buffer, pH 5.5, containing 20% PEG 4000 (20% w/v) and a drop that was a 1:1 solution of reservoir:RNase A stock solution.

The drops were incubated at 16 °C. Crystals appeared in *ca*. 7–10 days. Additional crystals were obtained by seeding small or fragmented crystals from the initial growth via a seeding tool into various solutions via the hanging drop method. In general, the seeded crystals grew larger at 16 °C with reservoir solutions of 20 mM sodium citrate buffer, pH 5.5, or 50 mM imidazole– HCl buffer, pH 5.5, containing PEG 4000 (20–25% w/v) and *tert*-butanol (0–10% w/v) and drops that were 1:1 solutions of reservoir:RNase A stock solution. Seeded crystals grew in *ca*. 3–7 days. After crystals formed, they were stable at 16–25 °C for at least a month in the dark. The highest quality crystals grew in 20 mM sodium citrate buffer, pH 5.5, containing PEG 4000 (25% w/v) and were used for subsequent soaking experiments with Ψ>v and m^1^Ψ>v. For the U>v experiments, crystals were formed via seeding into 50 mM imidazole–HCl, pH 5.5, containing PEG 4000 (20% w/v).

To obtain ligand-bound structures, nucleoside vanadates were soaked into RNase A crystals. The nucleoside vanadate solutions were prepared as described previously with minor modifications (Ladner et al. 1997). Briefly, uridine or a pseudouridine (45 mg) was mixed with ammonium vanadate (NH_4_VO_3_; 105 mg) in 6.1 mL of 50 mM imidazole–HCl buffer, pH 5.2, and the resulting solution was heated to 60 °C for 20 min on a hotplate with stirring. The resulting solutions were yellow and, after cooling to room temperature, constituted the stock solution for each ligand. For U>v, crystals were transferred into hanging drops containing a 2:1 mixture of U>v stock solution:25 mM imidazole–HCl buffer, pH 5.5, containing PEG 4000 (30% w/v) over a reservoir of the same composition. These hanging drops were incubated at room temperature for 2 days. Prior to freezing and data collection, the crystals were dipped into cryoprotectant conditions, which were 16.7 mM imidazole–HCl buffer, pH 5.5, containing U>v stock solution (33% v/v), PEG 4000 (30% w/v), and glycerol (5% v/v), before they were flash-cooled in a stream of cryogenic N_2_(g) for data collection. For Ψ>v, crystals were transferred into hanging drops of 16.7 mM imidazole–HCl buffer, pH 5.5, containing Ψ>v stock solution (33% v/v), PEG 4000 (30% w/v), and glycerol (5% v/v). These hanging drops were incubated at room temperature for 2 days before direct freezing in cryogenic N_2_(g) and data collection. For m^1^Ψ>v, crystals were transferred into hanging drops containing of 16.7 mM imidazole–HCl buffer, pH 5.5, containing m^1^Ψ>v stock solution (33% v/v), PEG 4000 (30% w/v), and glycerol (5% v/v).

These hanging drops were incubated at room temperature for 1 day before direct freezing in cryogenic N_2_(g) and data collection.

### Data collection, processing and structural determination

Data were collected on a Rigaku Micromax-007 rotating anode with Osmic VariMax-HF mirrors and a Rigaku Saturn 944 detector. The obtained data were processed with XDS (Kabsch 2010). Phaser (McCoy et al. 2007), as implemented in PHENIX (Liebschner et al. 2019), was used to solve the structures by molecular replacement, using the protein coordinates from the structure of RNase A in a complex with cytidine 3′-phosphate (PDB entry 5ogh (Prats-Ejarque et al. 2019)). Ligand structures were modeled and optimized with Gaussian 16 at the M06-2X/ 6- 31+G(d,p) level (Zhao and Truhlar 2008a; Zhao and Truhlar 2008b; Frisch et al. 2016) using restraints conforming to the configuration of the vanadyl group in the structure of RNase A in complex with U>v (PDB entry 1ruv (Ladner et al. 1997)). Structures were refined with PHENIX (DiMaio et al. 2013) and manual fitting in Coot (Emsley et al. 2010). Data and refinement statistics are listed in Table S4.

### Molecular docking

The structures of UpA, ѰpA, and m1ѰpA were optimized at the M06-2X/ 6-31+G(d,p) level of theory using Gaussian (Zhao and Truhlar 2008a; Zhao and Truhlar 2008b; Frisch et al. 2016) and docked into the active site of RNase A and RNase 1 using AutoDock Vina (Trott and Olson 2010). During docking, the side chains of catalytic residues His12, His119, and Lys41 were treated as flexible, while the rest of the enzyme was treated as rigid. The resulting docked poses were scored, and poses were selected in which (1) the pyrimidine and Thr45, (2) the 2′ hydroxy group and His12, (3) the phosphoryl group and Lys41, and (4) the 5′′-oxygen and His119 were within the interaction distances (Figs. S8 and S9).

### Molecular dynamics simulations set-up

Molecular dynamics simulations were based on the complexes that resulted from the docking of UpA, ѰpA, and m^1^ѰpA into the active site of RNase A and RNase 1. The structure of RNase A was from PDB entry 1ruv (Ladner et al. 1997). The protonation states of RNase A residues were adjusted to those at pH 6.0. The structure of RNase 1 was based on chain X of PDB entry 2q4g (Johnson et al. 2007). Missing residues (*i*.*e*., the N-terminal lysine and C-terminal threonine) were modeled with MODELER software (Webb and Sali 2016). The protonation states of RNase 1 residues were adjusted to those at pH 7.5.

### Molecular dynamics simulation protocol

Molecular dynamics simulations were performed on cuda-enabled gpus of pmemd (Particle Mesh Ewald Molecular Dynamics) in Amber 2022 (Case et al. 2023). We used the ff14SB and gaff parameter sets for the protein and ligands, respectively (Wang et al. 2004; Maier et al. 2015). We also performed the simulations using the RNA force field OL3 (Zgarbova et al. 2011) and modrna08 (Aduri et al. 2007) for canonical and modified dinucleotide ligands, respectively (Fig. S12). Restrained electrostatic potential charges (RESPs) were calculated with the R.E.D. server (Bayly et al. 1993; Vanquelef et al. 2011). Each system was solvated in the TIP3P water model and neutralized in 0.10 M NaCl (Jorgensen et al. 1983). Each system was minimized and heated to 300 K using Langevin dynamics with a collisional frequency of 1 ps in the NVT ensemble over 100 ps with harmonic restraints of 10.0 kcal mol^−1^ Å^−2^ on protein and ligand.

Further equilibration was performed in the NPT ensemble using isotropic position scaling and a pressure relaxation time of 2 ps at 300 K with harmonic restraints on the protein and ligand starting at 5.0 kcal mol^−1^ Å^−2^ and lifted slowly, giving a total of 12 ns of restrained equilibration and 20 ns of unrestrained equilibration time. The nonbonded cutoff was 9 Å. Each system was subjected to a 1500 ns production run in the NPT ensemble using time steps of 2 fs with Langevin dynamics and a Monte Carlo barostat at 300 K. The long-range electrostatic interactions were calculated using the particle mesh Ewald (PME) method.

### Analysis protocols for MD simulations

The binding free energies (Δ*G*_bind_ in kcal/mol) were computed by utilizing the molecular mechanics-generalized Born surface area (MMGBSA) method implemented in Amber 2022 using the last 750 ns (using 7500 equally spaced simulation frames) and decomposed on a pairwise per-residue basis (Kollman et al. 2000; Tsui and Case 2000; Rastelli et al. 2010; Ylilauri and Pentikäinen 2013; Mikulskis et al. 2014; Genheden and Ryde 2015).

In the MMGBSA method, the free energy for binding is expressed as

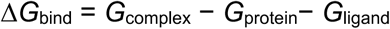

where Δ*G*_bind_ is the binding free energy and Δ*G*_complex_ Δ*G*_protein_, and Δ*G*_ligand_ are the free energies of complex, protein, and ligand, respectively. Δ*G*_bind_ can be decomposed into

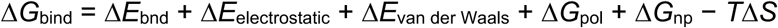

where the first three terms are MM energy components in gas phase from bonded (bond, angle, dihedral), electrostatic, and van der Waals interactions. Δ*G*_pol_ is the polar contribution to the solvation free energy, which can be obtained by using the generalized Born (GB) model. The nonpolar contribution to the solvation free energy, Δ*G*_np_, is estimated using the solvent-accessible surface area (SASA). Finally, *T*Δ*S* is the change in conformational entropy, which is often neglected when ranking binding free energies of similar ligands (Gohlke and Case 2004; Zhou and Madura 2004). In this study, the binding free energies were calculated using the single trajectory approach, resulting in the cancellation of bonded energy terms, and the entropy contribution is omitted to reduce computational cost. The GB calculations were performed using parameters developed previously (igb = 5 (α = 1.0, β = 0.8, γ = 4.85)) (Onufriev et al. 2004).

An average structure from each simulation was generated by k-means cluster analysis, and root-mean-square deviation (RMSD) analysis was performed to assess the stabilization of the protein structure (Figs. S13–S15), employing the CPPTRAJ module implemented in Amber Tools23 (Roe and Cheatham 2013; Case et al. 2023).

## Acknowledgments

We thank Alia A. Kassim for help in determining the extinction coefficients for UpA, ΨpA, and m^1^ΨpA. C.S.G. was supported by a National Defense Science and Engineering Graduate Fellowship sponsored by the Air Force Research Laboratory. B.S. was supported by a Walter Benjamin fellowship from the Deutsche Forschungsgemeinschaft. J.A.P. was supported by a Studienstiftung des Deutschen Volkes fellowship. This work was supported by grant R01 CA073808 (NIH) and used the MIT Structural Biology Core Facility and computational resources through allocation BIO230178 from the Advanced Cyberinfrastructure Coordination Ecosystem: Services & Support (ACCESS) program, which was supported by grants 2138259, 2138286, 2138307, 2137603, and 2138296 (NSF).

