## Supplementary Information for "Pseudouridine Residues as Substrates for Serum Ribonucleases"

#### Table of Contents

**1. General Information**
- 1.1. Abbreviations ..... S2
- 1.2. Chemical Reagents and Instrumentation ..... S2
- 1.3. Biological Reagents and Instrumentation ..... S2

**2. Synthesis** ..... S4

**3. Heterologous Production and Purification of RNase 1**
- 3.1. SDS–PAGE and LC–MS of RNase 1 ..... S14
- 3.2. Activity Validation of Commercial RNase A and Recombinant RNase 1 ..... S14

**4. Experimentally Derived Extinction Coefficients of UpA, ΨpA, and m$^1$ΨpA** ..... S17

**5. UV spectra of UpA, ΨpA, and m$^1$ΨpA** ..... S17

**6. Additional Kinetic Data for Catalysis of UpA, ΨpA, and m$^1$ΨpA Cleavage** ..... S19

**7. Kinetic Data for Catalysis of UpA Cleavage at Different pHs** ..... S20

**8. Statistics for X-Ray Crystallography** ..... S21

**9. Decavanadates in Crystal Structures** ..... S23

**10. Computational Analyses**
- 10.1. Top-Ranked Docking Poses of Enzyme·Substrate Complexes ..... S24
- 10.2. Molecular Dynamics Simulations of RNase A·Substrate Complexes ..... S25
- 10.3. Molecular Dynamics Simulations of RNase 1·Substrate Complexes ..... S25
- 10.4. RMSD Plots for Molecular Dynamics Simulations with RNase A and RNase 1 ..... S26

**11. Kinetic Data for Non-Enzymatic UpA and m$^1$ΨpA Cleavage** ..... S28

**12. NMR Spectra** ..... S30

**13. References** ..... S43

#### 1. General Information

##### 1.1. Abbreviations

| | |
|---|---|
| BCA | bicinchoninic acid |
| CHES | 2-(cyclohexylamino)ethanesulfonic acid |
| DCM | dichloromethane |
| DEPC | diethyl pyrocarbonate |
| DMF | dimethylformamide |
| EDTA | ethylenediaminetetraacetic acid |
| EtOAc | ethyl acetate |
| FRET | fluorescence resonance energy transfer |
| IPTG | isopropyl β-D-1-thiogalactopyranoside |
| LC | liquid chromatography |
| $m^1\Psi pA$ | $N^1$-methylpseudouridylyl(3′→5′)adenosine |
| $m^1\Psi$>v | $N^1$-methylpseudouridine 2′,3′-cyclic vanadate |
| MALDI | matrix-assisted laser desorption/ionization |
| MES | 2-morpholinoethanesolufonic acid |
| MS | mass spectrometry |
| MWCO | molecular weight cutoff |
| OVS | oligo(vinylsulfonic acid) |
| PAGE | polyacrylamide gel electrophoresis |
| poly(A) | poly(adenylic acid) |
| QTOF | quadrupole time-of-flight |
| ΨpA | pseudouridylyl(3′→5′)adenosine |
| Ψ>v | pseudouridine 2′,3′-cyclic vanadate |
| RNase A | bovine pancreatic ribonuclease (EC 3.1.27.5) |
| RNase 1 | human ribonuclease 1 (EC 3.1.27.5) |
| SDS | sodium dodecylsulfate |
| TB | terrific broth |
| TBSCl | *tert*-butyldimethylsilyl chloride |
| THF | tetrahydrofuran |
| TIC | total ion chromatogram |
| Tris | tris(hydroxymethyl)aminomethane |
| UpA | uridylyl(3′→5′)adenosine |
| U>v | uridine 2′,3′-cyclic vanadate |
| UV | ultraviolet |
| vis | visible |

##### 1.2. Chemical Reagents and Instrumentation

All procedures were performed at ambient temperature (~22 °C) and pressure (~1.0 atm) unless indicated otherwise. All reactions were performed in a reaction vial fitted with TFE-silicone septa under $N_2$(g) using standard Schlenk-line techniques. Reactions carried out at low temperatures were cooled by cooling agents in a Dewar vessel (water–ice bath at 0 °C).

Commercial chemicals were of reagent grade or better from Sigma–Aldrich (St. Louis, MO) and were used without further purification unless indicated otherwise. In all reactions involving anhydrous solvents, glassware was either oven- or flame-dried. Reagent-grade dichloromethane (DCM) was dried over a column of alumina and removed from a dry still under an inert atmosphere. All reactions were magnetically stirred and monitored by liquid chromatography–mass spectrometry (LC–MS) and analytical thin-layer chromatography (TLC). Purification was done with flash column chromatography performed with silica gel, typically using an Isolera One system from Biotage (Uppsala, Sweden), unless indicated otherwise. The term "concentrated under reduced pressure" refers to the removal of solvents and other volatile materials using a Buchi rotary evaporator (model R-210) at water aspirator pressure (<20 torr) while maintaining the water-bath temperature below 40 °C.

$^{1}$H, $^{13}$C, and $^{31}$P NMR spectra for compound characterization were acquired with a Bruker (Billerica, MA) Avance Neo 500 MHz spectrometer in the Department of Chemistry Instrumentation Facility (DCIF) at MIT. $^{1}$H and $^{13}$C NMR spectra were referenced to residual solvent peaks in $CDCl_3$ ($^{1}$H, 7.26 ppm; $^{13}$C, 77.16 ppm) or $D_2O$ ($^{1}$H, 4.79 ppm). $^{13}$C NMR spectra in $D_2O$ were referenced externally to TMS (0 ppm). $^{31}$P NMR spectra were referenced externally to 85% v/v $H_3PO_4$ (0 ppm). Multiplicities are abbreviated as s (singlet), d (doublet), t (triplet), q (quartet), and m (multiplet).

$^{31}$P NMR spectra for non-enzymatic cleavage kinetics were acquired with a two-channel Bruker Avance-III HD Nanobay 400 MHz spectrometer equipped with a 5 mm $N_2$(l)-cooled Prodigy broad band observe cryoprobe.

Mass spectrometry was performed with an LCT electrospray ionization (ESI) 1260 Infinity II instrument from Agilent Technologies (Santa Clara, CA) and an LC–MS column (Agilent Technologies, Poroshell 120, SB C18-reversed-phase, length 50 mm, internal diameter: 2.1 mm, particle size: 2.7 micron) with a gradient of 10−95% v/v MeCN (0.1% v/v formic acid) in water (0.1% v/v formic acid) over 10 min. High-resolution mass spectrometry (HRMS) was performed with a JEOL AccuTOF 4G LC-plus equipped with an intense DART (Direct Analysis in Real Time) source.

No unexpected or unusually high safety hazards were encountered during the reported work.

##### 1.3. Biological Reagents and Instrumentation

All procedures were performed at ambient temperature (∼22 °C) and pressure (~1.0 atm) unless indicated otherwise. RNase A (product #R6513-50MG; ≥79 Kunitz units mg$^{-1}$) was from Sigma–Aldrich. RNase 1 concentrations were determined using a DS-11 UV–vis spectrophotometer from DeNovix (Wilmington, DE) and validated using a BCA assay (Smith et al. 1985) with a kit from Thermo Fisher Scientific (product #23227). Cell lysis was done using a benchtop cell disruptor from Constant Systems (Kennesaw, GA). Dialysis tubing was from Spectrum Labs (Rancho Dominguez, CA). Contaminating OVS was removed from MES buffer by anion-exchange chromatography (Smith et al. 2003). Fluorescence and absorbance measurements for enzyme kinetics were made with a Spark plate reader from Tecan (Männedorf, Switzerland). Clear 96-well half area, UV-star (product #675801) from Greiner were used for absorbance assays. Dinucleotide extinction coefficients were assessed with the DS-11 UV–vis spectrophotometer, which uses a variable path length and automatically scales the absorbance to a path length of 1 cm in the output.

BL21(DE3) cells (product #69450-3) were from Sigma–Aldrich and were expanded to make electrocompetent cells. Crystals were seeded using a Seeding Tool (product #HR8-133) from Hampton Research.

#### 2. Synthesis

##### 2′,5′-Di(*t*-butyldimethylsilyl)uridine (1a)

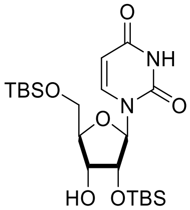


The synthetic procedure was adapted from Ogilvie and coworkers (Hakimelahi et al. 1982).

Uridine (3.00 g, 12.3 mmol) was dissolved in anhydrous THF (100 mL). Silver nitrate (4.59 g, 27.0 mmol) and TBSCl (4.07 g, 27.0 mmol) were added to the resulting solution. Then, pyridine (4.96 mL, 61.4 mmol) was added. The reaction mixture was stirred for 12 h in the dark. The reaction mixture was filtered over celite, washed with EtOAc (3 × 50 mL), and concentrated under reduced pressure. The crude product was purified by flash chromatography (5:1→1:1 hexanes/EtOAc). Product **1a** was obtained as a colorless oil (4.83 g, 10.2 mmol, 83%).

All analytical data were in accordance with the literature (Razkin et al. 2007).

##### 2′,5′-Di(*t*-butyldimethylsilyl)-pseudouridine (1b)

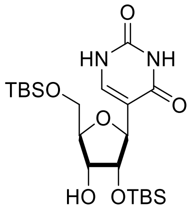


Pseudouridine (2.00 g, 8.2 mmol) was dissolved in anhydrous THF (75 mL). Silver nitrate (3.06 g, 18.0 mmol) and TBSCl (2.72 g, 18.0 mmol) were added to the resulting solution. Then, pyridine (3.31 mL, 40.9 mmol) was added. The reaction mixture was stirred for 12 h in the dark. The reaction mixture was filtered over celite and washed with EtOAc (3 × 50 mL). The reaction mixture was concentrated under reduced pressure. The crude product was purified by flash chromatography (5:1→1:1 hexanes/EtOAc). Product **1b** was obtained as a colorless oil (2.01 g, 4.3 mmol, 52%).

$^1$H NMR ($CDCl_3$, 500 MHz, $\delta$): 9.33 (m, 1H), 9.10 (m, 1H), 7.70 (dd, 1H, $J$ = 5.9, 1.2 Hz), 4.78 (dd, 1H, $J$ = 2.4, 1.2 Hz), 4.18 (dd, 1H, $J$ = 4.7, 2.4 Hz), 4.07 (ddd, 1H, $J$ = 9.0, 7.3, 4.7 Hz), 3.99 (m, 1H), 3.86 (dt, 1H, $J$ = 7.3, 2.6 Hz), 3.80 (dd, 1H, $J$ = 11.5, 2.6 Hz), 2.42 (d, 1H, $J$ = 9.0 Hz), 0.93 (s, 9H), 0.91 (s, 9H), 0.22 (s, 3H), 0.14 (s, 3H), 0.11 (s, 3H), 0.08 (s, 3H). $^{13}$C NMR ($CDCl_3$, 126 MHz, $\delta$): 162.6, 152.1, 138.8, 113.5, 83.1, 79.3, 76.5, 70.1, 62.2, 26.1, 26.0, 18.6, 18.2, 0.1, −4.23, −5.2, −5.22, −5.30.

**2′,5′-Di(*t*-butyldimethylsilyl)-*N*[1]-methylpseudouridine (1c)**

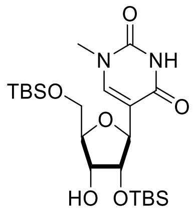


*N*[1]-Methylpseudouridine (2.00 g, 7.8 mmol) was dissolved in anhydrous THF (75 mL). Silver nitrate (2.57 g, 17.0 mmol) and TBSCl (2.89 g, 17.0 mmol) were added to the resulting solution. Then, pyridine (3.93 mL, 38.7 mmol) was added. The reaction mixture was stirred for 12 h in the dark. The reaction mixture was filtered over celite and washed with EtOAc (3 × 50 mL). The reaction mixture was concentrated under reduced pressure. The crude product was purified by flash chromatography (5:1→1:1 hexanes/EtOAc). Product **1c** was obtained as a colorless oil (1.36 g, 2.8 mmol, 36%).

$^1$H NMR ($CDCl_3$, 500 MHz, $\delta$): 8.82 (s, 1H), 7.52 (d, 1H, $J$ = 1.2 Hz), 4.79 (dd, 1H, $J$ = 2.5, 1.2 Hz), 4.16 (dd, 1H, $J$ = 4.8, 2.5 Hz), 4.07–3.99 (m, 2H), 3.87 (dt, 1H, $J$ = 7.3, 2.4 Hz), 3.81 (dd, 1H, $J$ = 11.6, 2.7 Hz), 3.35 (s, 3H), 2.44 (d, 1H, $J$ = 8.9 Hz,), 0.93 (s, 9H), 0.92 (s, 9H), 0.23 (s, 3H), 0.14 (s, 3H), 0.12 (s, 3H), 0.11 (s, 3H). $^{13}$C NMR ($CDCl_3$, 126 MHz, $\delta$): 162.4, 150.9, 142.9, 113.2, 82.9, 79.1, 70.0, 62.1, 36.3, 26.1, 25.9, 18.7, 18.1, −4.3, −5.19, −5.22, −5.27.

**2′,3′,5′-Tri(*t*-butyldimethylsilyl)adenosine (4)**

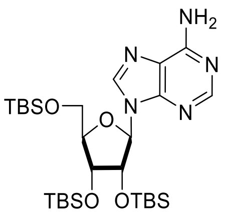


Adenosine (3.00 g, 11.1 mmol) was suspended in DMF (20 mL). Imidazole (6.83 g, 100.3 mmol) and TBSCl (7.56 g, 50.1 mmol) were added, and the reaction mixture was stirred overnight. The reaction was stopped by the addition of $H_2O$ (50 mL). The reaction mixture was extracted with EtOAc (3 × 100 mL). The combined organic phases were washed with 0.5 M hydrochloric acid (3 × 30 mL), saturated sodium bicarbonate solution (3 × 30 mL), and brine (40 mL). The organic phases were dried over $Na_2SO_4$(s) and concentrated under reduced pressure. The crude product was purified by flash chromatography (3:1 hexanes/EtOAc→EtOAc). Product **4** was obtained as a colorless solid (5.86 g, 9.57 mmol, 86%).

All analytical data were in accordance with the literature (Hisamatsu et al. 2006).

**2′,3′-Di(*t*-butyldimethylsilyl)adenosine (3)**

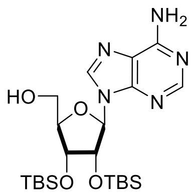


Compound **4** (3.00 g, 4.9 mmol) was dissolved in aqueous acetic acid (50 mL, 80% v/v), and the resulting solution was stirred at 100 °C for 3 h. The reaction mixture was concentrated under reduced pressure to an oil and purified by flash chromatography (3:1 hexanes/EtOAc→EtOAc). Product **3** was obtained as a colorless solid (1.92 g, 3.87 mmol, 79%).

All analytical data were in accordance with the literature (Kotch et al. 2003).

**Uridine Phosphoramidite (2a)**

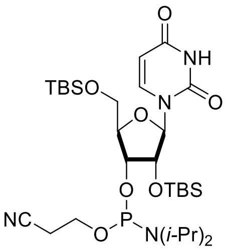


Compound **1a** (800 mg, 1.7 mmol) was dissolved in THF (5 mL). 2-Cyanoethyl *N*,*N*,*N′*,*N′*-tetraisopropylphosphorodiamidite (537 μL, 1.7 mmol) was added. Tetrazol (3.42 mL, 450 mM in MeCN, 1.5 mmol) was added and the reaction was stirred overnight at 40 °C. The reaction was stopped by the addition of a saturated aqueous solution of $NaHCO_3$ (5 mL). The mixture was extracted with EtOAc (3 × 10 mL). The combined organic phases were dried over $Na_2SO_4$(s) and concentrated under reduced pressure. The crude product was purified by flash chromatography (3:1 hexanes/EtOAc→EtOAc). Product **2a** was obtained as a colorless oil that was a mixture of diastereomers (883 mg, 1.31 mmol, 85%).

$^{31}$P NMR ($CDCl_3$, 202 MHz, $\delta$): 150.9, 149.9. HRMS–ESI (*m*/*z*): [M + $H^+$] calcd for $C_{24}H_{45}N_3O_8PSi_2$, 590.2477; found, 590.2485 (hydrolyzed).

**Pseudouridine Phosphoramidite (1b)**

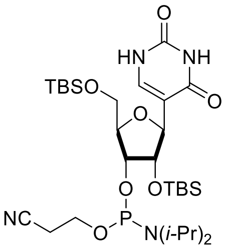


Compound **1b** (330 mg, 698 μmol) was dissolved in THF (4 mL). 2-Cyanoethyl *N*,*N*,*N′*,*N′*-tetraisopropylphosphorodiamidite (221 μL, 698 μmol) was added. Then, tetrazol (1.4 mL, 450 mM in MeCN, 634 µmol) was added and the reaction was stirred overnight at 40 °C. The reaction was stopped by the addition of a saturated aqueous solution of $NaHCO_3$ (5 mL). The mixture was extracted with EtOAc (3 × 10 mL). The combined organic phases were dried over $Na_2SO_4$(s) and concentrated under reduced pressure. The crude product was purified by flash chromatography (EtOAc→3:1 hexanes/EtOAc). Product **2b** was obtained as a colorless oil that was a mixture of diastereomers (273 mg, 406 µmol, 64%). Due to its limited stability, the product was characterized only by $^{31}$P NMR and used immediately in the next synthetic step.

$^{31}$P NMR ($CDCl_3$, 202 MHz, *δ*): 149.9, 148.6. HRMS–ESI (*m*/*z*): [M + $H^+$] calcd for $C_{24}H_{45}N_3O_8PSi_2$, 590.2477; found, 590.2466 (hydrolyzed).

##### ***N*****[1]-Methylpseudouridine Phosphoramidite (2c)**

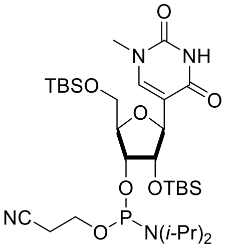


Compound **1c** (650 mg, 1.3 mmol) was dissolved in THF (7 mL). 2-Cyanoethyl *N*,*N*,*N'*,*N'*-tetraisopropylphosphorodiamidite (428 µL, 1.34 mmol) was added to the resulting solution. Then, tetrazol (2.7 mL, 450 mM in MeCN, 1.2 mmol) was added. The reaction mixture was stirred overnight at 40 °C. The reaction was stopped by the addition of a saturated aqueous solution of $NaHCO_3$ (8 mL). The mixture was extracted with EtOAc (3 × 15 mL). The combined organic phases were dried over $Na_2SO_4$(s) and concentrated under reduced pressure. The crude product was purified by flash chromatography (3:1 hexanes/EtOAc→EtOAc). Product **2c** was obtained as a colorless oil that was a mixture of diastereomers (558 mg, 812 µmol, 67%). Do to its limited stability, the product was characterized only by $^{31}$P NMR and used immediately in the next synthetic step.

$^{31}$P NMR ($CDCl_3$, 202 MHz, *δ*): 149.7, 148.9. HRMS–ESI (*m*/*z*): [M + $H^+$] calcd for $C_{31}H_{60}N_4O_7PSi_2$, 687.3733; found, 687.3741.

**Protected Uridylyl(3′→5′)adenosine (5a)**

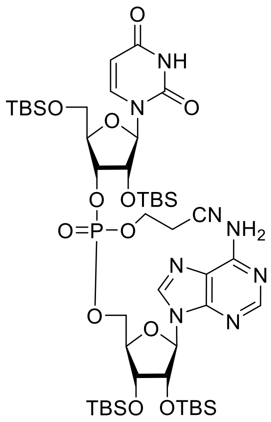


Phosphoramidite **2a** (100 mg, 148 μmol) and compound **3** (88 mg, 178 μmol) were dissolved in THF (2 mL). Tetrazole (495 μL, 450 mM in MeCN, 223 μmol) was added to the resulting solution, and the reaction mixture was stirred at 40 °C overnight. The reaction mixture was cooled to room temperature. $I_2$ (38 mg, 148 μmol) was dissolved in 2:1 THF/$H_2O$ (2 mL), and the resulting solution was added to the reaction mixture. The reaction mixture was stirred for 15 min at room temperature. The reaction was stopped by the addition of an aqueous solution of sodium bisulfite (1 mL, 10% w/v). The reaction mixture was extracted with DCM (3 × 4 mL) The combined organic phases were dried over $Na_2SO_4$(s) and concentrated under reduced pressure. The crude product was purified by flash chromatography (DCM→9:1 DCM/methanol). Product **5a** was obtained as a colorless oil that was a mixture of diastereomers (76 mg, 70 μmol, 47%).

$^1$H NMR ($CDCl_3$, 500 MHz, $\delta$): 10.24 (s), 8.86 (s), 8.44 (s), 8.41 (s), 8.07 (d, $J$ = 1.5 Hz), 7.79 (d, $J$ = 8.1 Hz), 7.68 (d, $J$ = 8.2 Hz), 7.52 (m), 6.95 (s), 6.90 (s), 6.12 (d, $J$ = 7 Hz), 5.96 (d, $J$ = 7.0 Hz), 5.89 (m,), 5.84 (m), 5.69 (m), 5.00 (m), 4.81 (m), 4.75 (m), 4.70–4.66 (m), 4.51–4.08 (m), 4.07–3.96 (m), 3.88 (m), 3.81 (m), 3.66–3.49 (m), 2.80–2.63 (m), 0.99–0.75 (m), 0.17–−0.04 (m). $^{13}$C NMR ($CDCl_3$, 126 MHz, $\delta$): 164.1, 162.9, 156.1, 153.0, 151.3, 151.0, 149.3, 142.9, 119.8, 116.3, 113.3, 102.92, 90.4, 86.3, 83.0, 82.1, 79.0, 74.3, 70.1, 62.9, 62.2, 36.2, 26.0, 26.0, 25.9, 25.8, 25.5, 18.6, 18.3, 18.1, 18.0, 17.9, 17.9, −4.2, −4.3, −4.5, −4.6, −4.9, −5.1, −5.2, −5.4, −5.6. $^{31}$P NMR ($CDCl_3$, 202 MHz, $\delta$): −1.64, −1.81. HRMS–ESI ($m/z$): [M + $H^+$] calcd for $C_{46}H_{84}N_8O_{12}PSi_4$, 1083.5018; found, 1083.5012.

**Protected Pseudouridylyl(3′→5′)adenosine (5b)**

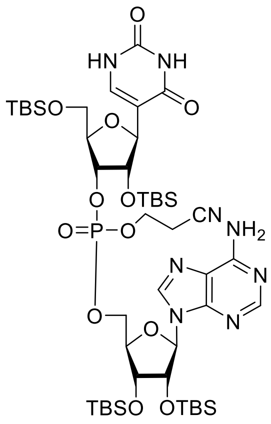


Phosphoramidite **2b** (250 mg, 372 µmol) and **4** (221 mg, 445 µmol) were dissolved in THF (2 mL). Tetrazole (1.2 mL, 450 mM in MeCN, 540 µmol) was added to the resulting solution, and the reaction mixture was stirred at 40 °C overnight. The reaction mixture was cooled to room temperature. $I_2$ (94 mg, 371 µmol) was dissolved in 2:1 THF/$H_2O$ (4 mL), and the resulting solution was added to the reaction mixture. The reaction mixture was stirred for 15 min at room temperature. The reaction was stopped by the addition of an aqueous solution of sodium bisulfite (2 mL, 10% w/v). The reaction mixture was extracted with DCM (3 × 8 mL). The combined organic phases were dried over $Na_2SO_4$(s) and concentrated under reduced pressure. The crude product was purified by flash chromatography (DCM→9:1 DCM:methanol). Product **5b** was obtained as a colorless oil that was a mixture of diastereomers (209 mg, 193 µmol, 52%).

$^1$H NMR ($CDCl_3$, 500 MHz, $\delta$): 9.23 (s), 8.63 (s), 8.39–8.29 (m), 8.21–7.77 (m), 7.59–7.31 (m), 5.87 (m), 5.73 (s), 5.21 (d), 5.02 (s, 2H), 4.87–4.16 (m), 3.58 (m), 3.22 (d, $J$ = 11.2 Hz, 1H), 3.11–2.90 (m), 2.85–2.61 (m), 0.98–0.63 (m), 0.24––0.15 (m). $^{13}$C NMR ($CDCl_3$, 126 MHz, $\delta$): 163.5, 156.0, 152.6, 152.3, 151.6, 149.5, 149.4, 143.9, 142.9, 141.5, 140.7, 139.9, 120.6, 120.2, 116.7, 116.3, 113.5, 112.5, 90.8, 89.8, 82.9, 82.3, 79.4, 74.1, 72.2, 71.3, 67.1, 62.8, 62.5, 62.2, 53.5, 36.0, 26.1, 25.9, 25.9, 25.8, 25.8, 25.7, 25.7, 19.6, 19.6, 18.50, 18.2, 18.2, 18.1, 18.1, 17.9, −4.2, −4.3, −4.5, −4.6, −4.6, −4.6, −4.7, −4.8, −4.8, −4.9, −5.1, −5.2, −5.3, −5.5. $^{31}$P NMR ($CDCl_3$, 202 MHz, $\delta$): −2.40. HRMS–ESI ($m/z$): [M + $H^+$] calcd for $C_{46}H_{84}N_8O_{12}PSi_4$, 1083.5018; found, 1083.5004.

**Protected $N^1$-Methylpseudouridylyl(3′→5′)adenosine (5c)**

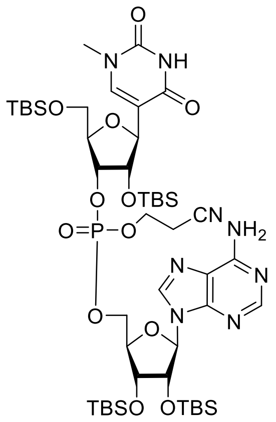

Phosphoramidite **2c** (200 mg, 291 µmol) and **4** (173 mg, 349 µmol) were dissolved in THF (2 mL). Tetrazole (970 µL, 450 mM in MeCN, 437 µmol) was added to the resulting solution, and the reaction mixture was stirred overnight at 40 °C. The reaction was cooled to room temperature. $I_2$ (74 mg, 291 µmol) was dissolved in 2:1 THF/$H_2O$ (4 mL), and the resulting solution was added to the reaction mixture. The reaction mixture was stirred for 15 min at room temperature. The reaction was stopped by the addition of an aqueous solution of sodium bisulfite (2 mL, 10% w/v). The reaction mixture was extracted with DCM (3 × 8 mL) The combined organic phases were dried over $Na_2SO_4$(s) and concentrated under reduced pressure. The crude product was purified by flash chromatography (DCM→9:1 DCM/methanol). Product **5c** was obtained as a colorless oil that was a mixture of diastereomers (181 mg, 165 µmol, 57%).

$^1$H NMR ($CDCl_3$, 500 MHz, $\delta$): 9.24 (s), 8.37 (s), 8.01 (s), 7.96 (s), 7.37 (s), 7.26 (s), 7.23 (s), 6.91 (s), 5.89 (d, $J$ = 4.6 Hz), 5.77 (d, $J$ = 3.7 Hz), 4.97 (m), 4.83 (d), 4.75 (s, 2H), 4.69 (m), 4.62 (d, $J$ = 5.5 Hz), 4.57 (d, $J$ = 7.8 Hz), 4.49–4.19 (m), 3.85 (m), 3.78 (s,), 3.69 (s), 3.43 (m), 3.38 (s,), 2.89 (m), 2.70 (m), 0.95–0.89 (m), 0.87–0.78 (mm), 0.08 (m), −0.01 (m). $^{13}$C NMR ($CDCl_3$, 126 MHz, $\delta$): 163.1, 156.3, 155.8, 152.9, 149.0, 144.2, 135.3, 116.6, 91.9, 83.3, 79.7, 74.2, 73.6, 72.2, 70.5, 62.4, 62.0, 60.4, 53.4, 47.9, 26.0, 25.8, 25.8, 25.7, 19.5, 19.4, 19.1, 18.2, 18.0, 17.9, 17.9, 1.0, −4.4, −4.7, −5.0, −5.1, −5.6. $^{31}$P NMR ($CDCl_3$, 202 MHz, $\delta$): −1.94, −2.12. HRMS–ESI ($m/z$): [M + $H^+$] calcd for $C_{47}H_{86}N_8O_{12}PSi_4$, 1097.5174; found, 1097.5182.

**Uridylyl(3′→5′)adenosine·NEt₃**

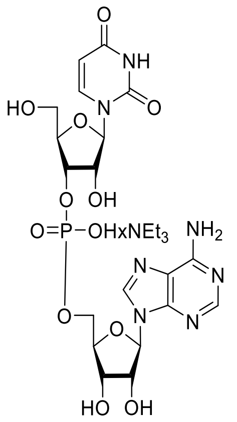

Compound **5a** (20 mg, 18 μmol) was dissolved in in 2:1 ammonium hydroxide/ethanol (2 mL), and the resulting solution was stirred at 55 °C overnight. The reaction mixture was concentrated under reduced pressure. The resulting oil was taken up in THF (1 mL), TBAF (369 μL, 369 μmol, 1 M in THF) was added, and the resulting mixture was stirred for 48 h at room temperature. The reaction mixture was diluted twofold with $H_2O$, the THF was removed under reduced pressure, and the aqueous solution was extracted with diethyl ether (1 × 3 mL). Any ether dissolved in the aqueous layer was removed under reduced pressure. A HiTrap Q HP anion-exchange column was charged with the nucleotide, which was eluted with a linear gradient of 0–40% v/v triethylammonium bicarbonate buffer, pH 8. UpA eluted soon after the beginning of the gradient. The fractions containing UpA were concentrated under reduced pressure to an oil. The oil was dissolved in a minimal amount of MeOH, and EtOAc was added to precipitate the dinucleotide. The solution was stored at −20 °C overnight to complete precipitation. The precipitate was dried, and lyophilization yielded the triethylammonium salt of **UpA** as a fluffy white solid (4.5 mg, 7.9 μmol, 43%).

$^1$H NMR ($D_2O$, 400 MHz, $\delta$): 8.44 (s, 1H), 8.26 (s, 1H), 7.76 (d, 1H, $J$ = 8.1 Hz,), 6.12 (d, 1H, $J$ = 4.7 Hz), 5.78 (d, 1H, $J$ = 8.1 Hz), 5.73 (d, 1H, $J$ = 4.5 Hz), 4.72 (pt, 1H, $J$ = 4.9 Hz,), 4.54 (pt, 1H, $J$ = 5.1 Hz,), 4.47 (m, 1H), 4.37 (m, 1H), 4.31–4.21 (m, 2H), 4.17 (m, 3H), 3.79 (dd, 1H, $J$ = 12.9, 2.8 Hz), 3.73 (dd, 1H, $J$ = 12.9, 3.9 Hz), 3.21 (q, 6H, $J$ = 7.3 Hz), 1.37–1.13 (t, 9H, $J$ = 7.3 Hz). $^{13}$C NMR ($D_2O$, 101 MHz, $\delta$): 152.6, 146.5, 141.3, 139.6, 102.1, 89.2, 87.4, 74.0, 72.9, 69.8, 60.2, 46.7, 8.2. $^{31}$P NMR ($D_2O$, 162 MHz, $\delta$): −0.73. HRMS–ESI ($m/z$): [M − $H^+$] calcd for $C_{19}H_{23}N_7O_{12}P$, 572.1148; found, 572.1140.

**Pseudouridylyl(3′→5′)adenosine·NEt$_3$**

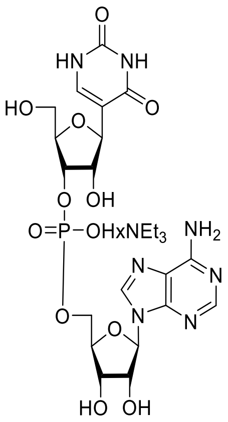

Compound **5b** (30 mg, 28 µmol) was dissolved in in 2:1 ammonium hydroxide/ethanol (2 mL), and the resulting solution was stirred at 55 °C overnight. The reaction mixture was concentrated under reduced pressure. The resulting oil was taken up in THF (1 mL), and TBAF (414 µL, 414 µmol, 1 M in THF) was added, and the resulting mixture was stirred for 48 h at room temperature. The reaction mixture was diluted twofold with $H_2O$, the THF was removed under reduced pressure, and the aqueous solution was extracted with diethylether (1 × 3 mL). Any ether dissolved in the aqueous layer was removed under reduced pressure. A HiTrap Q HP anion-exchange column was charged with the nucleotide, which was eluted with a linear gradient of 0–40% v/v triethylammonium bicarbonate buffer, pH 8. ΨpA eluted soon after the beginning of the gradient. The fractions containing ΨpA were concentrated under reduced pressure to an oil. The oil was dissolved in a minimal amount of MeOH, and EtOAc was added to precipitate the dinucleotide. The solution was stored at −20 °C overnight to complete precipitation. The precipitate was dried, and lyophilization yielded the triethylammonium salt of **ΨpA** as a fluffy white solid (7.45 mg, 13.0 µmol, 47%).

$^1$H NMR ($D_2O$, 500 MHz, $\delta$): 8.48 (s, 1H), 8.32 (s, 1H), 7.55 (s, 1H), 6.13 (d, 1H, $J$ = 4.3 Hz,), 4.64 (pt, 1H, $J$ = 4.7 Hz,), 4.59 (d, 1H, $J$ = 4.7 Hz), 4.51 (pt, 1H, $J$ = 5.1 Hz), 4.40–4.36 (m, 1H), 4.36–4.32 (m, 1H), 4.26 (m, 1H), 4.22 (m, 1H), 4.13 (m, 1H), 4.09 (s, 1H), 3.79 (dd, 1H, $J$ = 12.8, 2.8 Hz), 3.68 (dd, 1H, $J$ = 12.8, 4.6 Hz), 3.18 (q, 6H, $J$ = 7.3 Hz), 1.25 (td, 9H, $J$ = 7.3) $^{13}$C NMR ($D_2O$, 126 MHz, $\delta$): 149.6, 148.5, 140.9, 140.5, 110.2, 87.9, 83.3, 81.8, 79.4, 74.4, 72.5, 69.6, 64.3, 60.9, 46.7, 8.2. $^{31}$P NMR ($D_2O$, 202 MHz, $\delta$): 0.08. HRMS–ESI ($m/z$): [M − H$^+$] calcd for $C_{19}H_{23}N_7O_{12}P$, 572.1148; found, 572.1157.

##### *N*[1]-Methylpseudouridylyl(3′→5′)adenosine (m$^1$ΨpA)

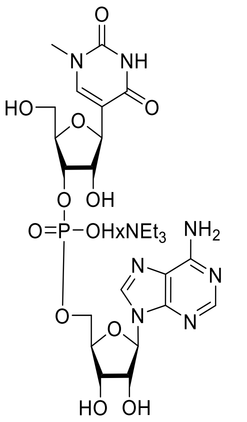


Compound **5c** (50 mg, 46 µmol) was dissolved in 2:1 ammonium hydroxide/ethanol (3 mL), and the resulting solution was stirred at 55 °C overnight. The reaction mixture was concentrated under reduced pressure. The resulting oil was taken up in THF (1.5 mL), and TBAF (682 µL, 682 µmol, 1 M in THF) was added. The resulting mixture was stirred for 48 h at room temperature. The reaction mixture was diluted twofold with $H_2O$, the THF was removed under reduced pressure, and the aqueous solution was extracted with diethylether (1 × 3 mL). Any ether dissolved in the aqueous layer was removed under reduced pressure. A HiTrap Q HP anion-exchange column was charged with the nucleotide and eluted with a linear gradient from 0–40% v/v triethylammonium bicarbonate buffer, pH 8. m$^1$ΨpA eluted soon after the beginning of the gradient. The fractions containing m$^1$ΨpA were concentrated under reduced pressure to an oil. The oil was dissolved in a minimal amount of MeOH, and EtOAc was added to precipitate the dinucleotide. The solution was stored at −20 °C overnight to complete the precipitation. The precipitate was dried and lyophilization yielded the triethylammonium salt of **m$^1$ΨpA** as a fluffy white solid (11 mg, 19 µmol, 41%).

$^1$H NMR ($D_2O$, 500 MHz, *δ*): 8.47 (s, 1H,), 8.30 (s, 1H), 7.66 (s, 1H), 6.13 (d, 1H, *J* = 4.3 Hz), 4.64 (pt, 1H, *J* = 4.7 Hz), 4.61 (d, 1H, *J* = 4.6 Hz), 4.52 (pt, 1H, *J* = 5.2 Hz), 4.42 (m, 1H), 4.36 (m, 1H), 4.31–4.27 (m, 1H), 4.25 (t, 1H, *J* = 4.9 Hz), 4.15 (dt, 1H, *J* = 11.8, 3.6 Hz), 4.13–4.08 (m, 1H), 3.82 (dd, 1H, *J* = 12.8, 2.8 Hz), 3.71 (dd, 1H, *J* = 12.8, 4.5 Hz), 3.33 (s, 3H), 3.20 (q, 6H, *J* = 7.3 Hz), 1.28 (t, 9H, *J* = 7.3 Hz). $^{13}$C NMR ($D_2O$, 126 MHz, *δ*): 164.3, 153.8, 152.0, 150.4, 148.5, 145.6, 140.1, 118.6, 110.3, 87.9, 83.0, 81.8, 79.3, 74.4, 74.0, 72.6, 69.4, 64.2, 60.8, 46.6, 45.1, 36.0, 8.2. $^{31}$P NMR ($D_2O$, 202 MHz, *δ*): −0.48. HRMS–ESI (*m*/*z*): [M − $H^+$] calcd for $C_{20}H_{25}N_7O_{12}P$, 586.1304; found, 586.1315.

#### 3. Heterologous Production and Purification of RNase 1

##### 3.1. SDS–PAGE and LC–MS of RNase 1

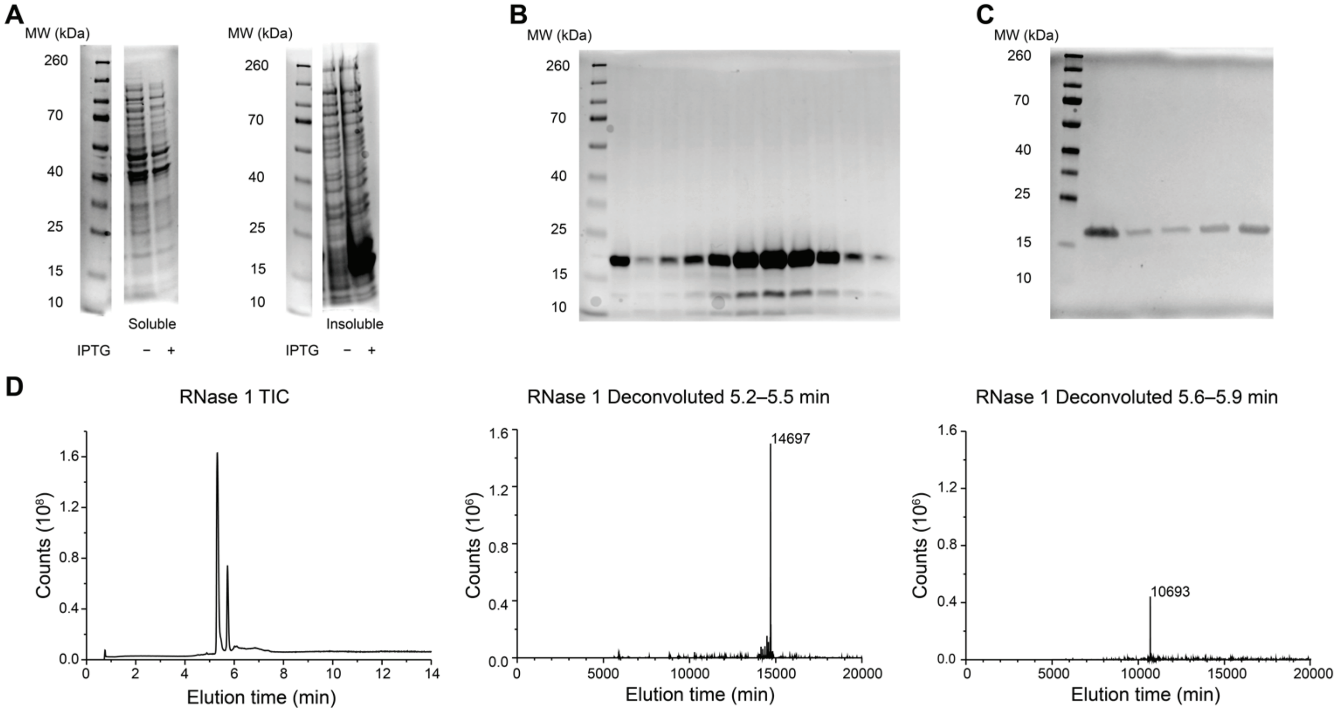


**FIGURE S1.** (*A*) SDS–PAGE gel for the heterologous production of human RNase 1. Human RNase 1 is present in the insoluble fraction after 3 h of induction with IPTG. (*B*) SDS–PAGE gel of protein-containing fractions from cation-exchange chromatography. (*C*) Gel of subsequent buffer exchange and concentration, which led to the loss of lower molecular weight species. (*D*) QTOF–MS of RNase 1 used for assays of enzymatic activity. RNase 1 was present at the expected molecular weight (14,697 Da) with a minor contaminant at 10,093 Da that was not apparent from SDS–PAGE analysis.

##### 3.2. Activity Validation of Commercial RNase A and Recombinant RNase 1

An assay with a hypersensitive FRET substrate, FAM–dArUdAdA–6-TAMRA (Fig. S2), was used to ensure that the purified RNase 1 and commercial RNase A were active catalysts of RNA cleavage (Kelemen et al. 1999; Wralstad and Raines 2024). Assays were carried out in the same buffers as those of dinucleotide cleavage (RNase A: 0.10 M OVS-free MES–NaOH, pH 6.0, containing 0.10 M NaCl; RNase 1: 0.10 M Tris–HCl, pH 7.5, 0.10 M NaCl) with 12.5 pM of enzyme and 200 nM of FAM–dArUdAdA–6-TAMRA. Fluorescence intensity ($I$) was measured with a Tecan Spark microplate reader, monitoring emission at $\lambda_{em}$ = 515 nm after excitation at $\lambda_{ex}$ = 493 nm with a 5-nm bandwidth. Assays were performed with 5 replicates in a flat, black, half-area 96-well plate. Assuming a linear relationship for substrate conversion to product and in the initial rates region of Michaelis–Menten kinetics and that assays are performed at a substrate concentration well below the $K_M$, which we estimated to be 22 μM for RNase A (Kelemen et al. 1999), the following relationship was used to determine catalytic efficiency via linear regression:

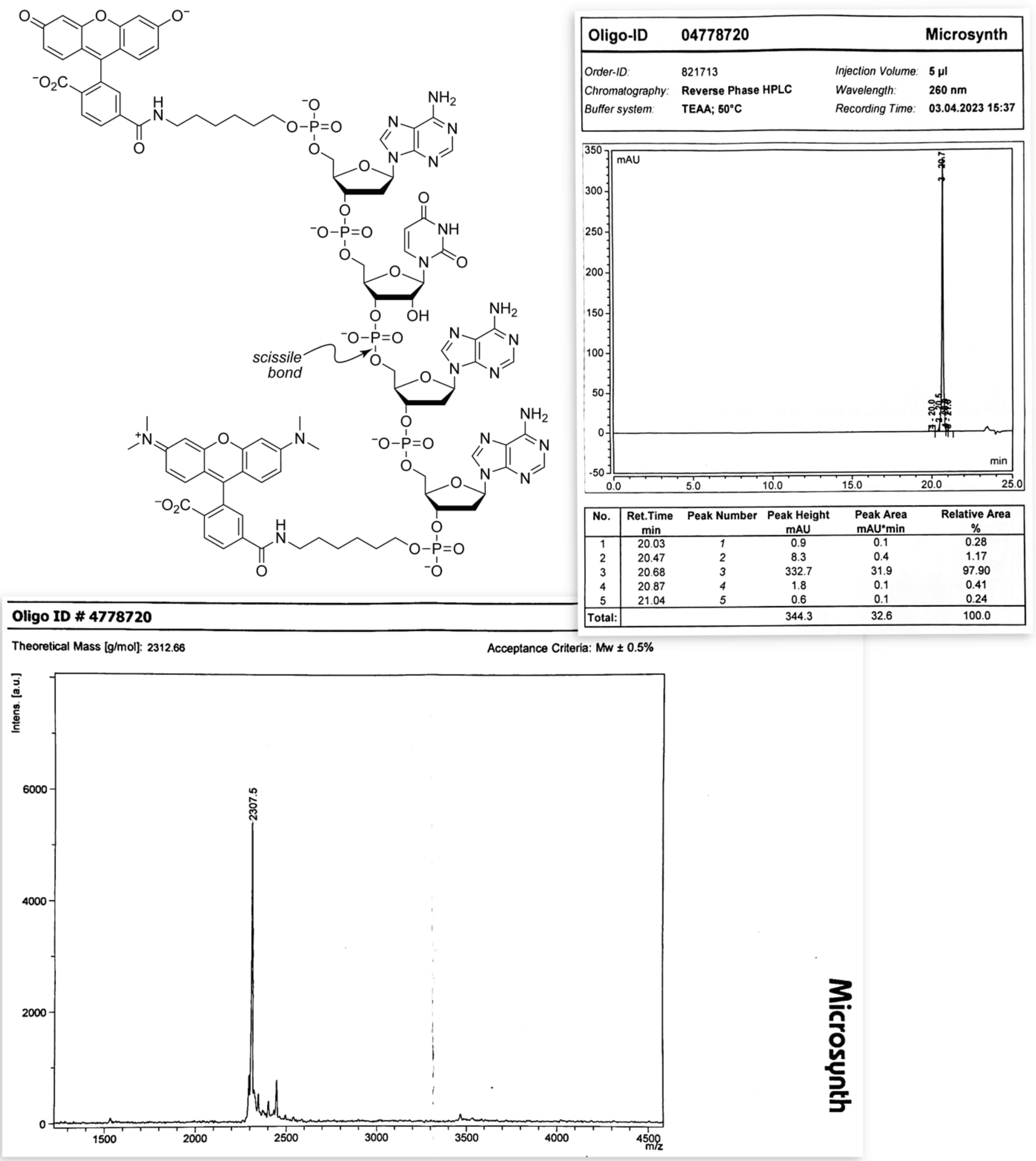


| No. | Ret.Time min | Peak Number | Peak Height mAU | Peak Area mAU*min | Relative Area % |
|---|---|---|---|---|---|
| 1 | 20.03 | 1 | 0.9 | 0.1 | 0.28 |
| 2 | 20.47 | 2 | 8.3 | 0.4 | 1.17 |
| 3 | 20.68 | 3 | 332.7 | 31.9 | 97.90 |
| 4 | 20.87 | 4 | 1.8 | 0.1 | 0.41 |
| 5 | 21.04 | 5 | 0.6 | 0.1 | 0.24 |
| Total: | | | 344.3 | 32.6 | 100.0 |

**FIGURE S2.** Structure of 6-FAM–dArU(dA)$_2$–6-TAMRA, which is a hypersensitive fluorogenic substrate for assays of ribonucleolytic activity (Kelemen et al. 1999; Wralstad and Raines 2024). The cleavage of the scissile bond increases the fluorescence of the FAM moiety with $\lambda_{ex}$ = 493 ± 5 nm and $\lambda_{em}$ = 515 ± 5 nm. Insets: reversed-phase HPLC chromatogram (top) and MALDI–TOF mass spectrum (bottom) of synthetic 6-FAM–dArU(dA)$_2$–6-TAMRA.

$$\frac{(I-I_0)}{(I_{\max}-I_0)} = \frac{[\text{product}]_t}{[\text{substrate}]_0} = \frac{k_{\text{cat}}}{K_{\text{M}}}[\text{enzyme}]t \quad (1)$$

$I_0$ was determined by measuring the fluorescence intensity before the addition of a ribonuclease, and $I_{\max}$ was determined by measuring the fluorescence intensity after the addition of excess RNase A (5 μM). The resulting values of $k_{\text{cat}}/K_{\text{M}}$ are listed in Table S1 along with literature values using a similar substrate.

**TABLE S1.** Values of $k_{\text{cat}}/K_{\text{M}}$ for the cleavage of FAM–dArUdAdA–6-TAMRA (Fig. S2) by RNase A and RNase 1.

| Ribonuclease | pH | $k_{\text{cat}}/K_{\text{M}}$ ($10^6$ $M^{-1}$ $s^{-1}$) | Literature $k_{\text{cat}}/K_{\text{M}}$ ($10^7$ $M^{-1}$ $s^{-1}$) |
|---|---|---|---|
| RNase A | 6.0 | 19.5 ± 2.7 | 25 ± 3 (Kelemen et al. 1999) |
| | 7.5 | 2.28 ± 0.18 | — |
| RNase 1 | 6.0 | 0.65 ± 0.05 | — |
| | 7.5 | 5.11 ± 0.51 | 9.5 ± 0.3 (Sayers et al. 2021) |

#### 4. Experimentally Derived Extinction Coefficients of UpA, ΨpA, and m$^1$ΨpA

Extinction coefficients were derived to determine the concentration of solutions of synthetic UpA, ΨpA, and m$^1$ΨpA. In short, >10 mg of each dinucleotide was weighed out on an analytical balance and dissolved in water to a concentration of 2 mg/mL. This solution was then diluted 1:20 into water, and the absorbance was measured at 260 nm using 1 μL volumes ($n$ = 5 technical replicates). Extinction coefficients were calculated using the Beer–Lambert law: $A_{260\ \mathrm{nm}} = \varepsilon_{260\ \mathrm{nm}} \cdot l \cdot c$. where $A_{260\ \mathrm{nm}}$ is the absorbance at 260 nm, $\varepsilon_{260\ \mathrm{nm}}$ is the extinction coefficient at 260 nm, $l$ is the path length, and $c$ is the concentration of dinucleotide. The resulting values of $\varepsilon_{260\ \mathrm{nm}}$ were UpA, 19770 ± 750 cm$^{-1}$ M$^{-1}$; ΨpA, 17890 ± 440; and m$^1$ΨpA, 17910 ± 200. The higher value for UpA is consistent with the known hypochromicity of Ψ and m$^1$Ψ (Finol et al. 2024). The value of $\varepsilon_{260\ \mathrm{nm}}$ for UpA was 25% less than that in an early report (Beaven et al. 1955).

We also evaluated how the overall spectra in the UV range differed for each substrate. These were obtained using 800 μM of substrate in 100 mM MES–NaOH buffer, pH 6.0, containing NaCl (100 mM) or 100 mM Tris–HCl buffer, pH 7.5, containing NaCl (100 mM) with absorbance scanned over 220 nm to 320 nm. In general, the spectrum of each substrate is similar, though that of m$^1$ΨpA is shifted slightly toward longer wavelengths, as shown in Fig. S3. There were no noticeable differences due to buffer or pH differences.

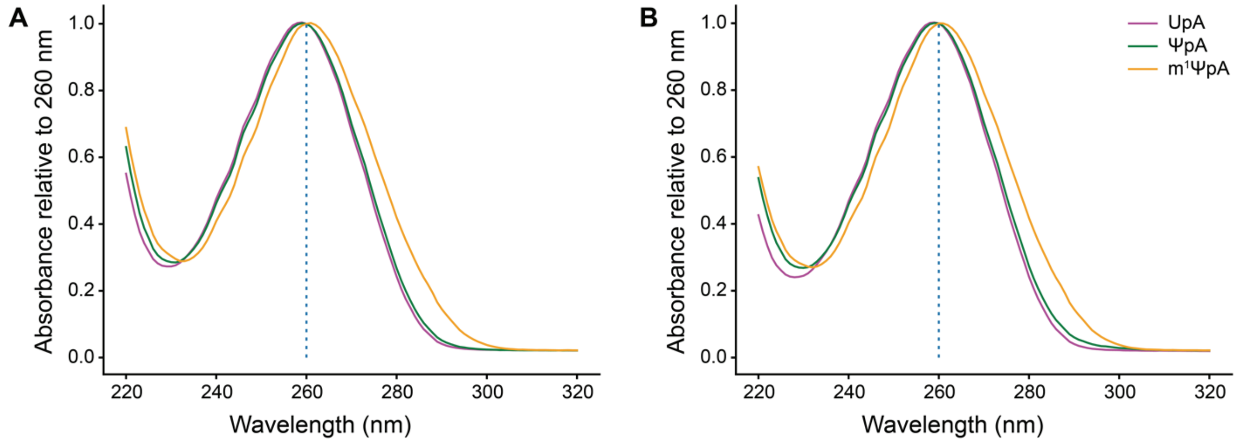


**FIGURE S3.** UV spectra of UpA, ΨpA, and m$^1$ΨpA at pH 6.0 (*A*) and pH 7.5 (*B*). Spectra are shown as the mean ($n$ = 4 technical replicates) normalized to the absorbance at 260 nm.

#### 5. UV Spectra of UpA, ΨpA, and m1 ΨpA

Previous reports on what wavelength to use for UpA cleavage assays vary. For each substrate and buffer condition, we obtained UV spectra and examined changes in absorbance after cleavage to select a wavelength that had the highest sensitivity. These examinations were done prior to kinetic assays, where a scan of absorbances between 220 and 320 nm with 1 nm increments was performed before and after the addition of enzyme. The spectra of the scan of the initial substrate (no enzyme added) and the scan of the enzyme-saturated condition (with 2 μM RNase A) were then averaged for each concentration of substrate, and the difference was subtracted to obtain the change in absorbance upon enzyme-catalyzed cleavage.

We sought a wavelength that had a large change in absorbance and was distal from the absorbance maximum, such that the absorbance change was directly proportional to substrate concentration, particularly at higher concentrations (Fig. S4). For assays in different buffers, we used the wavelength chosen for the enzyme–substrate pair at the optimal pH for catalysis. The wavelengths thus chosen for the enzyme-substrate pairs are listed in Table S2.

**TABLE S2.** Wavelength for kinetic assays of each enzyme–substrate pair

| Substrate | RNase A | RNase 1 |
|---|---|---|
| UpA | 278 nm | 276 nm |
| ΨpA | 276 nm | 276 nm |
| m$^1$ΨpA | 287 nm | 287 nm |

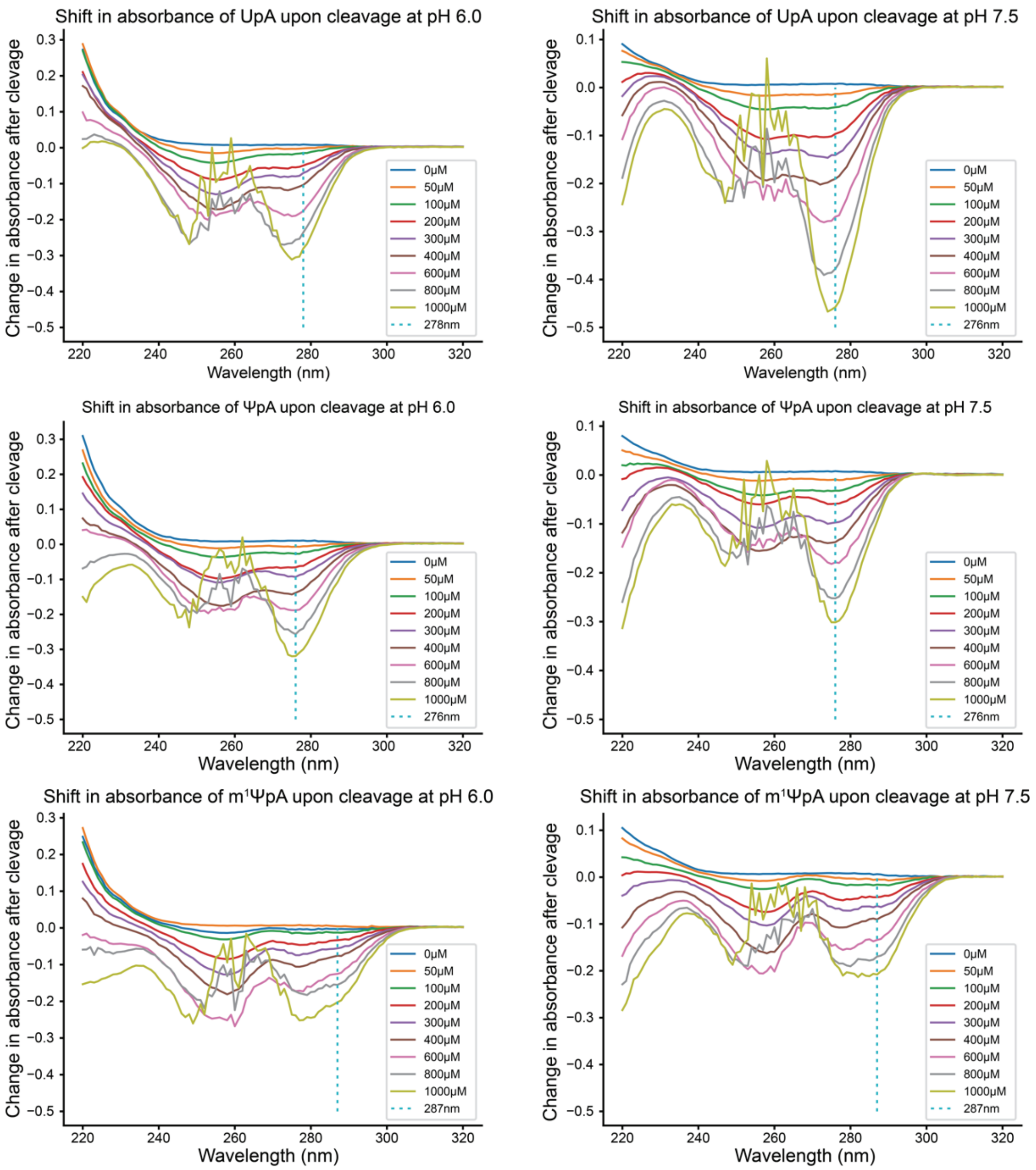


**FIGURE S4.** Shift in absorbance due to the cleavage of UpA, ΨpA, and m$^1$ΨpA at pH 6.0 and 7.5. The wavelengths used in kinetic measurements are indicated with a dotted line. Spectra are shown as the mean (*n* = 4 technical replicates).

#### 6. Additional Kinetic Data for Catalysis of UpA, ΨpA, and m$^1$ΨpA Cleavage

To assess differences in catalysis, we generated plots of the rate normalized to enzyme concentration for each substrate (Fig. S5). RNase A is a better catalyst for the cleavage of each dinucleotide substrates, and RNase A and RNase 1 differed most in catalysis of UpA cleavage.

The raw data, the code used to fit these data, and the raw data for Fig. 2 can be accessed at https://github.com/clair-gutierrez/pseudouridine-rnase.

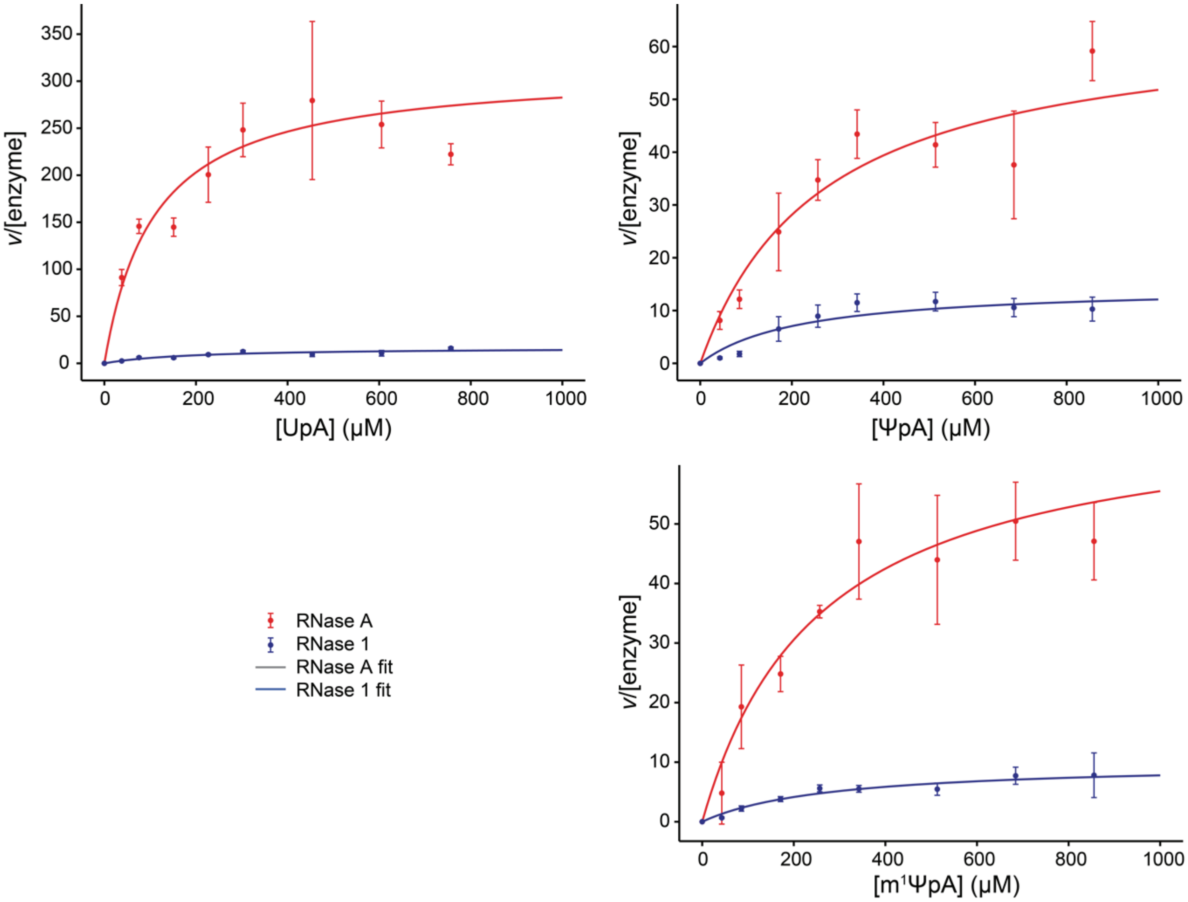


**FIGURE S5.** Enzyme concentration-adjusted rate plots for each dinucleotide substrate and ribonuclease at its optimal pH. Each data point is the mean ± SD ($n$ = 4 technical replicates).

#### 7. Kinetic Data for Catalysis of UpA Cleavage at Different pHs

To understand how pH might affect the cleavage of the different substrates, we conducted kinetic assays for RNase A at pH 7.5 and RNase 1 at pH 6.0, which differ from the optimal pH values for catalysis. These assays were done using the same buffers and conditions as described in the Materials and Methods section for these pH values, switching only the enzymes. These suboptimal pH conditions resulted in large $K_M$ values, making it difficult to obtain $k_{cat}$ and $K_M$ independently with accessible substrate concentrations (<1 mM). Instead, values of $k_{cat}/K_M$ were obtained in the $[S] \ll K_M$ regime, where the Michaelis–Menten equation collapses to a linear equation:

$$\frac{d[P]}{dt} \approx \frac{k_{cat}}{K_M}[E][S]_0 \tag{2}$$

In this regime, the accuracy in the values of $k_{cat}$ and $K_M$ is low, but the accuracy in their ratio is high (Bisswanger 2014). The data were fitted in two ways. First, the typical fit was done with the Michaelis–Menten equation, which provides $k_{cat}$ and $K_M$ independently, albeit with large errors, and those values were used to determine the ratio $k_{cat}/K_M$. Second, a linear fit was done by applying Eq 2 to kinetic data with $[S]_0 \leq 300$ μM. These latter values were used in Fig. 6.

**TABLE S3**. Values of $k_{cat}/K_M$ for the cleavage of UpA, ΨpA, and m$^1$ΨpA by RNase A and RNase 1 at the optimal pH of the other enzyme. Values from the fit to the Michaelis–Menten equation and from the linear fit in the region of $[S] \ll K_M$ and are reported as the mean ± SD for 4 technical replicates for each substrate condition.

| Ribonuclease | pH | Substrate | $k_{cat}/K_M$ ($10^5$ M$^{-1}$ s$^{-1}$) Michaelis–Menten fit | $k_{cat}/K_M$ ($10^5$ M$^{-1}$ s$^{-1}$) linear fit |
|---|---|---|---|---|
| RNase A | 7.5 | UpA | 1.36 ± 0.45 | 1.21 ± 0.07 |
| | 7.5 | ΨpA | 0.67 ± 0.78 | 0.69 ± 0.07 |
| | 7.5 | m$^1$ΨpA | 0.36 ± 0.25 | 0.25 ± 0.02 |
| RNase 1 | 6.0 | UpA | 0.68 ± 0.47 | 0.54 ± 0.02 |
| | 6.0 | ΨpA | 0.33 ± 0.24 | 0.33 ± 0.02 |
| | 6.0 | m$^1$ΨpA | 0.36 ± 0.91 | 0.35 ± 0.01 |

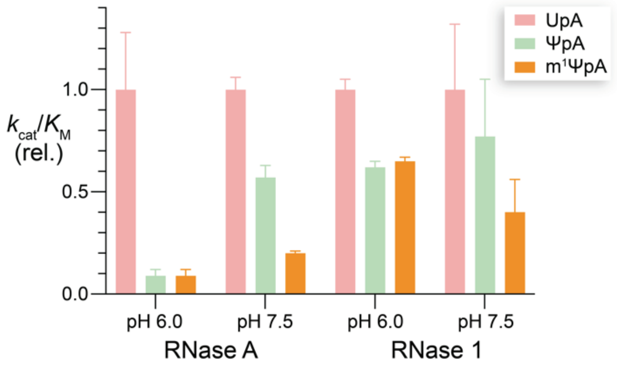


**FIGURE S6.** pH-Dependence of the relative $k_{cat}/K_M$ values for catalysis of the cleavage of dinucleotide substrates by RNase A and RNase 1 at 25 °C. Values are the mean ± SD of 4 replicates.

#### 8. Statistics for X-ray Crystallography

**TABLE S4**. Data collection, processing, and refinement statistics of the complexes of RNase A with U>v, Ψ>v, and $m^1$Ψ>v*

| | **RNase A·U>v** | **RNase A·Ψ>v** | **RNase A·m$^1$Ψ>v** |
|---|---|---|---|
| PDB code | 9ncs | 9o4v | 9o5b |
| Wavelength (Å) | 1.54 | 1.54 | 1.54 |
| Resolution range (Å) | 31.06–1.83 (1.895–1.83) | 29.17–1.70 (1.761–1.70) | 25.08–1.71 (1.771–1.71) |
| Space group | C 1 2 1 | C 1 2 1 | C 1 2 1 |
| Unit cell: *a*, *b*, *c* (Å)<br>*α*, *β*, *γ* (°) | 100.053, 32.677, 72.327<br>90, 90.361, 90 | 99.631, 32.512, 71.879<br>90, 90.093, 90 | 100.31, 32.83, 72.67<br>90, 90.547, 90 |
| Total reflections | 70224 (4198) | 74892 (1909) | 98194 (2106) |
| Unique reflections | 19255 (1316) | 22942 (1022) | 22880 (1172) |
| Multiplicity | 3.6 (3.2) | 3.3 (1.9) | 4.3 (1.8) |
| Completeness (%) | 91.40 (62.95) | 89.02 (40.33) | 87.84 (45.16) |
| Mean $I/\sigma$(I) | 13.42 (2.97) | 17.54 (3.82) | 14.33 (2.47) |
| Wilson *B*-factor (Å$^2$) | 20.63 | 18.55 | 19.62 |
| $R_{merge}$ | 0.06153 (0.375) | 0.04228 (0.1611) | 0.06403 (0.1723) |
| $R_{meas}$ | 0.07231 (0.444) | 0.05009 (0.2218) | 0.07191 (0.2156) |
| $R_{pim}$ | 0.03766 (0.2352) | 0.02652 (0.1513) | 0.03201 (0.1267) |
| $CC_{½}$ | 0.995 (0.934) | 0.998 (0.956) | 0.995 (0.956) |
| CC* | 0.999 (0.983) | 0.999 (0.989) | 0.999 (0.989) |
| Reflections used in refinement | 19251 (1315) | 22941 (1022) | 22873 (1170) |
| Reflections used for $R_{free}$ | 1929 (130) | 1989 (92) | 2008 (107) |
| $R_{work}$ | 0.1793 (0.4433) | 0.1712 (0.4870) | 0.1789 (0.3680) |
| $R_{free}$ | 0.2139 (0.5077) | 0.2023 (0.4634) | 0.2169 (0.3633) |
| $CC_{work}$ | 0.950 (0.843) | 0.950 (0.851) | 0.949 (0.719) |
| $CC_{free}$ | 0.954 (0.736) | 0.937 (0.750) | 0.922 (0.673) |
| No. non-hydrogen atoms<br>macromolecules<br>ligands<br>solvent | 2278<br>1889<br>160<br>229 | 2321<br>1872<br>185<br>272 | 2273<br>1854<br>184<br>247 |
| Protein residues | 243 | 242 | 239 |
| RMSD bond lengths (Å) | 0.003 | 0.004 | 0.005 |
| RMSD bond angles (°) | 0.53 | 0.60 | 0.64 |
| Ramachandran favored | 96.60% | 97.02 | 96.09% |
| Ramachandran allowed | 3.40% | 2.98% | 3.91% |
| Ramachandran outliers | 0.00% | 0.00% | 0.00% |
| Rotamer outliers (%) | 0.93 | 0.93 | 2.80 |
| Clashscore | 7.29 | 9.63 | 8.44 |
| Average *B*-factor (Å$^2$)<br>macromolecules<br>ligands<br>solvent | 28.53<br>25.24<br>64.09<br>30.87 | 24.39<br>21.46<br>50.04<br>27.85 | 29.49<br>26.22<br>60.93<br>32.16 |
| No. TLS groups | 13 | 5 | 8 |

*Statistics for the highest-resolution shell are shown in parentheses.

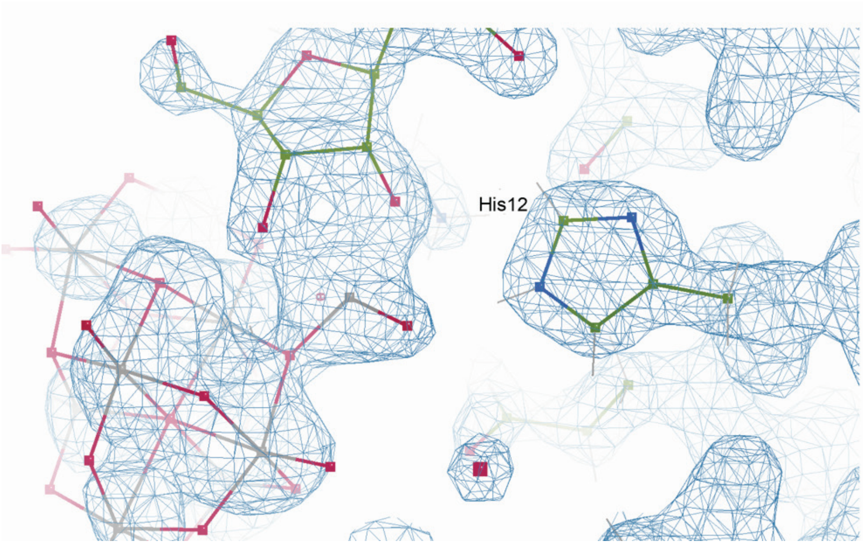


**FIGURE S7.** Electron density near the vanadyl group in the crystal structure of the RNase A·m[1]Ψ>v complex. Vanadium atoms are depicted in grey, oxygen atoms in pink, carbon atoms in green, and nitrogen atoms in blue. The vanadyl group adopts a tetrahedral geometry that includes the 2′ and 3′ oxygens of ribose, an oxygen in a proximal decavanadate, and a nonbridging oxygen. The image is depicted at a contour level of 0.75.

#### 9. Decavanadates in Crystal Structures

Decavanadate was found in each crystal structure (Fig. S8). That is not surprising per se because the crystals were grown at pH 5.5, where decavanadates are a predominant species (Aureliano et al. 2022), and because decavanadate is known to bind to RNase A (Messmore and Raines 2000). A previous X-ray crystallographic study of RNase A·U>v did not report the presence of decavanadate (Ladner et al. 1997), though the mother liquor therein contained a high concentration of 2-methyl-2-propanol, which can precipitate vanadate.

In addition, the electron density of the vanadyl group in each complex was in a tetrahedral conformation (Fig. S7) and appeared to be a complex with a proximal decavanadate. These interactions had no impact on nucleobase binding in the active site (Fig. 3). To the extent of our knowledge this vanadate species (decavanadate in complex with an additional oxidovanadate) has not been reported previously as a stable species at this pH and could be stabilized by the enzymic active site.

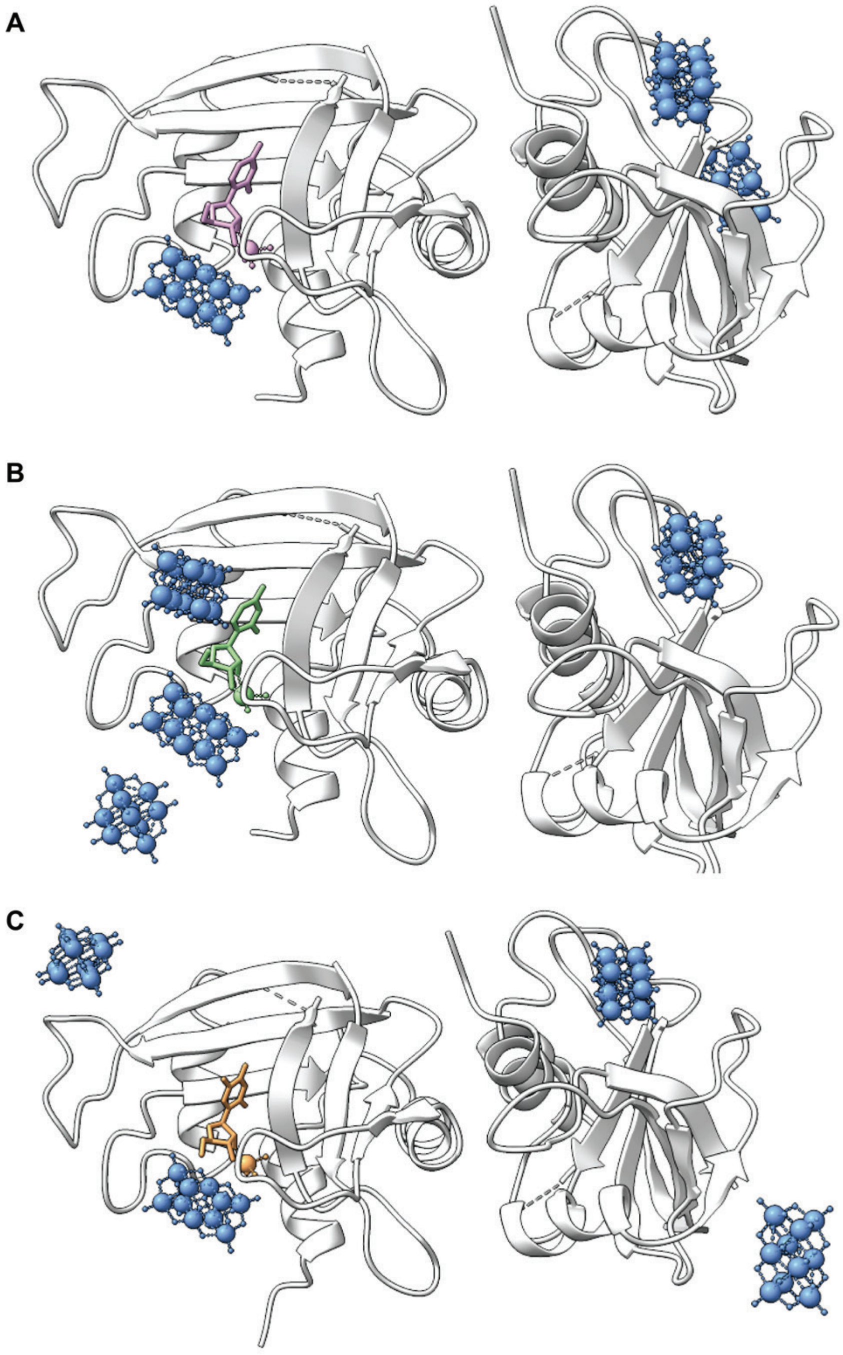


**FIGURE S8.** Vanadium in the unit cells of (*A*) U>v, (*B*) Ψ>v, and (*C*) m¹Ψ>v. Nucleoside 2′,3′-cyclic vanadate complexes are in only one monomer of the unit cell whereas decavanadates exist in different regions around the two monomers.

### 10. Computational Analyses

#### 10.1. Top-Ranked Docking Poses of Enzyme·Substrate Complexes

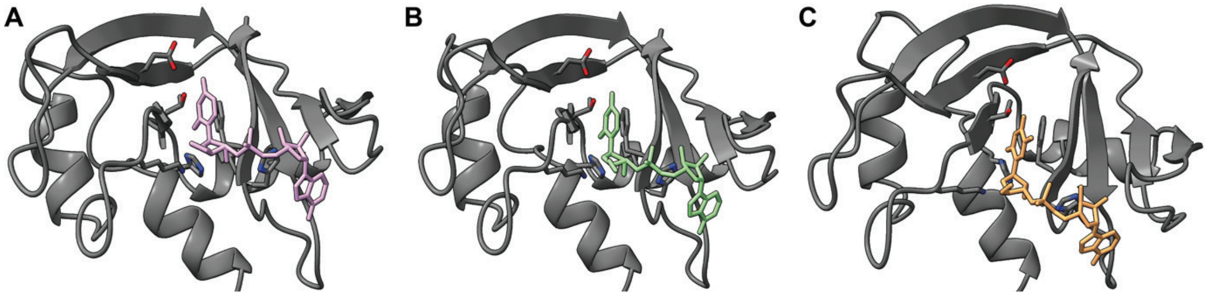


**FIGURE S9.** Top-ranked docking poses of UpA (*A*, pink), ΨpA (*B*, green) and m[1]ΨpA (*C*, orange) bound to RNase A (gray).

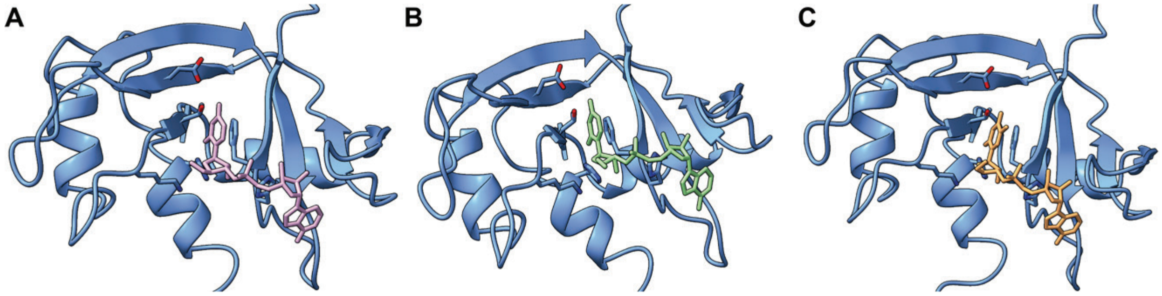


**FIGURE S10.** Top-ranked docking poses of UpA (*A*, pink), ΨpA (*B*, green) and m[1]ΨpA (*C*, orange) bound to RNase 1 (blue).

##### 10.2. Molecular Dynamics Simulations of RNase A·Substrate Complexes

MD simulations for each substrate were initiated from the RNase A·substrate complexes resulting from molecular docking studies (Fig. S9) and were performed for 1500 ns. We used the TIP3P water model along with the ff14SB and gaff parameter sets for the protein and ligands, respectively. For each simulation, an average structure was generated using 7,500 frames from the final 750 ns of the simulation.

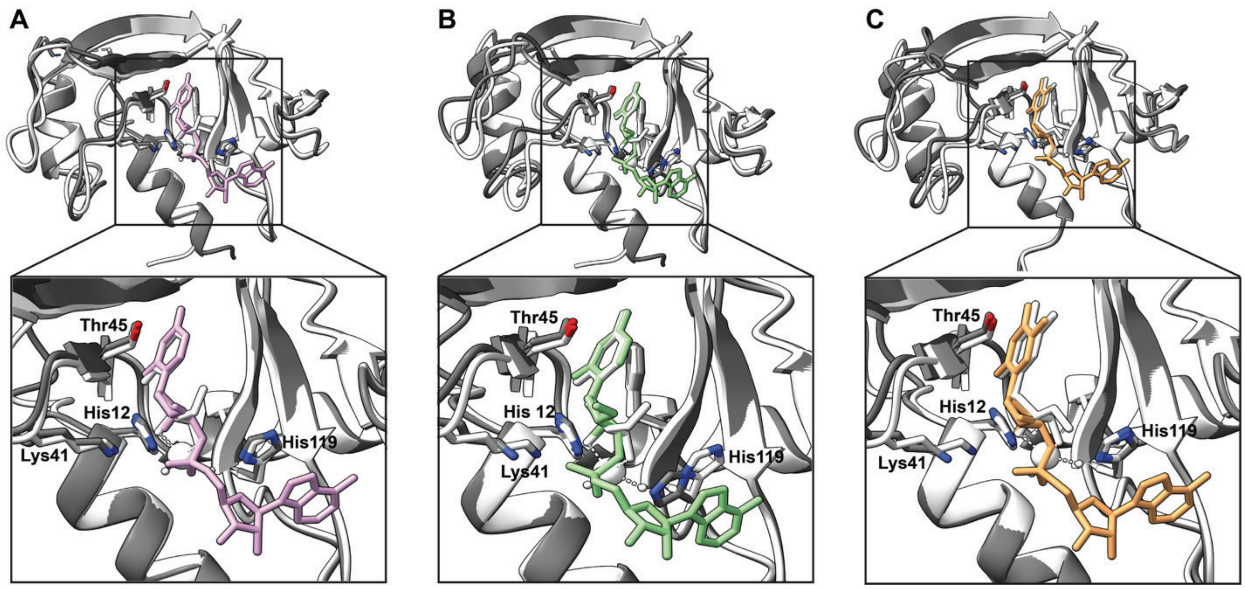


**FIGURE S11.** Overlay of UpA (*A*, pink), ΨpA (*B*, green) and $m^1$ΨpA (*C*, orange) bound to RNase A (gray) obtained from MD simulations with the crystal structures of U>v (*A*, white), Ψ>v (*B*, white), and $m^1$Ψ>v (*C*, white) bound to RNase A (white).

##### 10.3. Molecular Dynamics Simulations of RNase 1·Substrate Complexes

MD simulations for each substrate were initiated from the RNase 1·substrate complexes resulting from molecular docking studies (Fig. S10) and were performed for 750 ns. We used the TIP3P water model and the ff14SB parameter set for the protein. For the ligands, the RNA force field OL3 and modrna08 parameters were applied to natural and modified dinucleotides, respectively. For each simulation, an average structure was generated using 7500 frames.

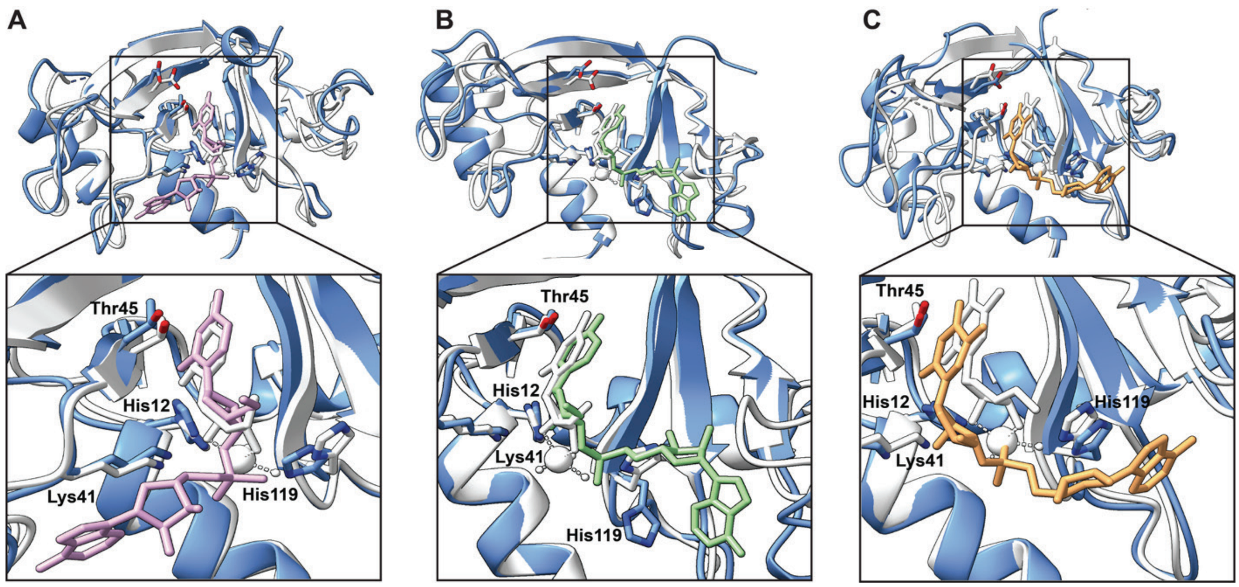


**FIGURE S12.** Overlay of UpA (*A*, pink), ΨpA (*B*, green) and m[1]ΨpA (*C*, orange) bound to RNase 1 (blue) obtained from MD simulations with the crystal structures of U>v (*A*, white), Ψ>v (*B*, white), and m[1]Ψ>v (*C*, white) bound to RNase A (white).

##### 10.4. RMSD Plots for Molecular Dynamics Simulations with RNase A and RNase 1

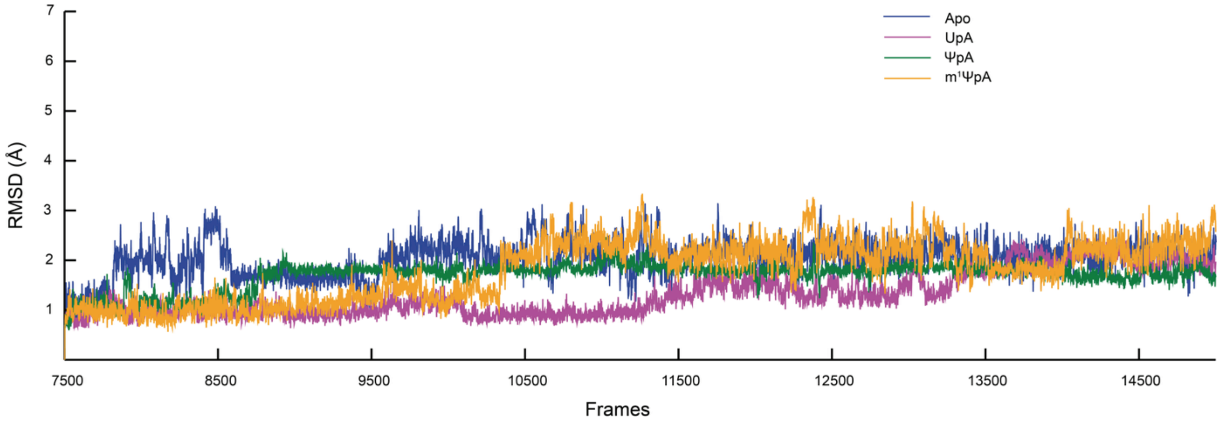


**FIGURE S13.** RMSD plots based on the last 750 ns of each of the four simulations of RNase A. For the simulations, the TIP3P water model was used along with the ff14SB and gaff parameter sets for the protein and ligands, respectively. The plateaued RMSDs indicate the stabilization of the protein backbone.

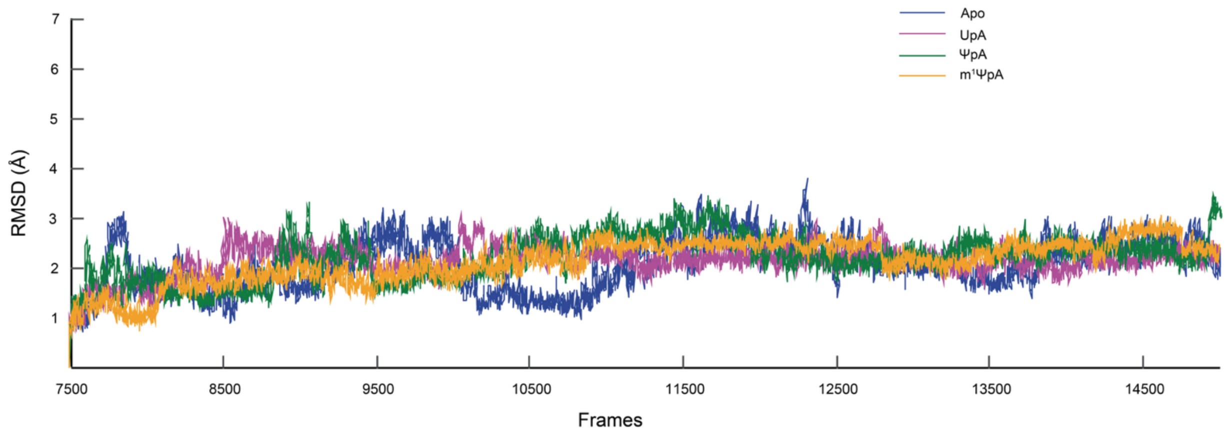


**FIGURE S14.** RMSD plots based on the last 750 ns of each of the four simulations of RNase 1. For the simulations, the TIP3P water model was used along with the ff14SB and gaff parameter sets for the protein and ligands, respectively. The plateaued RMSDs indicate the stabilization of the protein backbone.

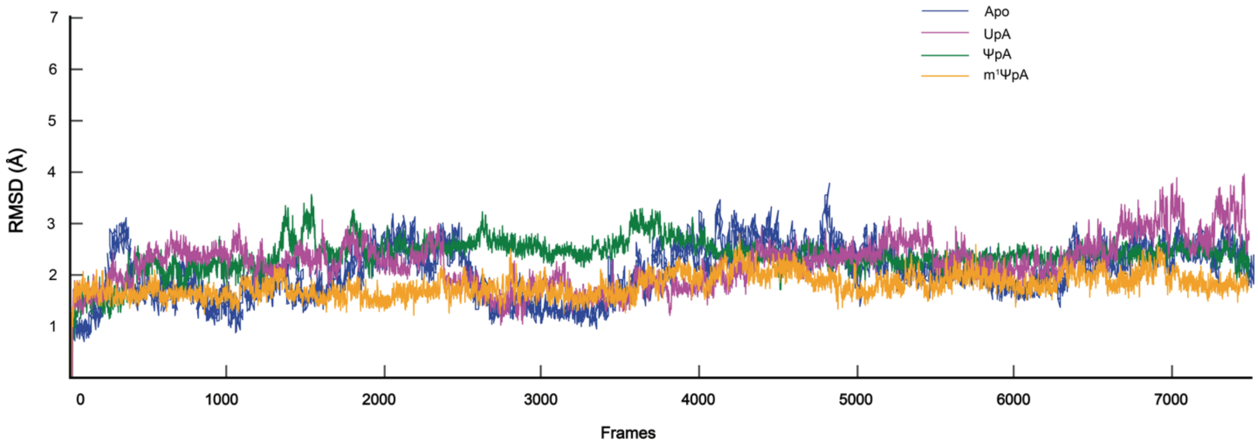


**FIGURE S15.** RMSD plots of each of the four 750 ns simulations of RNase 1. For the simulations, the TIP3P water model was used along with the ff14SB parameter set for the protein. For the ligands, the RNA force field OL3 and modrna08 parameters were applied to the canonical and modified dinucleotides, respectively. The plateaued RMSDs indicate the stabilization of the protein backbone.

#### 11. Kinetic Data for Non-Enzymatic UpA and m¹ΨpA Cleavage

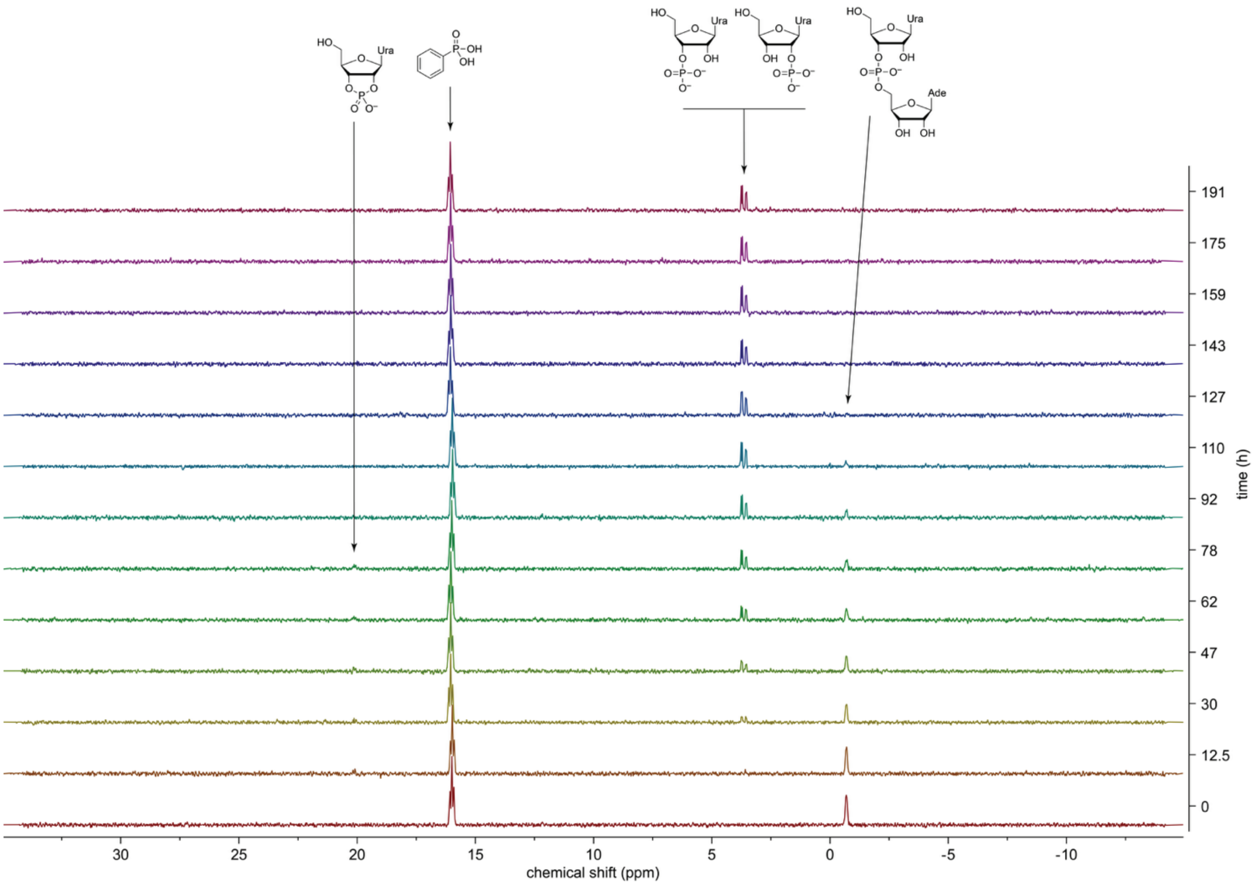


**FIGURE S16.** Time course of the $^{31}$P NMR spectra for the uncatalyzed cleavage of UpA.

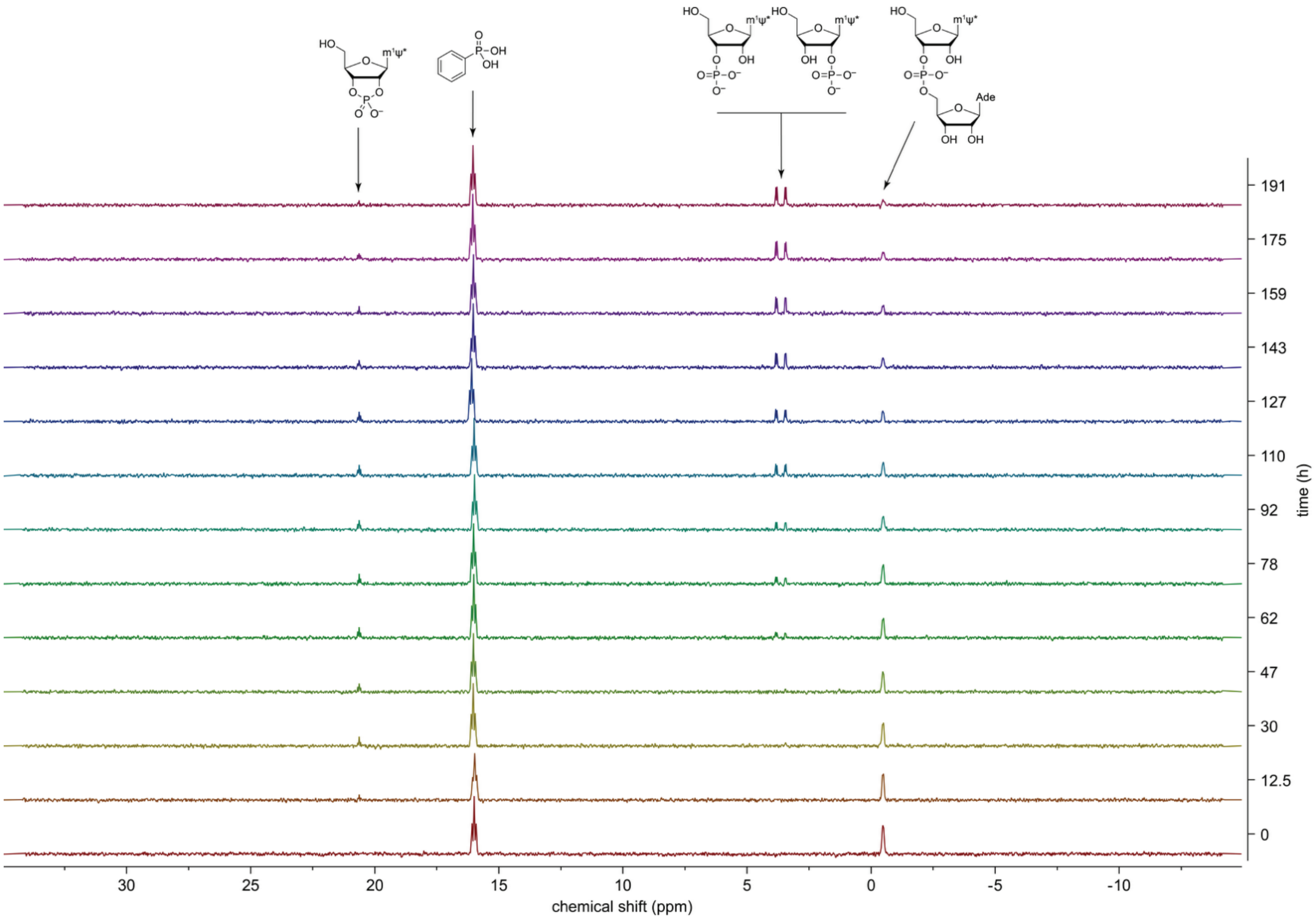


**FIGURE S17.** Time course of the $^{31}$P NMR spectra for the uncatalyzed cleavage of m$^{1}\Psi$pA. m$^{1}\Psi$* = *N*$^{1}$-methylpseudouracil nucleobase.

#### 12. NMR Spectra

$^{1}$H NMR spectrum (500 MHz, $CDCl_3$) of **1b**

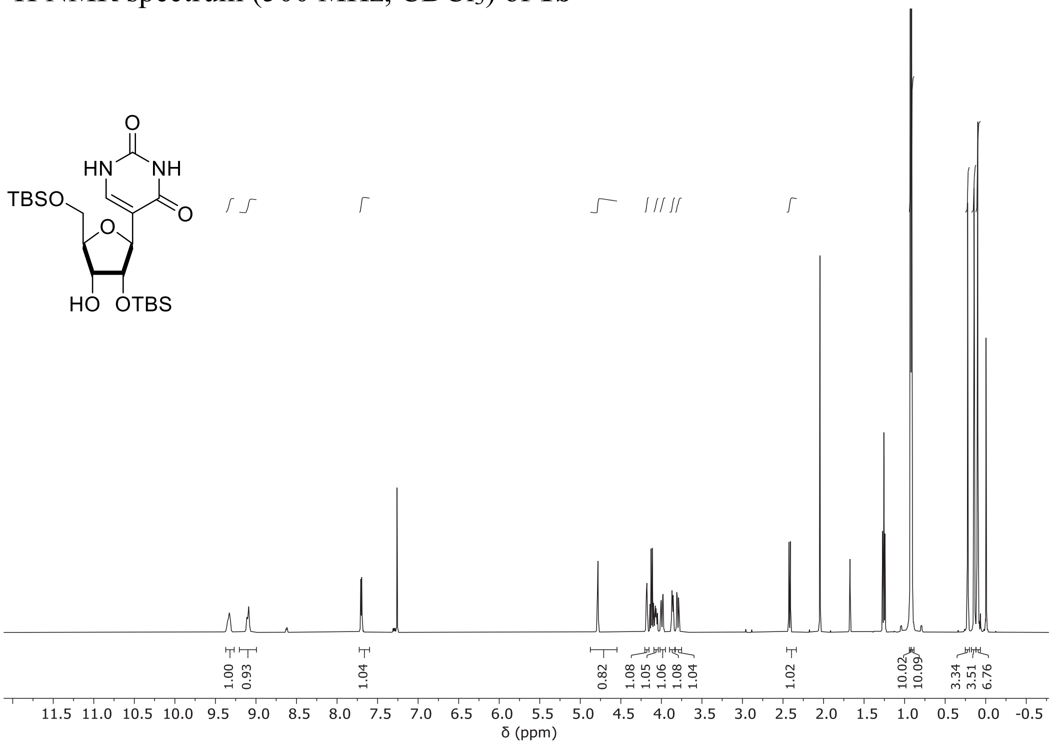


$^{13}$C NMR spectrum (126 MHz, $CDCl_3$) of **1b**

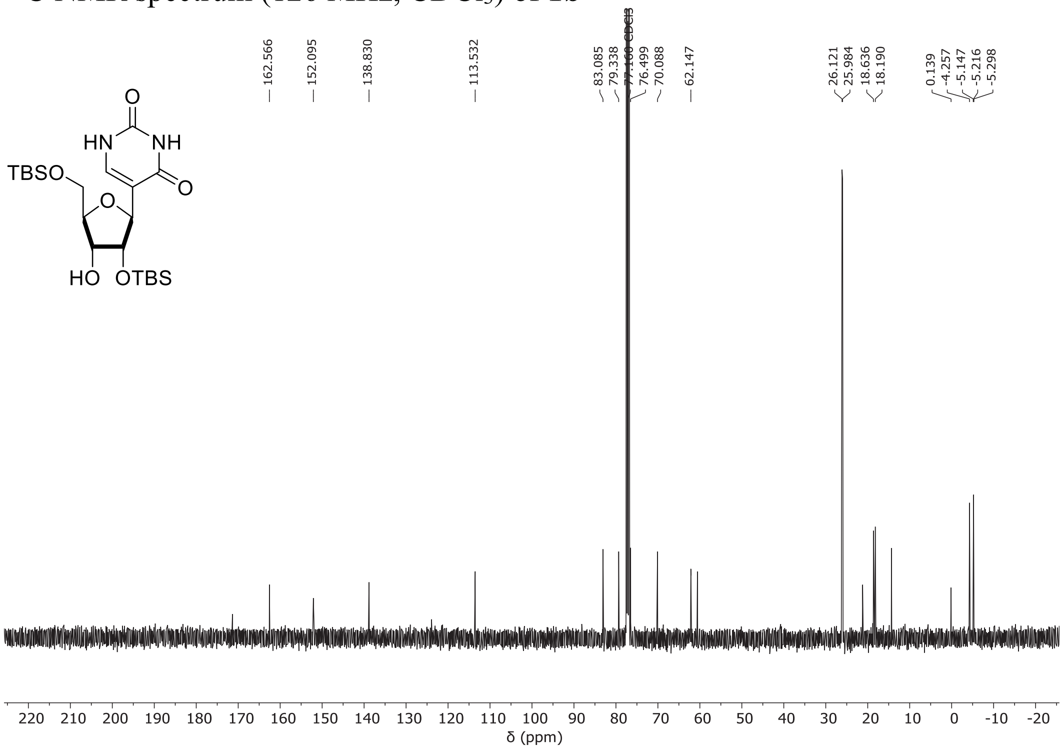

$^{1}$H NMR spectrum (500 MHz, $CDCl_3$) of **1c**

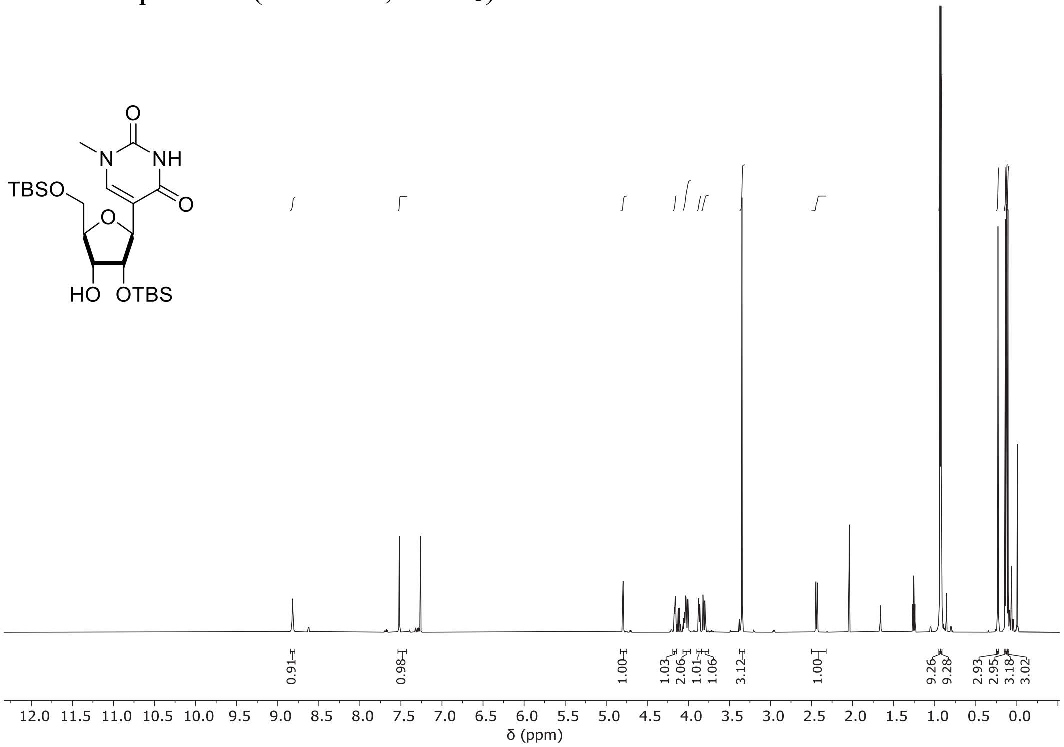


$^{13}$C NMR spectrum (126 MHz, $CDCl_3$) of **1c**

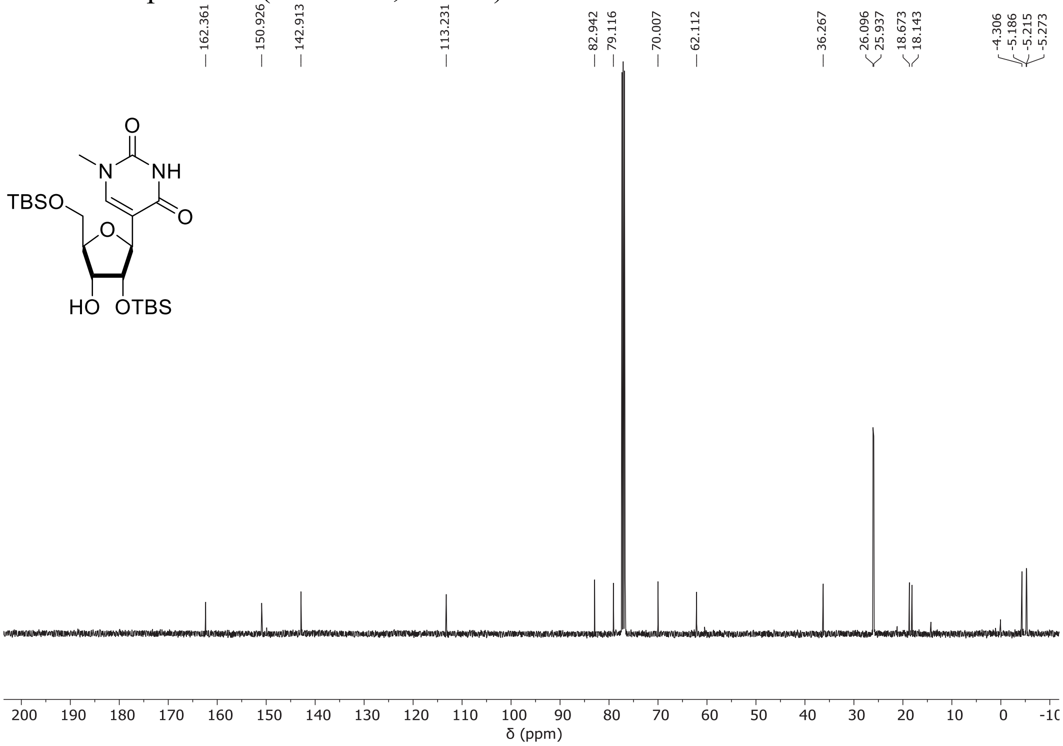

$^{31}$P NMR spectrum (162 MHz, $CDCl_3$) of **2a**

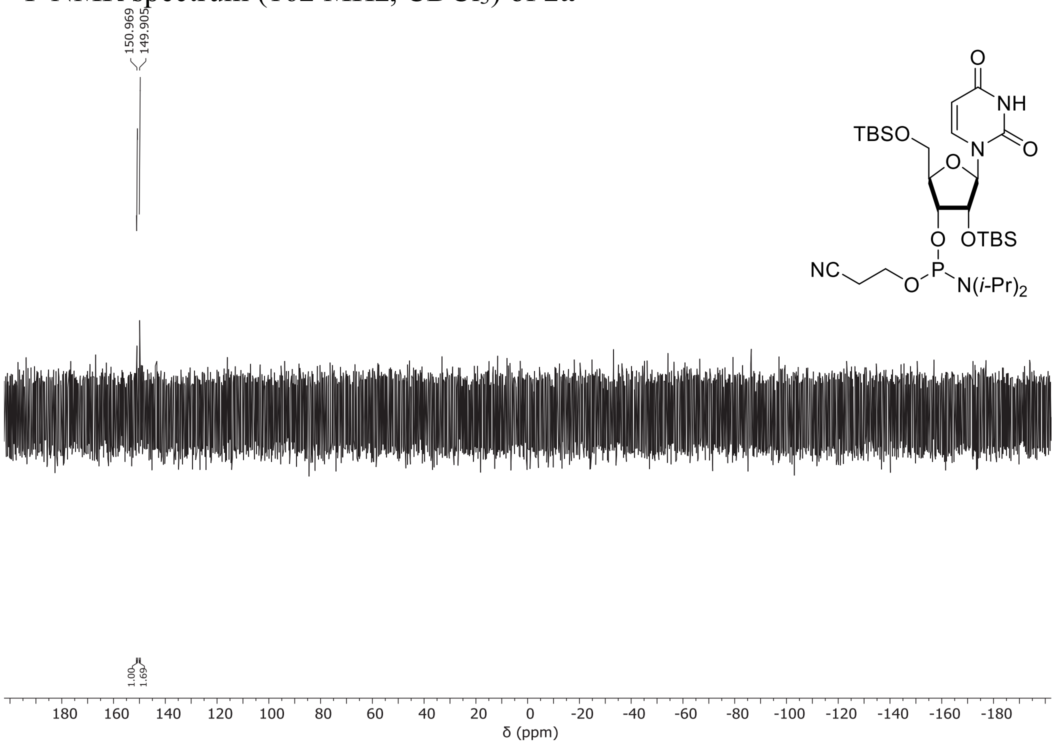


$^{31}$P NMR spectrum (162 MHz, $CDCl_3$) of **2b**

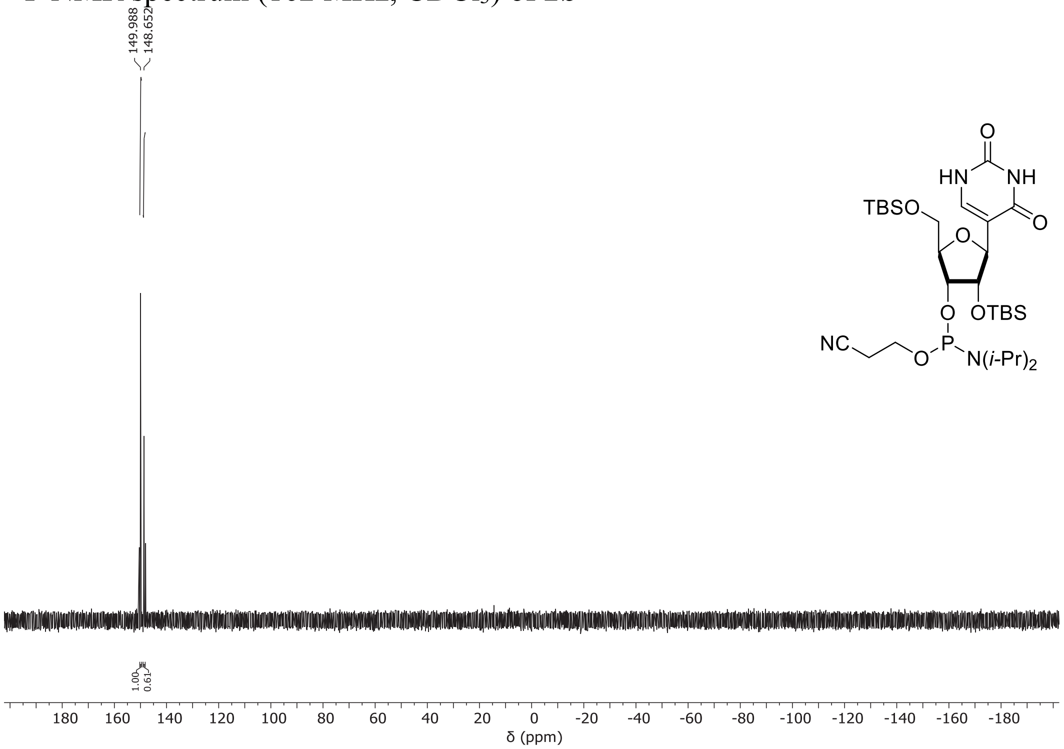

[31]P NMR spectrum (162 MHz, $CDCl_3$) of **2c**

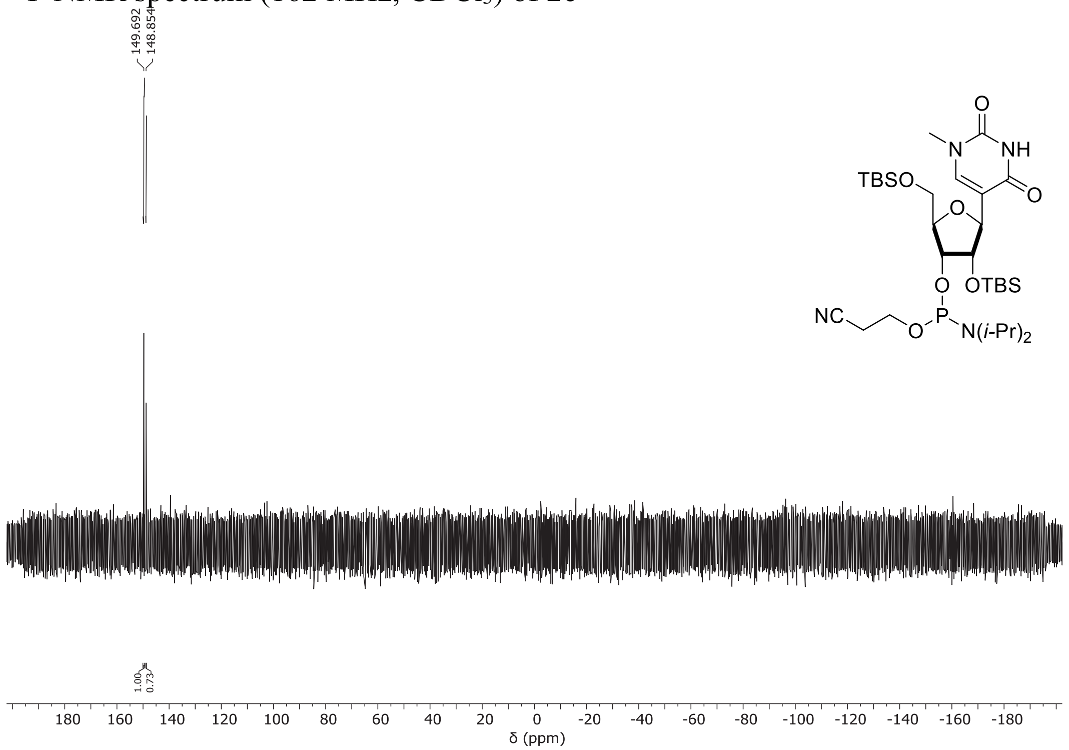


[1]H NMR spectrum (500 MHz, $CDCl_3$) of **5a**

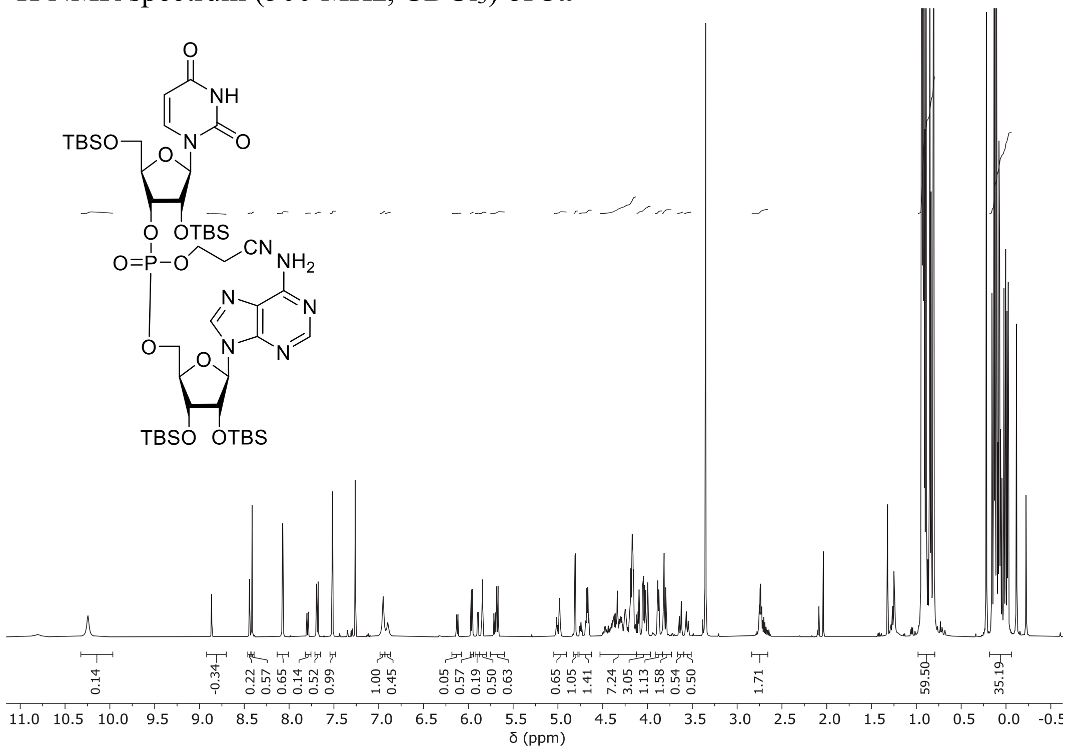

[13]C NMR spectrum (126 MHz, $CDCl_3$) of **5a**

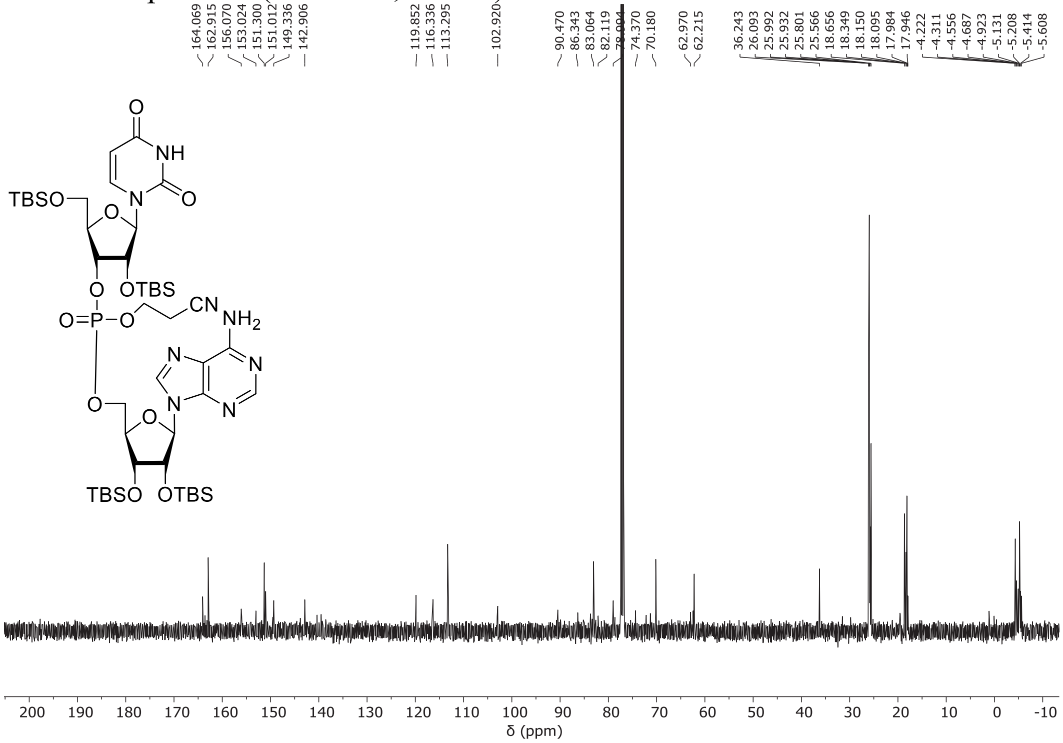


[31]P NMR spectrum (162 MHz, $CDCl_3$) of **5a**

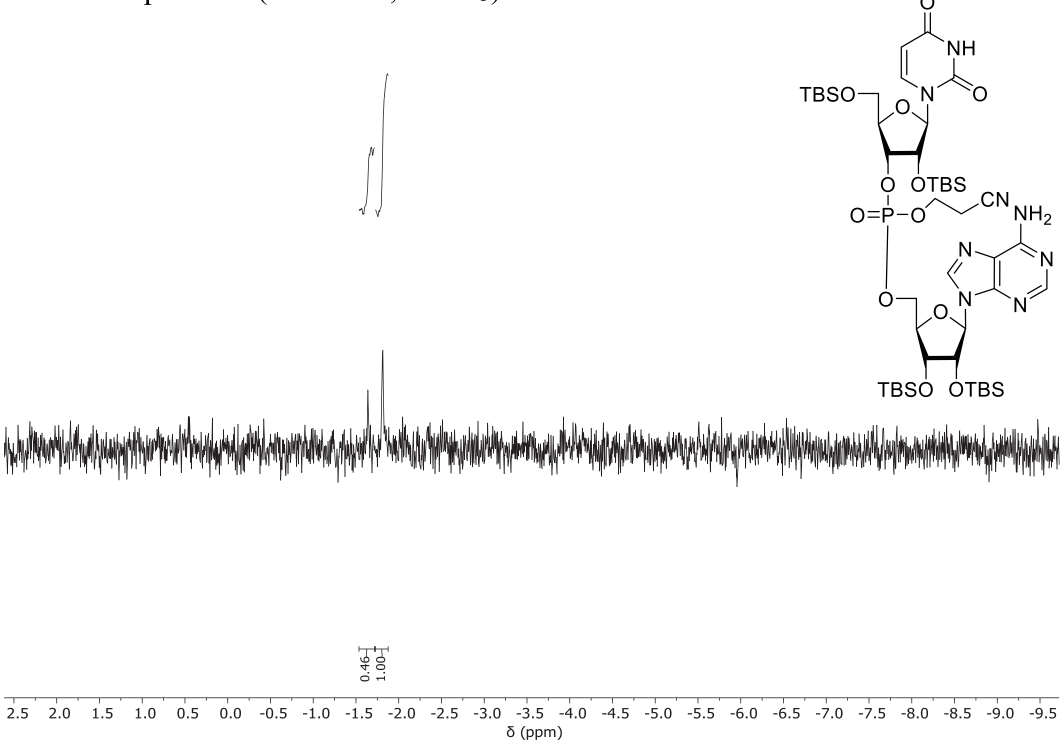

$^{1}$H NMR spectrum (500 MHz, $CDCl_3$) of **5b**

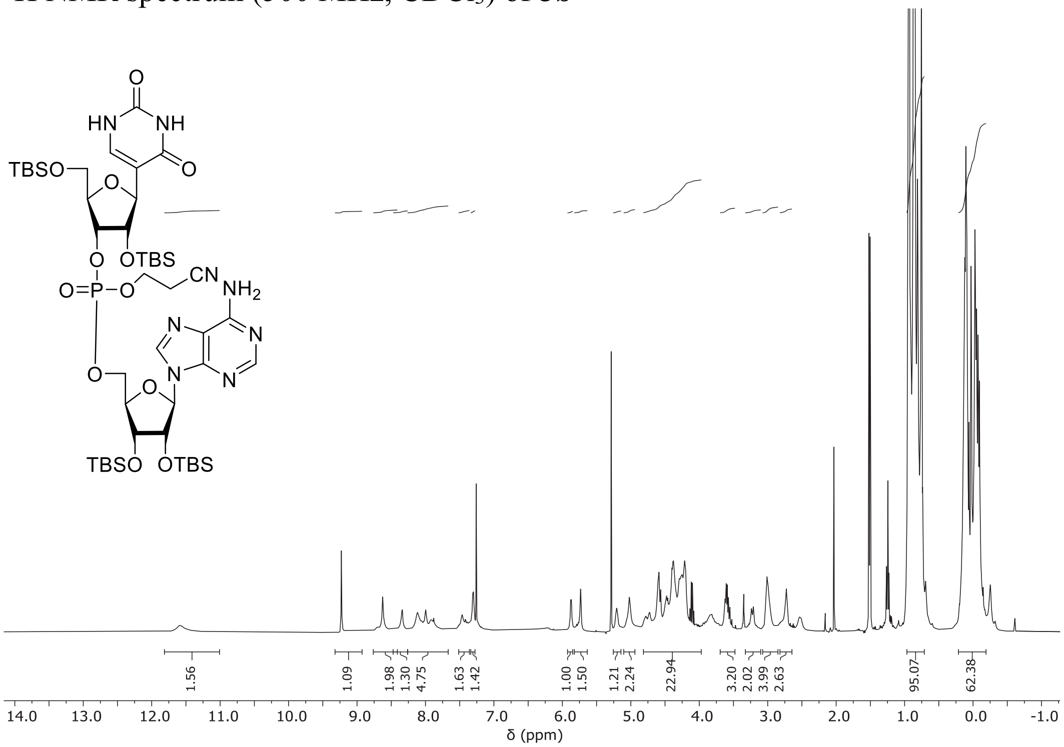


$^{13}$C NMR spectrum (126 MHz, $CDCl_3$) of **5b**

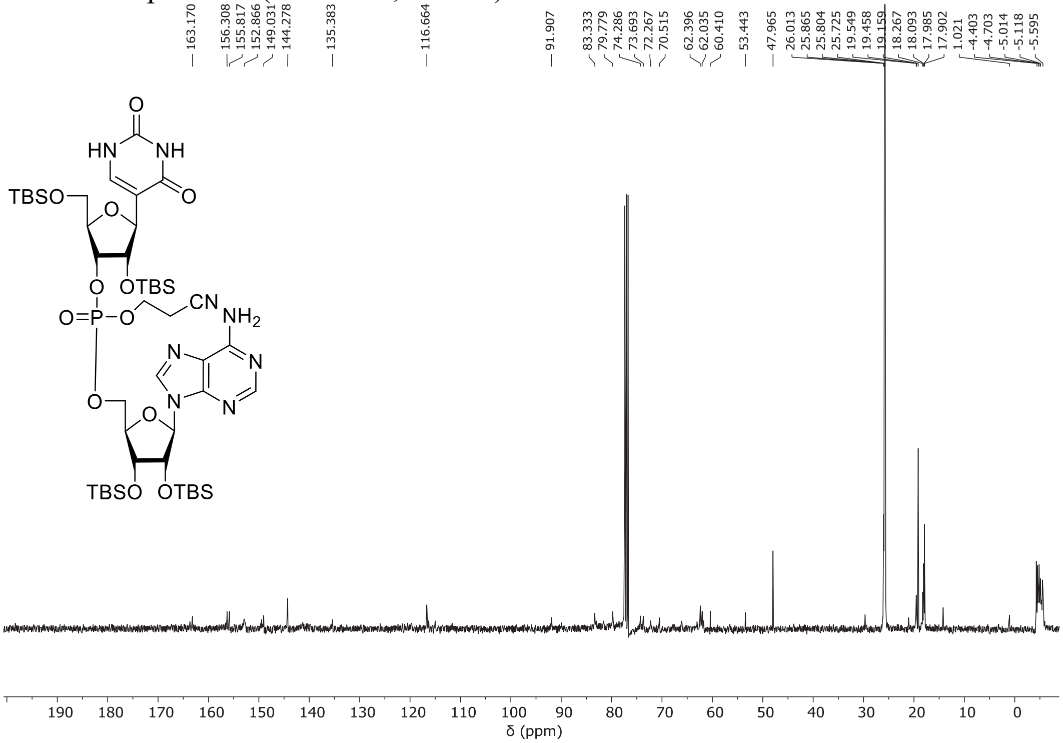

$^{31}$P NMR spectrum (162 MHz, $CDCl_3$) of **5b**

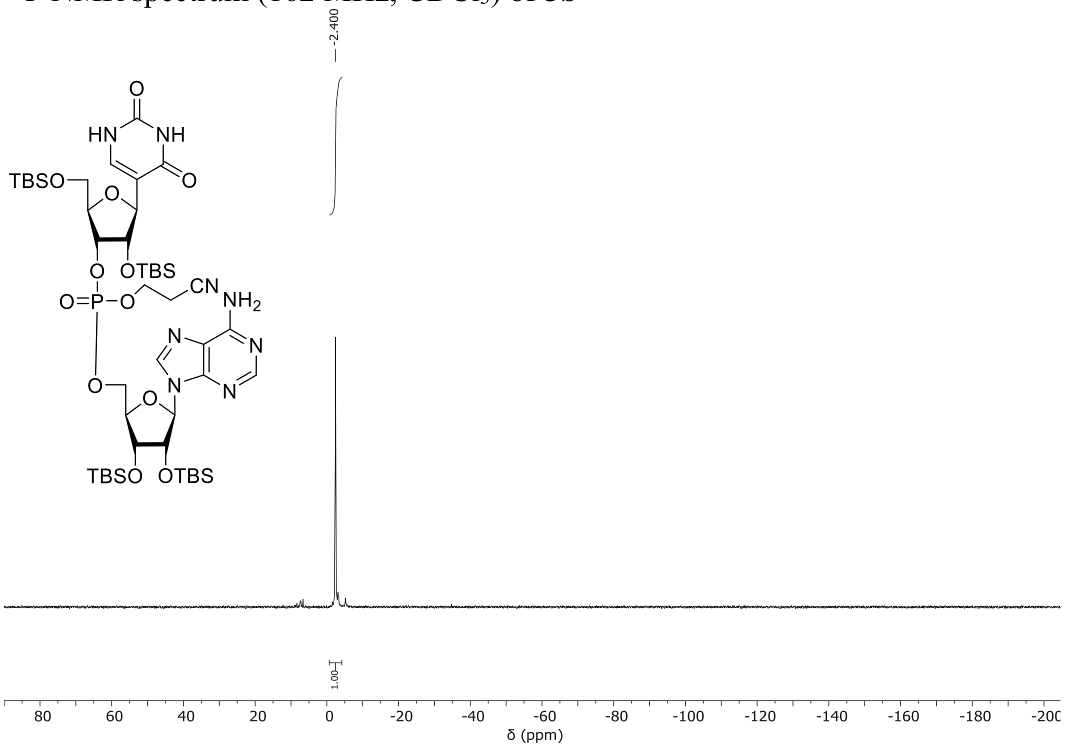


$^{1}$H NMR spectrum (500 MHz, $CDCl_3$) of **5c**

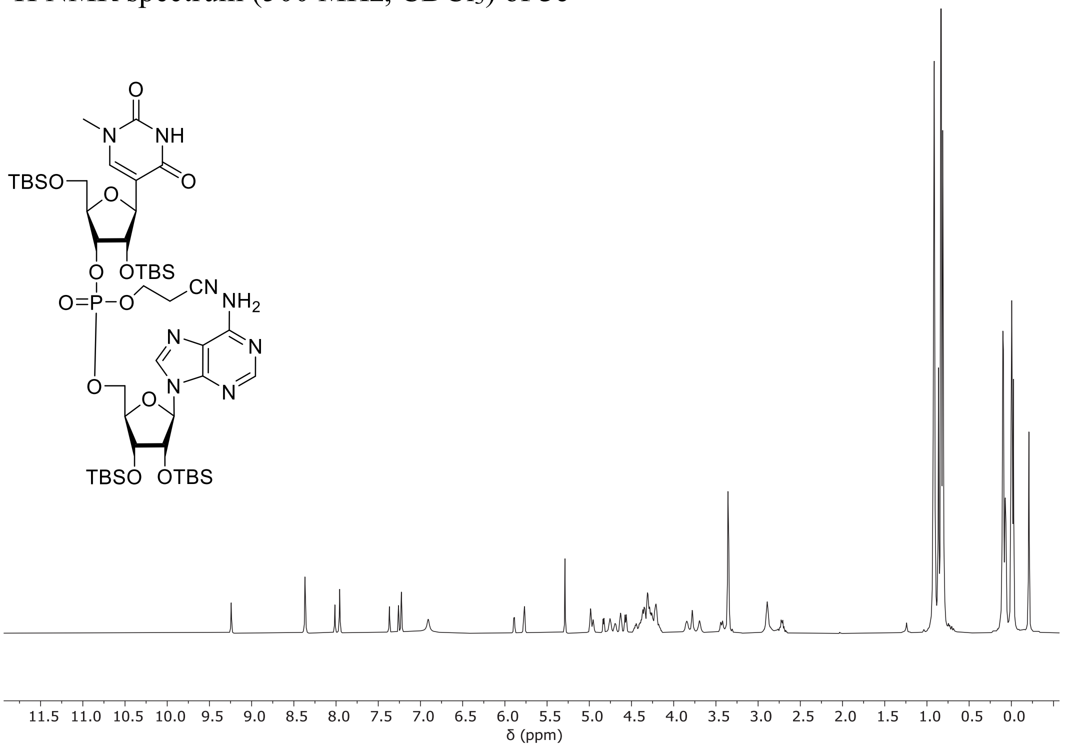

[13]C NMR spectrum (126 MHz, $CDCl_3$) of **5c**

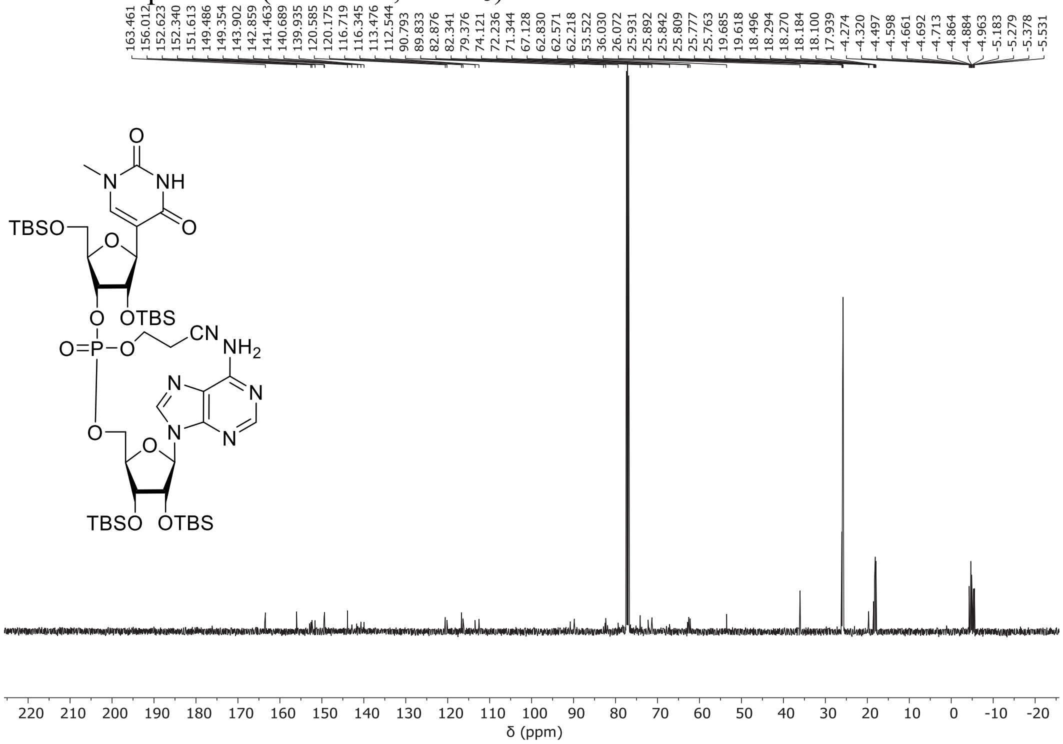

[31]P NMR spectrum (162 MHz, $CDCl_3$) of **5c**

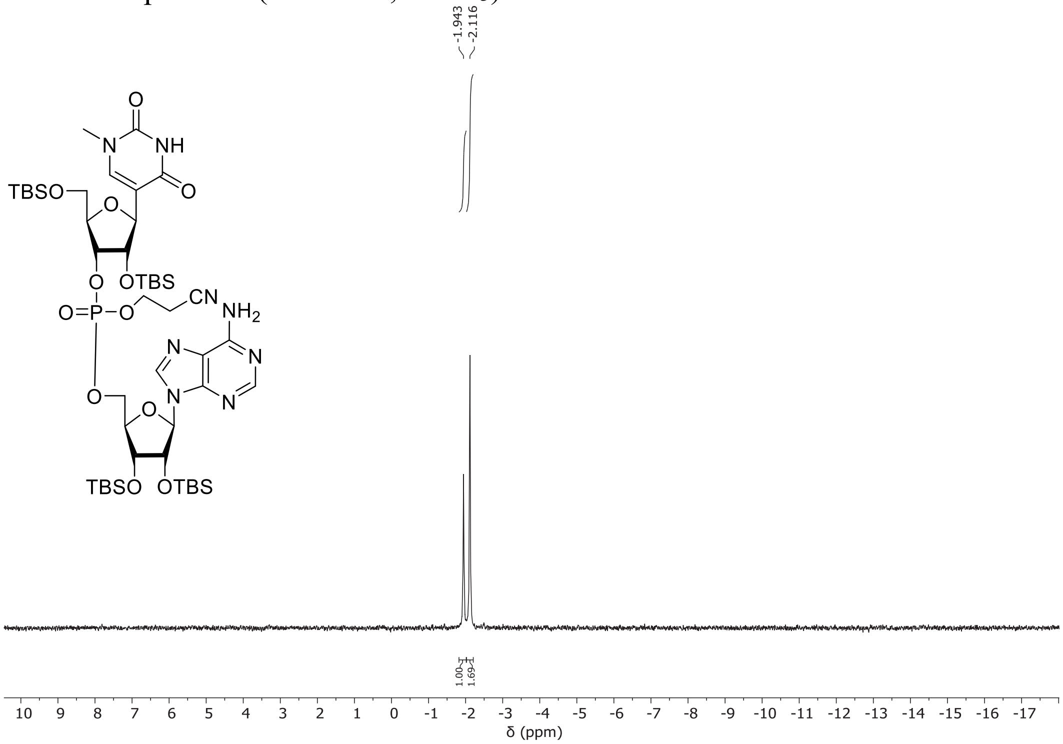

[1]H NMR spectrum (400 MHz, $D_2O$) of **UpA**·$NEt_3$

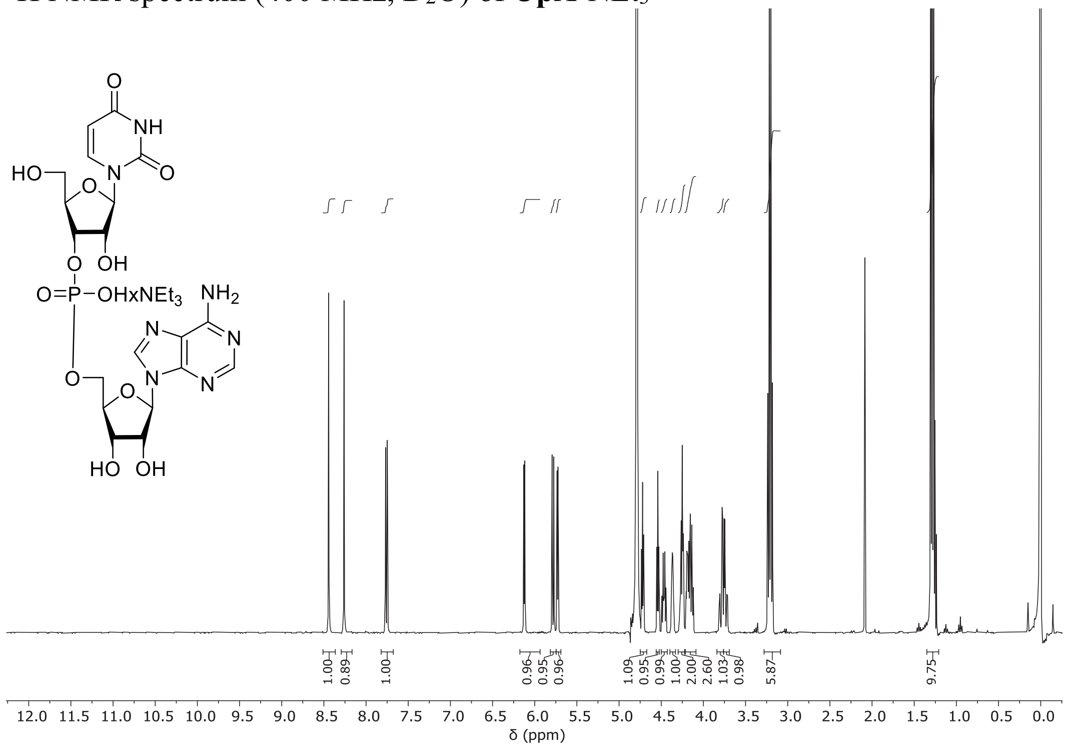

[13]C NMR spectrum (101 MHz, $D_2O$) of **UpA**·$NEt_3$

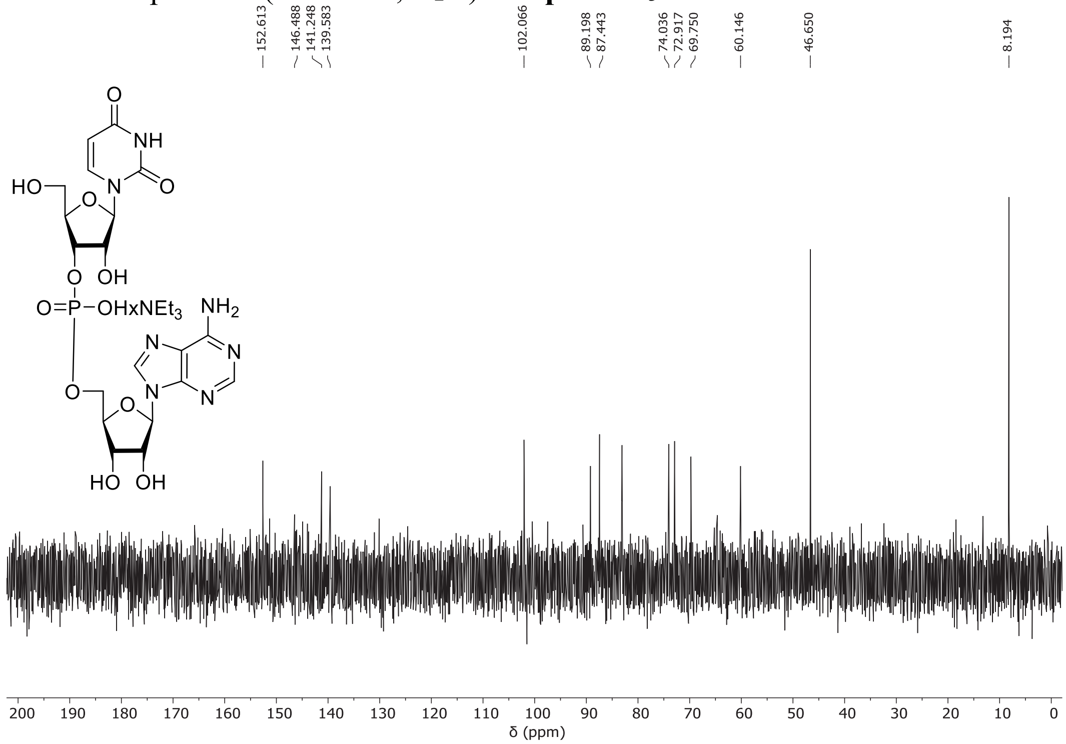

$^{31}$P NMR spectrum (162 MHz, $D_2O$) of **UpA**·$NEt_3$

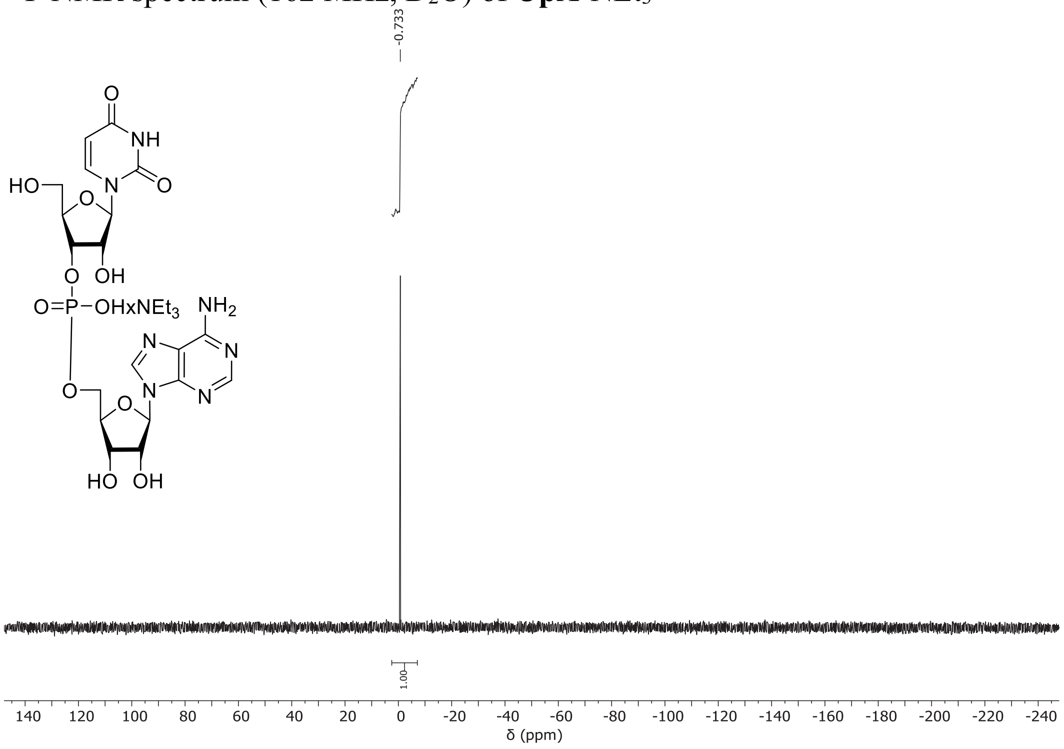

$^{1}$H NMR spectrum (500 MHz, $D_2O$) of **ΨpA**·$NEt_3$

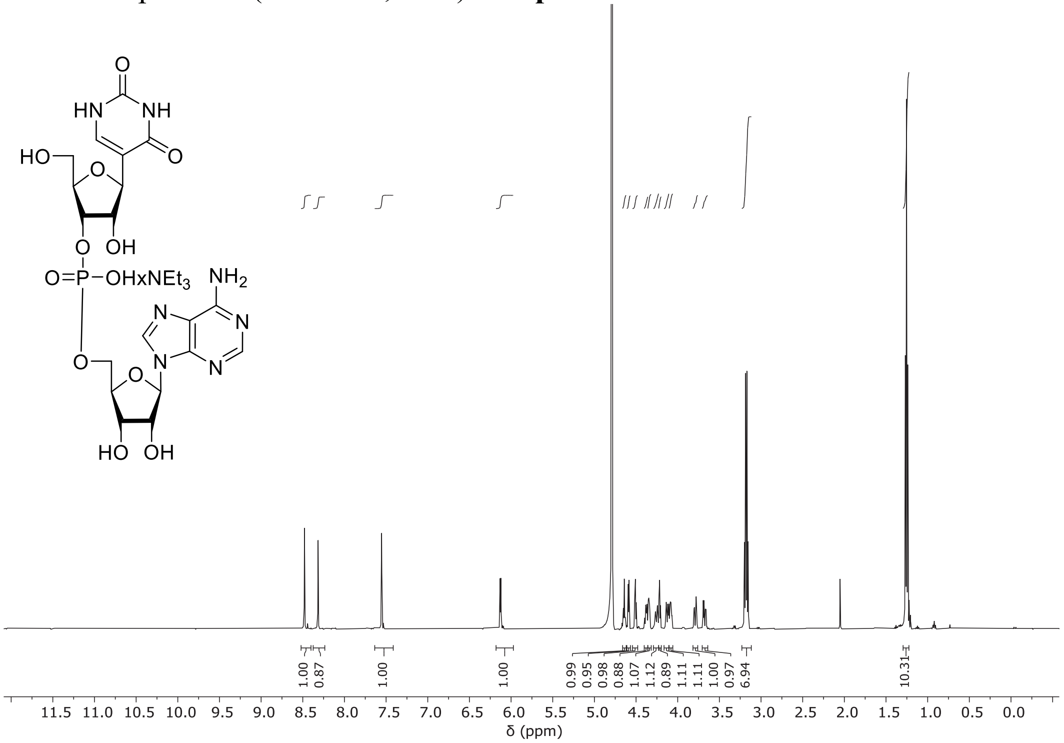

$^{13}$C NMR spectrum (126 MHz, $D_2O$) of **ΨpA**·$NEt_3$

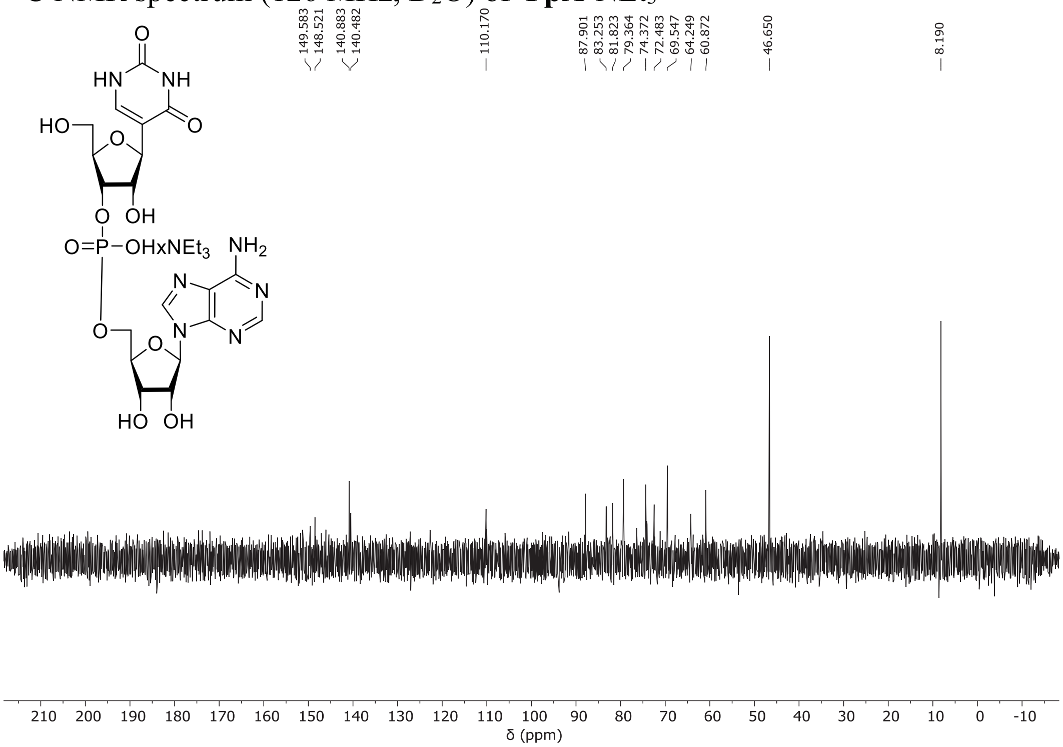


$^{31}$P NMR spectrum (202 MHz, $D_2O$) of **ΨpA**·$NEt_3$

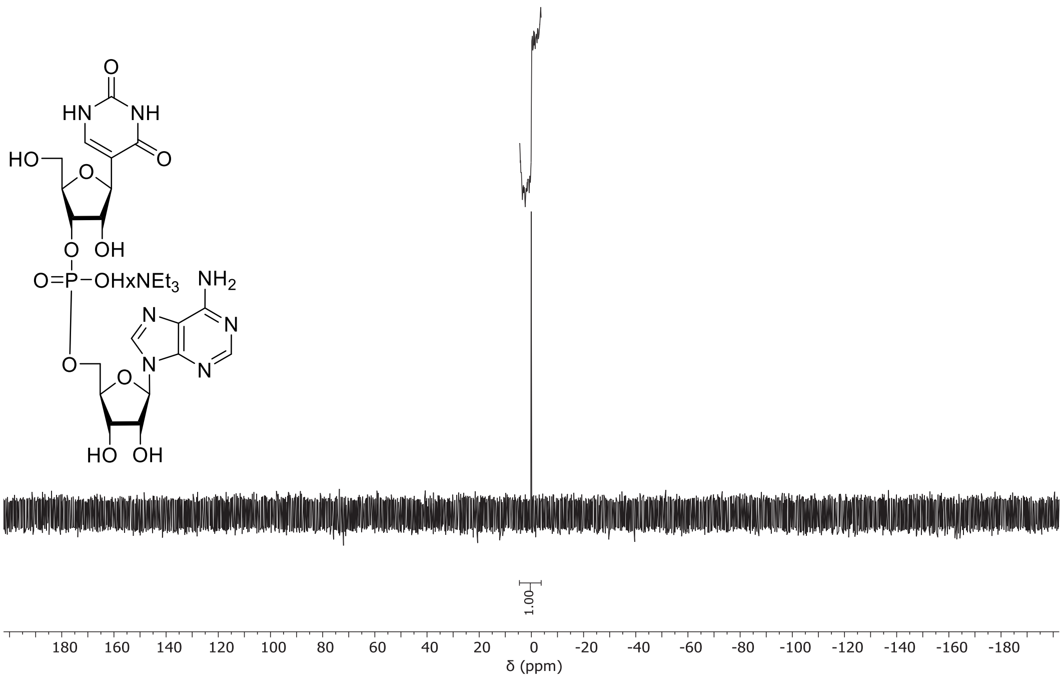

$^{1}$H NMR spectrum (500 MHz, $D_2O$) of **m$^1$ΨpA**·$NEt_3$

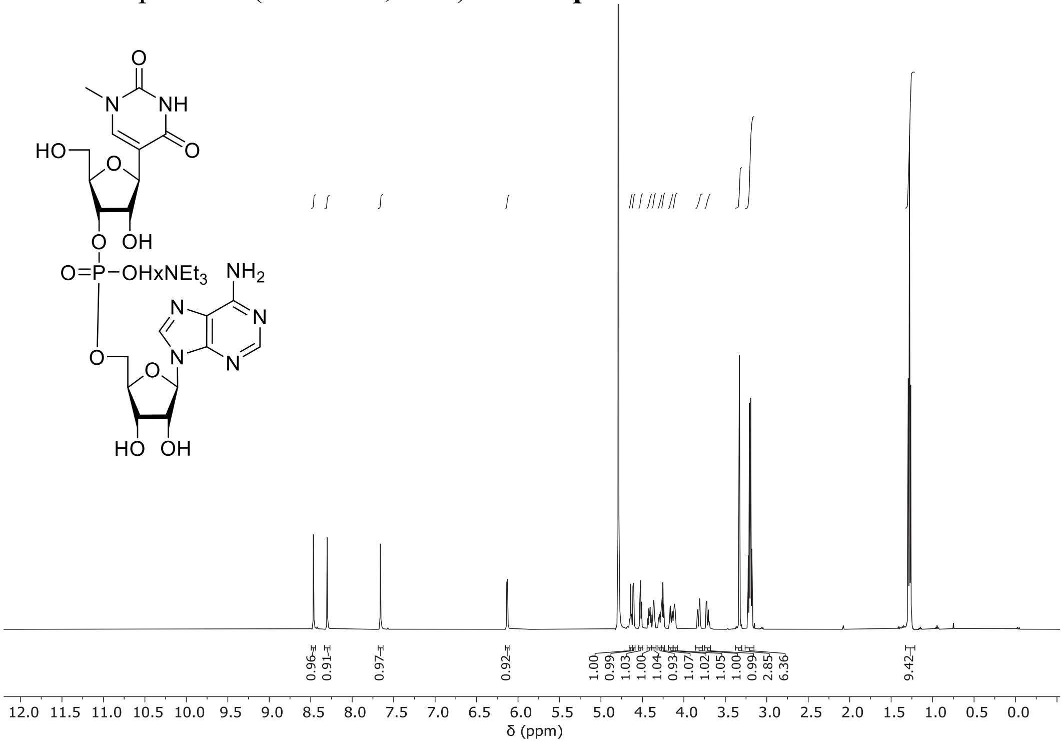

$^{13}$C NMR spectrum (126 MHz, $D_2O$) of **m$^1$ΨpA**·$NEt_3$

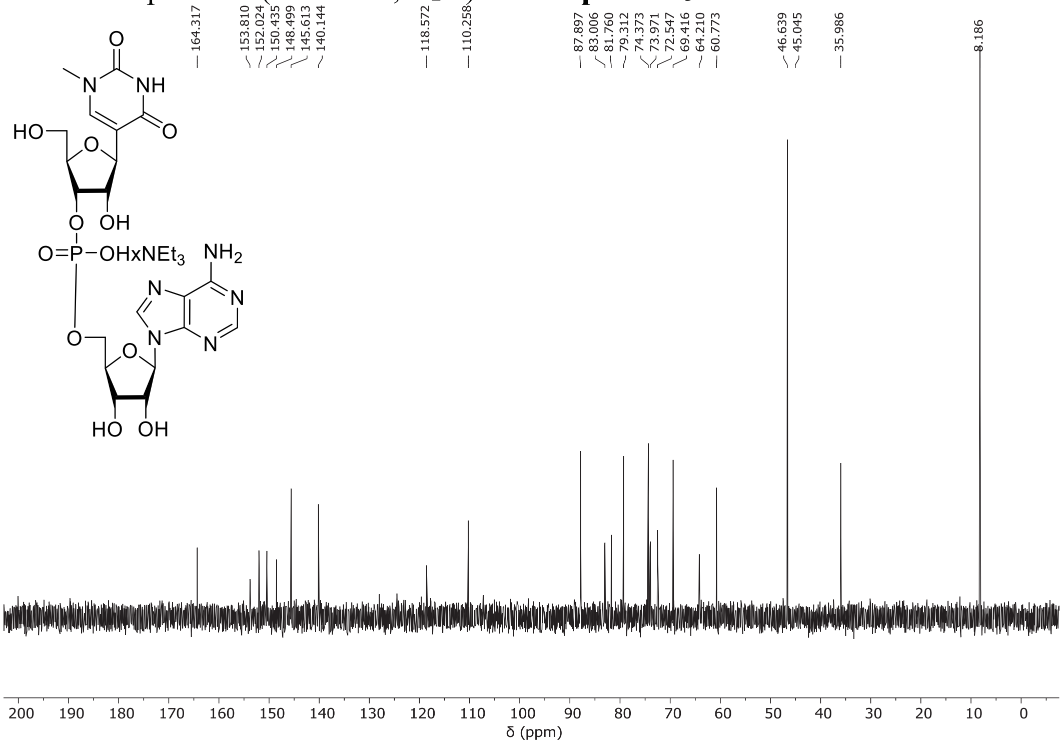

$^{31}$P NMR spectrum (202 MHz, $D_2O$) of **m¹ΨpA**·$NEt_3$

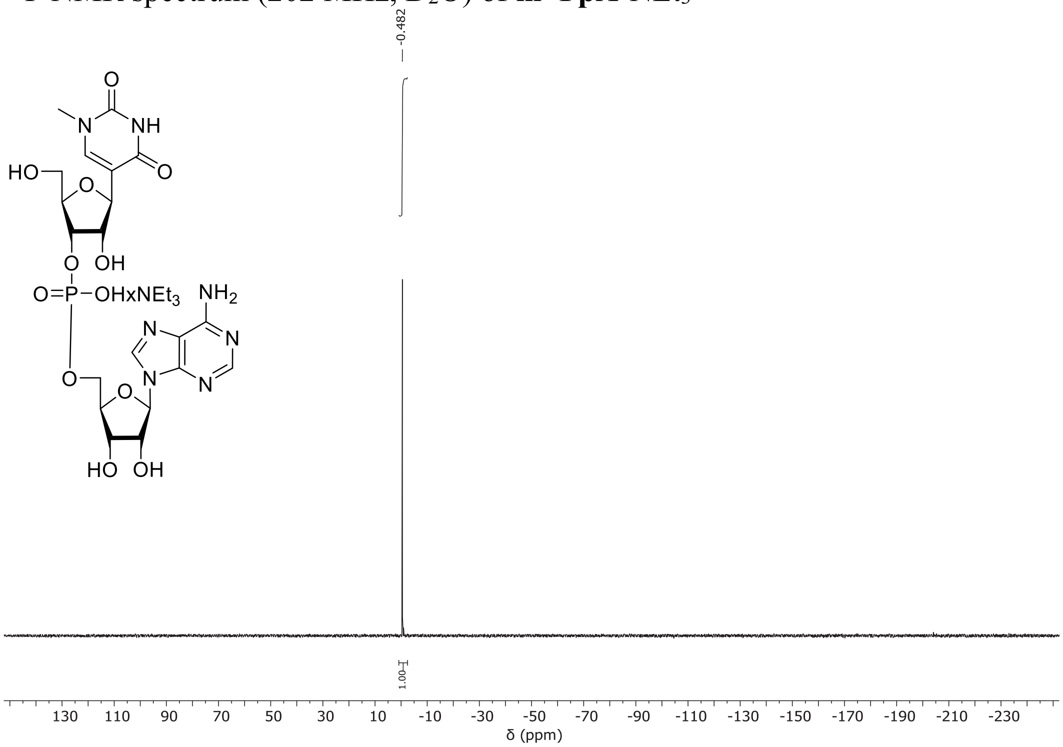
